# Model of a striatal circuit exploring biological mechanisms underlying decision-making during normal and disordered states

**DOI:** 10.1101/2024.07.29.605535

**Authors:** Dirk W. Beck, Cory N. Heaton, Luis D. Davila, Lara I. Rakocevic, Sabrina M. Drammis, Danil Tyulmankov, Paulina Vara, Atanu Giri, Shreeya Umashankar Beck, Qingyang Zhang, Michael Pokojovy, Kenichiro Negishi, Serina A Batson, Alexis A. Salcido, Neftali F. Reyes, Andrea Y. Macias, Raquel J. Ibanez-Alcala, Safa B. Hossain, Graham L. Waller, Laura E. O’Dell, Travis M. Moschak, Ki A. Goosens, Alexander Friedman

## Abstract

Decision-making requires continuous adaptation to internal and external contexts. Changes in decision-making are reliable transdiagnostic symptoms of neuropsychiatric disorders. We created a computational model demonstrating how the striosome compartment of the striatum constructs a mathematical space for decision-making computations depending on context, and how the matrix compartment defines action value depending on the space. The model explains multiple experimental results and unifies other theories like reward prediction error, roles of the direct versus indirect pathways, and roles of the striosome versus matrix, under one framework. We also found, through new analyses, that striosome and matrix neurons increase their synchrony during difficult tasks, caused by a necessary increase in dimensionality of the space. The model makes testable predictions about individual differences in disorder susceptibility, decision-making symptoms shared among neuropsychiatric disorders, and differences in neuropsychiatric disorder symptom presentation. The model reframes the role of the striosomal circuit in neuroeconomic and disorder-affected decision-making.

**Highlights:**

1. **Striosomes prioritize decision-related data used by matrix to set action values.**
2. **Striosomes and matrix have different roles in the direct and indirect pathways.**
3. **Abnormal information organization/valuation alters disorder presentation.**
4. **Variance in data prioritization may explain individual differences in disorders.**

**eTOC:** Beck et al. developed a computational model of how a striatal circuit functions during decision-making. The model unifies and extends theories about the direct versus indirect pathways. It further suggests how aberrant circuit function underlies decision-making phenomena observed in neuropsychiatric disorders.

## Introduction

Decision-making is an indispensable and ubiquitous process utilized universally by organisms for survival and optimal functioning^1–4^, and is a quantifiable metric for exploring cognition^5^. Critically, abnormal decision-making is recognized as a transdiagnostic symptom^6^ of several neuropsychiatric disorders^7,8^. Numerous brain regions are implicated in decision-making processes^1,9,10^. We modeled a circuit that includes the striatum, Globus Pallidus internus (GPi), Lateral Habenula (LHb), Rostral Medial Tegmental nucleus (RMTg), and dopaminergic neurons of the Substantia Nigra Compacta (daSNC).

Striatal neurons can be neurochemically categorized into groups, including striosomal Spiny Projection Neurons, (striosomal compartment, i.e. sSPNs, ∼10-20% of striatum) and matrix Spiny Projection Neurons (matrix compartment, i.e. mSPNs, ∼80-90% of striatum). sSPNs are implicated in multiple decision-making^11–18^ and motor functions^19,20^. In contrast, the functional role of mSPNs during decision-making remains unclear. Furthermore, striatal neurons are also equally distributed between the direct pathway (direct striosomal Spiny Projection Neurons, dsSPN; and direct matrix Spiny Projection Neurons, dmSPN), identified by D1 receptor expression, and indirect pathway (indirect striosomal Spiny Projection Neurons, isSPN; and indirect matrix Spiny Projection Neurons, imSPNs) identified by D2 receptor expression^21–26^. These pathways play distinct roles in action selection^26–30^, movement^31^, and other decision-making processes^28,32–34^.

sSPNs project to downstream basal ganglia regions including the GPi^35–37^, LHb^38,39^, RMTg, and daSNC^40–43^ subcircuit. This subcircuit has been found to influence behavioral states^44,45^, affect action selection^46–49^, determine subjective valuations^50–52^, respond to aversive/rewarding cues^53–57^ are rewards/costs^58–60^, and encode reward prediction errors (RPE)^61–64^. Neurons in the circuit are differentially affected in neuropsychiatric disorders^7,37,65–76^.

Though extensive research has explored various brain regions and neuronal circuits during abnormal and normal decision-making, there remain numerous questions about the mechanisms that result in different disorder phenotypes^77,78^, individual differences in disorder presentation despite a shared clinical diagnosis^79,80^, individual differences in disorder susceptibility^81–83^, and daily within-person variance in decision-making behavior^84–86^. Additionally, the functional role of sSPN versus mSPN is unclear, as is their interaction with daSNC and associated basal ganglia structures. Also, RPE^61,62,87–89^ has been detected in most of the regions of our modeled circuit, but the interplay of RPE encodings across each of the modeled brain regions remains unclear. Finally, the intersection of sSPNs versus mSPNs and the direct versus indirect pathways of the striatum is incompletely understood. We sought to address these questions by creating a computational model of physiology of sSPNs, mSPNs, and surrounding brain regions during decision-making. Using our model, we propose and test hypotheses regarding how these regions and mechanisms interact during decision-making and how aberrancies in function may give rise to experimentally observed normal and disordered decision-making patterns.

Other models have been developed depicting how a brain region or circuit functions to set value or select an action^90^. A subset of models focusses, as we do, on the role of the basal ganglia. One approach that computational models use to examine basal ganglia function is by exploring how RPE signals facilitate adaptability^91–94^. Other models consider how the direct and indirect pathways of the basal ganglia interact to moderate action selection^95–98^. Others explore the role of basal ganglia pathways in performing dimensionality reduction^99^, aiding in reinforcement learning^98,100–102^, or responding to events as sequences that influence each successive action^103–105^. However, there remains a crucial need for a model which defines the functional role of sSPNs, and their interactions with neighboring brain regions, based on recent experimental literature.

We developed a computational model of the physiological mechanisms behind choice (**Figure 1**). We tested the model using selected experimental studies for each structure as well as analysis of neural recordings and decisions in models of healthy and disordered behavior (**Figures 2,3**). We demonstrate its alignment with the experimental literature on prediction error and direct and indirect pathways (**Figures 4**). Finally, we outline hypotheses of how different patterns of circuit activity cause aberrant decision-making in disorders and altered disorder progression (**Figures 5,6**,**7**). Overall, our model provides a unifying framework for this circuit’s physiological function during decision-making and motivates testable hypotheses about the link between the circuit and choice.

**Figure 1.**
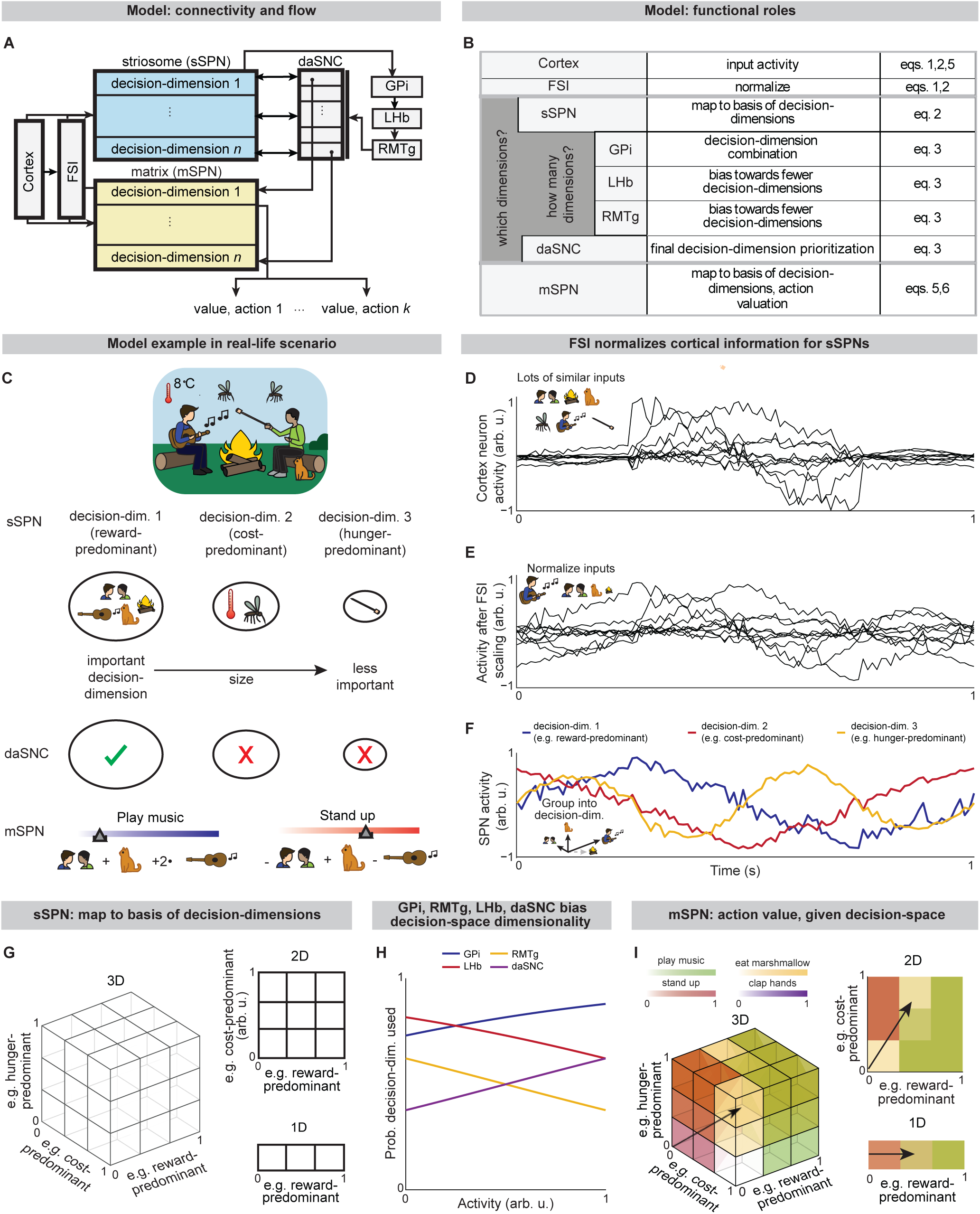
The circuit defines a decision-space for action valuation. **A-C.** Cortex encodes environmental and internal information. Striatal FSIs normalize activity from cortex. sSPN subpopulations each encode data along a decision-dimension. GPi→LHb→RMTg re-weight the importance of using each decision-dimension. daSNC subpopulations, each corresponding to a decision-dimension, activate when their decision-dimension is important. This causes dopamine to be released to select mSPN subpopulations, and thus, a “decision-space” is formed from the basis of the important decision-dimensions. mSPNs define action values within decision-space. **A.** Circuit architecture. **B.** Functional role of circuit elements. **C.** Cortex encodes signals about food, social, and environmental cues (cartoon). sSPNs project cortical information to decision-dimensions. Here, a “reward-predominant” decision-dimension captures information about music, cat, fire, and social interaction; a “cost-predominant” decision-dimension captures information about temperature and mosquitoes, and a “hunger-predominant” decision-dimension captures information about marshmallows. The most important decision-dimensions (here, the reward-predominant decision-dimension only, assigned the checkmark) are retained. mSPN forms action values using rules corresponding to the retained decision-dimensions. Several actions (here, “play music” and “stand up”) are assigned values, and then decisions are made based on the values of possible actions. “dim.” = dimension. **D-F**. Example showing how FSIs and SPNs parse cortical activity. The signals of 10 cortex neurons encoding sensory information (top) are normalized by FSI such that activities are on a more uniform scale (middle) and then mapped to decision-dimensions in sSPN (bottom). **G**. During decision-making, sSPNs help to determine which decision-dimensions are most important. Important decision-dimensions are later formed into decision-space (grids of cubes or squares) when dopamine is released from daSNC to mSPNs. For example, a 1D decision-space might be constructed where decision-making is conducted using information predominantly about reward, or multi-dimensional decision-space might be constructed for situations where multiple decision-dimensions are important to the decision. **H**. GPi, LHb, RMTg, and daSNC activities affect the probability a decision-dimension will be used to form the decision-space. The activities of the brain regions are incremented and daSNC activity (equivalent to probability the decision-dimension is used) is plotted as the responding variable. **I.** After the decision-space is formed, mSPNs define action values (colors) based on cortical input (vectors) and the decision-space (grids). In a 1D decision-space, action values are assigned purely based on the activity of mSPN along a single decision-dimension. In multi-dimensional spaces, action values depend on the activity along multiple decision-dimensions.

**Figure 2.**
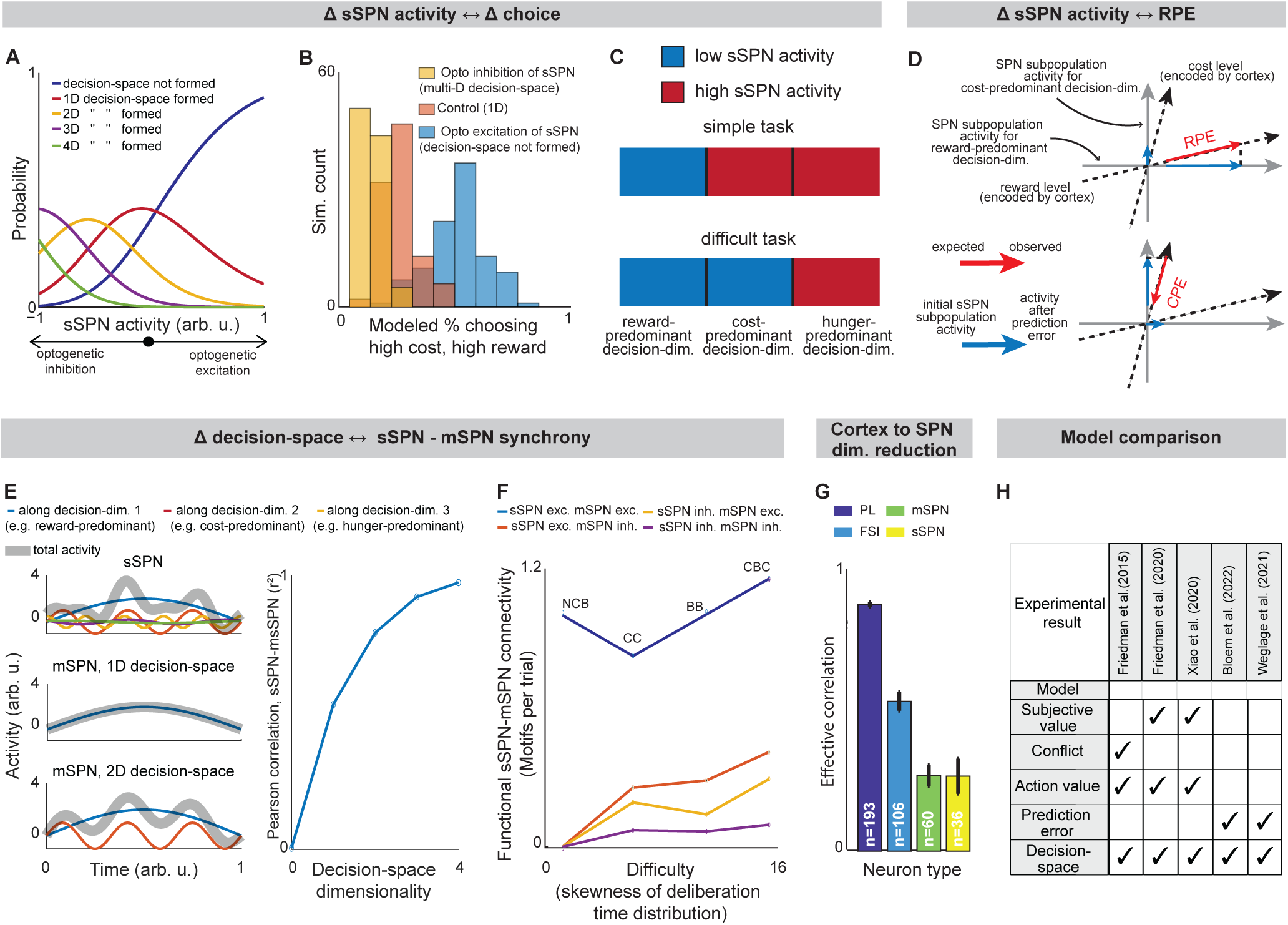
Evidence of decision-space formation in sSPNs and mSPNs. **A.** Changes to sSPN activity, for instance during optogenetic manipulation, cause changes in the number of decision-dimensions used to form decision-space. *b*_sSPN_ is incremented in eq. (1) from -1 (low activity) to 1 (high activity). **B.** More consistent choices are made at lower sSPN activity. 100 choices are simulated for each of the three decision-space scenarios. Modeled choices are made between a high-cost, high-reward option and a low-cost, low-reward option. The multi-D decision-space applies rules for both reward and cost, the 1D decision-space rules about only reward, and the “decision-space not formed” case applies neither. Then action values and choice are derived. **C.** An sSPN subpopulation has reduced activity when it is used to form decision-space. Therefore, mean sSPN activity is different in simple tasks requiring low-dimensional decision-space versus difficult tasks requiring high-dimensional decision-space. **D.** In some cases, sSPNs and mSPNs will shift their activities proportionally to a prediction error. Cortical neurons continuously encode mixed information about reward, costs, and other features across many inputs (dashed lines show 2 inputs, e.g. chocolate milk “reward” and flashlight “cost”). When cortical information about a certain feature (e.g. level of chocolate milk) suddenly changes, this change is mapped to all decision-dimensions. However, certain decision-dimensions will have more alignment to data axes like reward and cost (gray lines). Thus, when sSPNs change their activities to account for new cortical information, reward or cost prediction errors may be revealed by the activities of certain (and mostly separate) SPN subpopulations. RPE = reward prediction error, CPE = cost prediction error. **E.** When one-dimensional decision-space is formed, sSPN (top-left panel) and mSPN (middle-left) activities have low correlation over time. As more decision-dimensions are used to form decision-space, sSPN and mSPN activities increasingly correlate (bottom-left). Pearson’s correlation between sSPN and mSPN activities (right panel). **F.** Experimental data showing that task difficulty (measured through skewness of the deliberation time distribution) increases with functional sSPN-mSPN connectivity (measured through Granger causality) during a decision. Tasks: NCB = non-conflict cost-benefit (sSPNs = 14, mSPNs = 260), CC = cost-cost (sSPNs =46 , mSPNs = 400), BB = benefit-benefit (sSPNs =83 , mSPNs = 1246), CBC = cost-benefit conflict (sSPNs = 84, mSPNs =717). The CBC task has significantly more motif counts per trial (p<0.003, compared to shuffled data). **G.** Experimental data showing that simultaneously recorded SPNs have less correlation (measured as effective correlation) than FSIs or prelimbic cortical neurons. This indicates dimensionality reduction from cortex to SPNs. Significances of difference from cortex: FSI p<10^-18^, mSPN p<10^-45^, sSPN p<10^-63^. **H.** Success of the decision-space and our alternative models in explaining experimental sSPN activity during decision-making.

**Figure 3.**
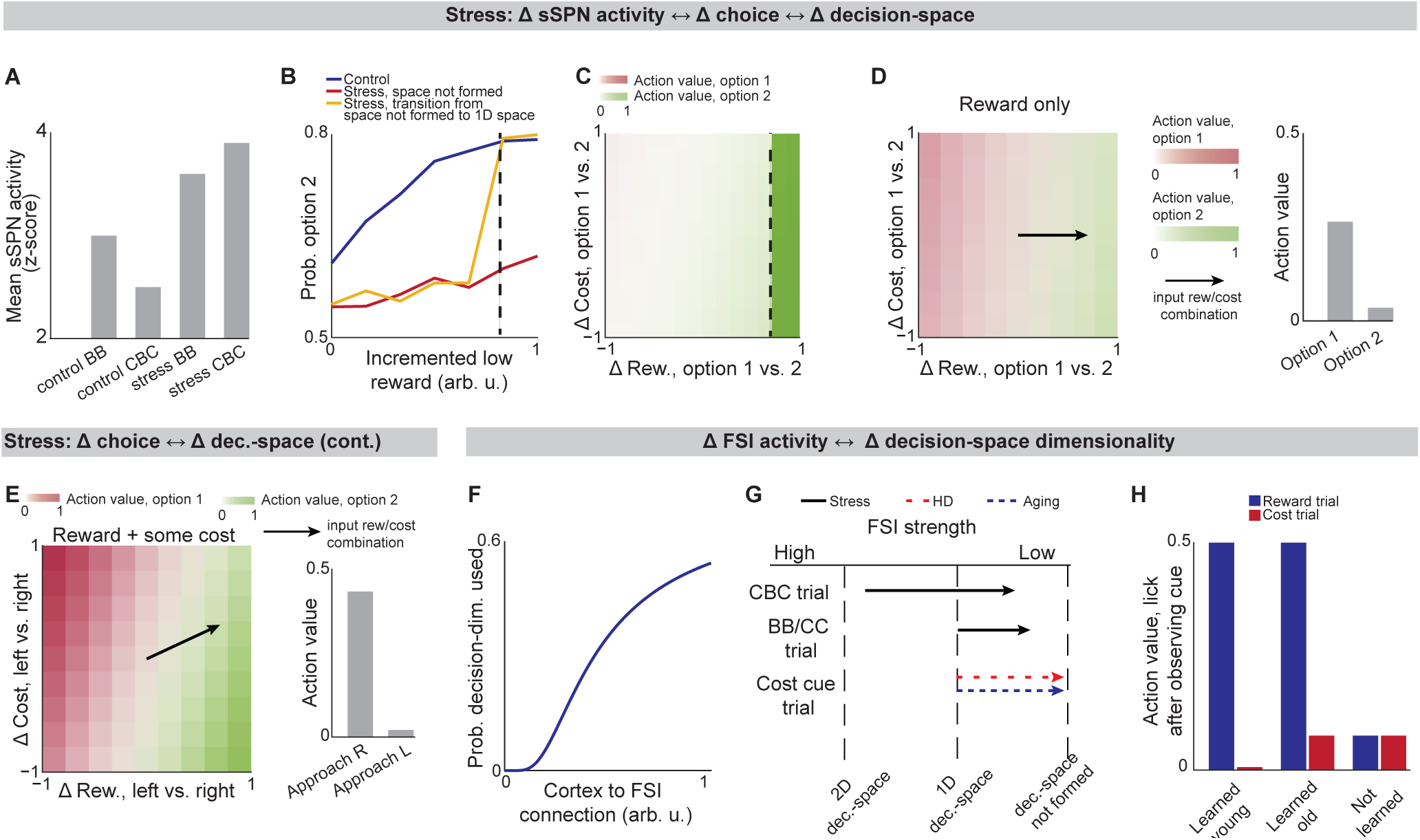
Lower-dimensional decision-spaces are produced in stress, aging, and HD, affecting decisions. Data was taken from experiments where rodents performed the cost-benefit conflict (CBC) task, in which rodents had to select between a high cost-high reward option and a lower cost-lower reward option. Two behavioral control tasks were also used: the benefit-benefit (BB) task, in which rodents selected between a high reward option and a low reward option (equal and minimal cost for both), and the cost-cost (CC) task, in which rodents selected between a high cost option and a low cost option (equal and minimal reward for both). **A**. Summary of the experimental finding that sSPN activity is increased after stress, especially during the more difficult CBC task. **B,C**. Modeled psychometric functions (**B**) and action values across experimental conditions (**C**) for a rodent in a T-maze task after chronic stress. The circuit forms a 1D decision-space only after reward exceeds a critical concentration (dashed line). At this point, the stress-group rodents switch from choosing the options roughly evenly to most often turning right towards the lower-cost option. Psychometric functions resemble experimental decision-making data. **D,E**. After stress, a low-dimensional decision-space may be used by default (**D**) but a higher-dimensional decision-space may sometimes form during difficult tasks, for example those with both reward and cost (**E**). This leads to the counterintuitive result that adding cost to an offer (vectors on colormaps) can increase its action value (bar plots). **F.** Cortex→FSI connection strength affects decision-space dimensionality by altering the propensity of a decision-dimension to form decision-space. **G**. Cortex→FSI connection strength is reduced in stress, Huntington’s disease, and aging. This leads to lower-dimensional decision-space. **H**. Model of an operant conditioning task in Friedman et al (2020). Modeled action values for licking versus not licking in an operant conditioning task. There are two tasks: 1) responding to a reward cue by forming decision-space from a reward-predominant decision-dimension, and 2) likewise for cost. “Learned young” succeeds at 1 and 2, “learned old” at 1 but not 2, and “Huntington’s disease” at neither. Resembles experimental licking rates.

**Figure 4.**
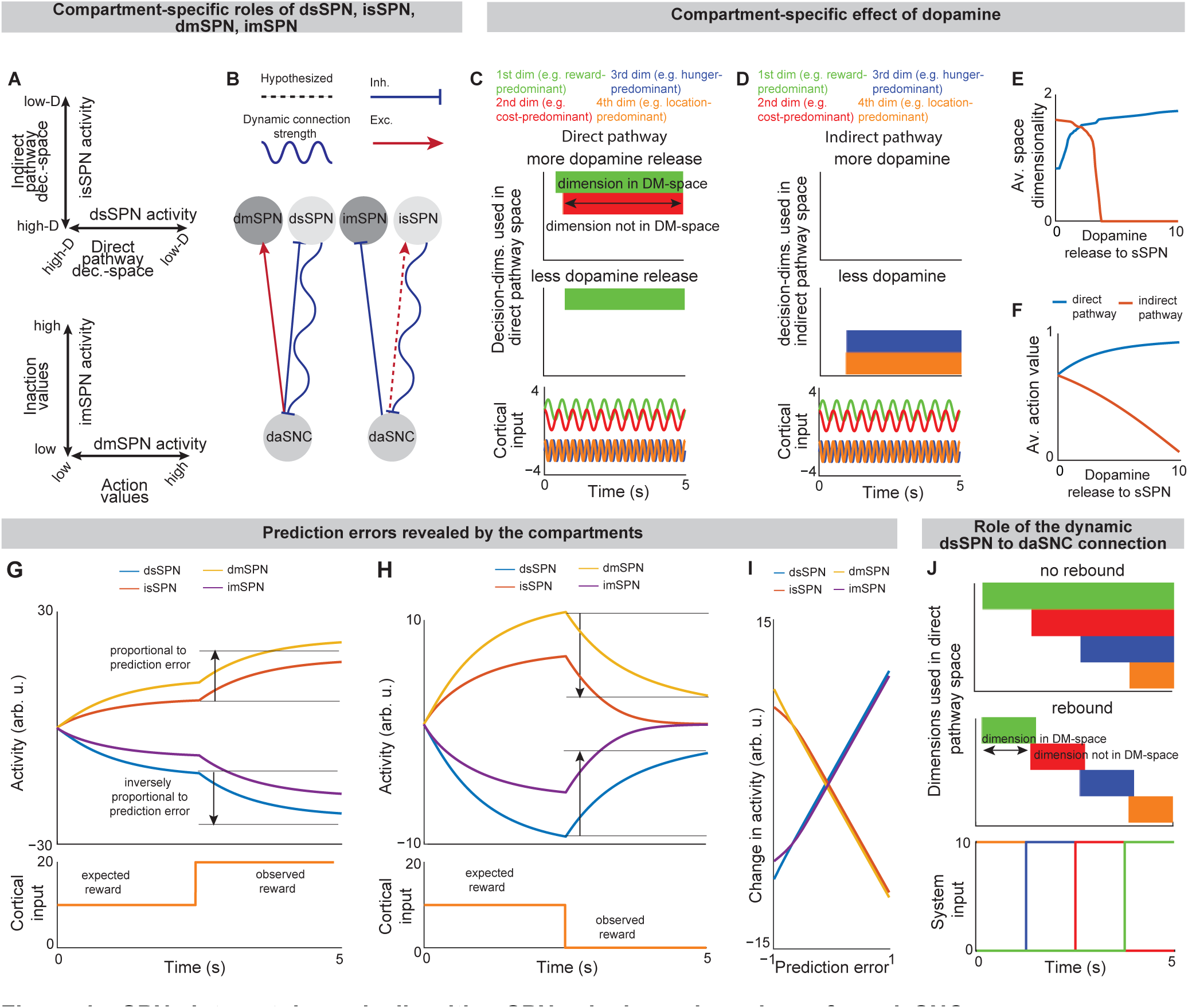
sSPNs interact dynamically with mSPNs via dopamine release from daSNC. **A.** Direct pathway and indirect pathway decision-spaces are constructed in parallel by sSPNs. mSPNs uses those decision-spaces to define action values (direct pathway) and inaction values (indirect pathway). **B.** Dynamic dSPN, iSPN, sSPN, mSPN and daSNC interaction. sSPN inhibits daSNC. Dopamine lengthens upstates in dmSPN and shortens them in imSPN. The connection from sSPN to daSNC adjusts over time based on sSPN activity (spring). **C-E.** Simulations showing dimensions used to form direct pathway decision-space over time in response to a cortical input (bottom panel) carrying information along several example decision-dimensions. Responses are shown for the direct (**C**) and indirect (**D)** pathways at two levels of dopamine release to sSPN or mSPN and across incremented levels of dopamine release (**E**). Increased dopamine release leads to higher-dimensional decision-space formed in the direct pathway and lower-dimensional decision-space in the indirect pathway. **F.** Dopamine release to dmSPN tends to increase action values and dopamine release to imSPN tends to decrease action values. **G,H.** sSPN signaling of positive (**G**) or negative (**H**) prediction error. **I.** The simulations in **G**,**H** are run for incremented differences in expected versus observed reward, from -1 (negative prediction error) to +1 (positive prediction error), demonstrating that each population changes its activity roughly proportionally to prediction error. **J.** 1.25s pulses of input from cortex (bottom panel) are applied along the four decision-dimensions in succession. Connection from sSPN to daSNC facilitates the rapid de-prioritization of decision-dimensions no longer required in decision-space in the daSNC “rebound” scenario (middle panel). This is because the longer sSPN inhibits daSNC, the more the connection weight decreases in strength, leading to daSNC receiving decreasing signal about the importance of the decision-dimension. Then, when sSPN ultimately signals that the decision-dimension is no longer important, this signal is enhanced upon reception by daSNC. In the “no rebound” scenario causes decision-dimensions to continue to be prioritized for the duration of the simulation (top panel).

**Figure 5.**
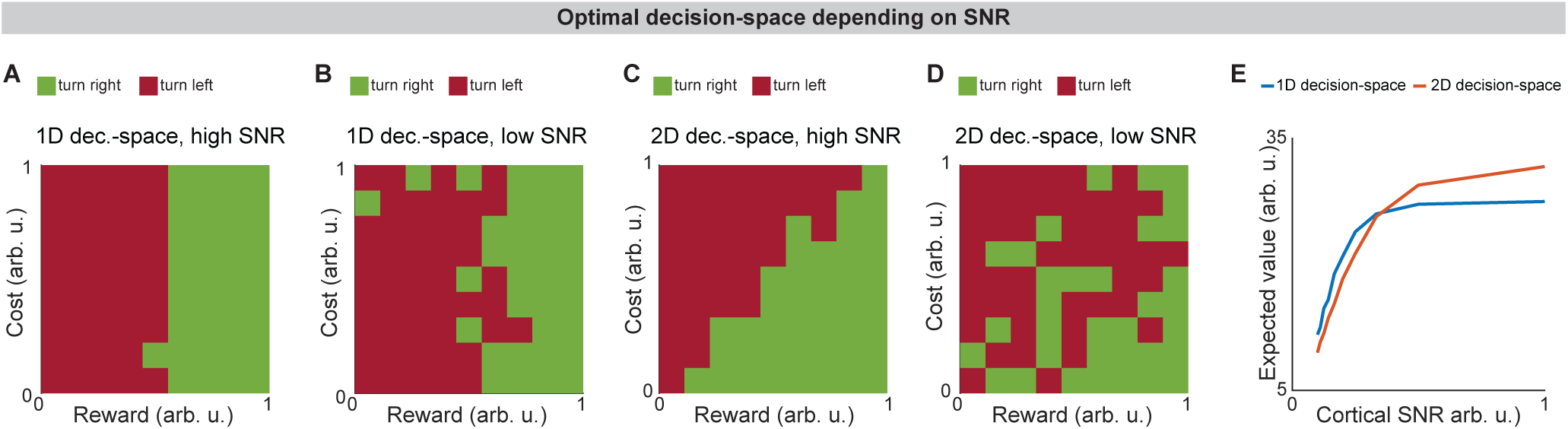
Decision-space is differentially constructed based on cortical signal-to-noise ratio (SNR). **A-D.** Modeled T-maze task where an animal turns right to choose a reward/cost offer or turns left to avoid it. Cases where there is high cortical SNR (**A**,**C**) or low cortical SNR (**B,D**) and a 1D decision-space (**A,B**) or 2D decision-space (**C,D**) are formed. Modeled “turn right” actions are considered successful (positive value) when reward > cost, and “turn left” when reward < cost. 2D decision-spaces lead to more value when there is low cortical SNR (**C**) but not when there is high SNR (**D**). **E.** Choices using different types of decision-spaces have different expected reward minus cost (expected value), depending on cortical SNR.

**Figure 6.**
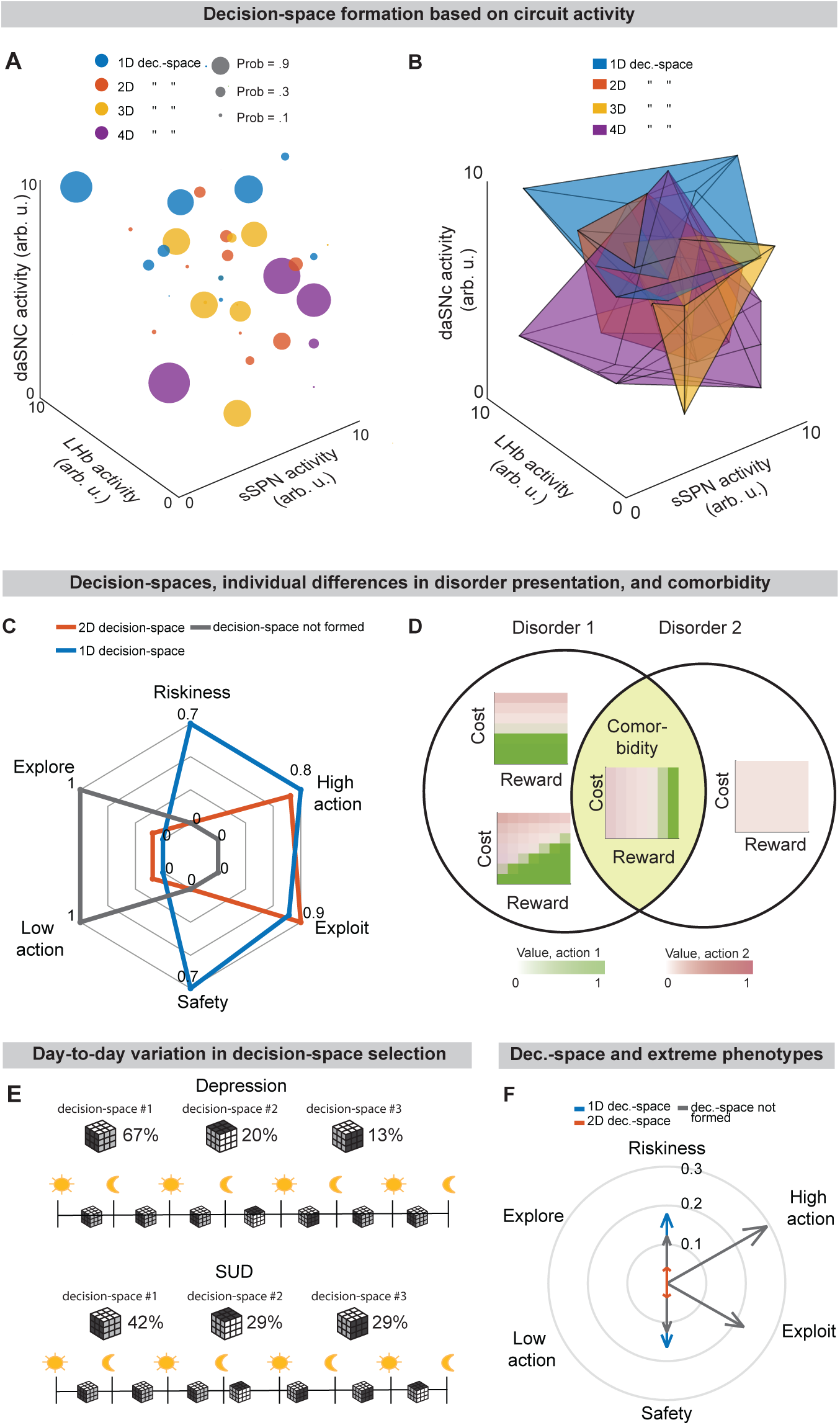
Differences in decision-space could explain comorbidity, individual differences, and daily variations. **A**. 100 simulations are run at arbitrary circuit activities and points where the decision-space formed are retained, illustrating the overall relationship between circuit activity (which is different across disorders) and decision-space, although multiple types of decision-space can form at the same circuit activity. **B**. Similar to **A** but with the points connected by faces, illustrating the possibility of formation of multiple decision-spaces at the same circuit activities. **C**. Summaries of overall decision-making profiles across trials in a T-maze task, scored in six ways, showing that transitions between decision-spaces can lead to very different decisions. A disorder could produce a bias, for instance, towards low-dimensional decision-spaces, which would in turn alter the decision-making profile. Scores are formed by quantifying the trend of action values across reward and cost levels. Riskiness/safety: treatment of high-reward, high-cost (or low-cost, low-reward) levels; high/low action: tendency towards high (or low) action values; exploit/explore: tendency to focus on one action versus many (**see Figure S6C**). **D,E**. Differences in circuit activity between individuals could lead to decision strategies observed at different rates, as is the case in individuals with disorder comorbidity **(D**). Further, day-to-day shifts in circuit activity shifts could cause stark differences in decision strategies between days (**E**). **F**. Certain decision-spaces more often lead to action values that are extreme (ratio formed over 1000 simulations, as scored using the metrics in **C**), a feature of disorders. Vector length corresponds to outlier rate (proportion of scores for each group that fall within the top 10% of observations across all groups).

## Results

### Description of the model

Our model demonstrates how a neural circuit (cortex, dsSPNs, isSPNs, dmSPNs, imSPNs, GPi, LHb, RMTg, and daSNC) defines action values during decision-making (**Figures 1A-C**, S1A, Analyzed Instances of the Model, STAR Methods).

Cortical input to striatum consists of millions of neurons that encode rewards/costs^58,59^, decision-making schemas^106^, decision-making states^107,108^, value^109–111^, and environmental information^112,113^ that synapse onto several hundred thousand SPNs^114–116^, indicative of dimensionality reduction. Within the dorsomedial striatum, Fast Spiking Interneurons (FSIs) receive convergent cortical signals and suppress SPNs^117^, similar to the operation they perform in other brain regions^118–120^. We model this suppression as normalization of cortical activity (**Figures 1D,E**) where cortical signals of different magnitudes are rescaled, allowing striatal neurons to respond to cortical activity appropriately. See Defining FSI activity, STAR Methods.

Cortical neurons synapse onto subpopulations of proximate sSPNs^121,122^ which are critical for conflict decision-making^11,123^, value/stimulus-based learning^12,124^, responding to negative stimuli^14^, responding to probabilistic cues/RPEs^15,16^, and switching strategies as contexts change^13^. We postulate that sSPNs in different subpopulations respond differently to cortical input. Within each subpopulation, sSPNs encode inputs along a data axis we term a “decision-dimension,” modeled as a principal component^125^ of cortical activity, such as could be learned via Hebbian plasticity rules^126,127^. For simplicity, we can conceptualize decision-dimensions as encoding abstract variables like reward, cost, or hunger (**Figures 1F,G**), although in practice they may have less interpretable semantic labels and will each map onto some combination of the variables encoded by the cortex. Mathematically, we define a matrix **W**_*P*_ for each pathway *P* (direct or indirect), the columns of which define the decision-dimensions. This matrix maps cortical activity x_*P*_ to the coordinate space of sSPNs, where it undergoes divisive normalization by FSI activity *c_P_* and a shift in activity **b**_sSPN_ to form sSPN activity s_sSPN,*P*_:

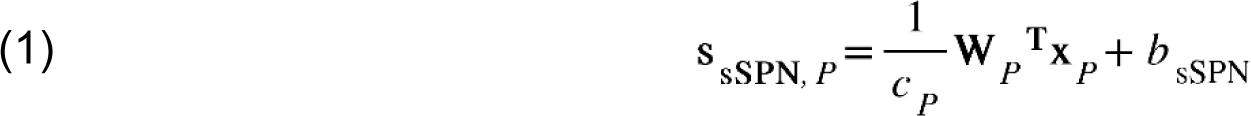

Signals are passed from sSPN subpopulations to daSNC in two ways: 1) sSPN→daSNC^21,41,42,128,129^; and 2) sSPN→GPi→LHb→RMTg→daSNC^38,130,131^. The GPi encodes reward^60,132^, motivation^36^, and facilitates behavioral adaptation. The LHb has been found to be important for uncertainty/aversion^73,133^, behavioral flexibility^134^, and encodes RPEs^61^. The RMTg is important for punishment learning and may be sensitive to the correlation between a cue and the associated outcome^135^. RMTg projects to multiple daSNC subpopulations^136^, which each encode distinct information^137^ and release dopamine to the striatum^138^. Dopamine is important for various aspects of decision-making such as influencing action selection^139^, RPE^62,140^, effort^141^, and the encoding of value^142,143^. GPi integrates signals from many sSPN subpopulations through synapses that release both GABA and glutamate^144^. The LHb, when activated by GPi^60,132^, inhibits daSNC through RMTg^133,145^, and the RMTg inhibits daSNC activity^146^. In our model, these circuit elements serve the functional role determining the priority of using each decision-dimension during the decision (**Figure 1H**, S1B-I). We represent this as daSNC combining activity from sSPNs **s**_sSPN,*P*_ (with connection weights *w*_sSPN→daSNC,*i,P*_ corresponding to each decision-dimension *i*) with RMTg activity and an additive shift *z*_daSNC,*i,P*_:

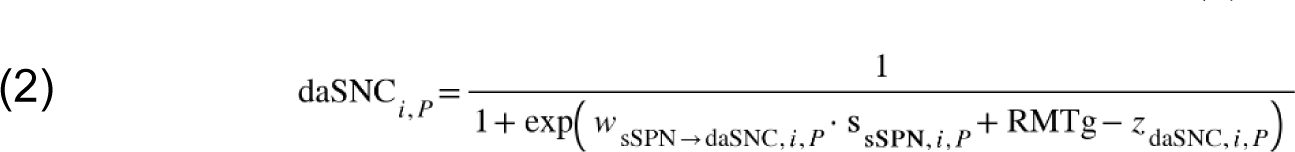

When a daSNC subpopulation is active, dopamine is released to a corresponding subpopulation of mSPNs, termed a matrisome^44,147^, resulting in enhanced or inhibited mSPN reception of cortical signal^37,44^. Thus, dopamine converts the priority of a decision-dimension (as encoded by daSNC) to a weight indicating whether cortical activity along the dimension is, in fact, processed by mSPNs. In this way, a “decision-space” is formed among mSPNs. We use the convention that the decision-space is constructed from only those decision-dimensions deemed important. So, for instance, a high-dimensional decision-space would be formed when many decision-dimensions are important to the decision-making process. Mathematically, mSPN activity is defined similarly to sSPN activity, but for a diagonal matrix corresponding to the dopamine release that probabilistically defines the decision-space, with 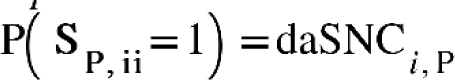:

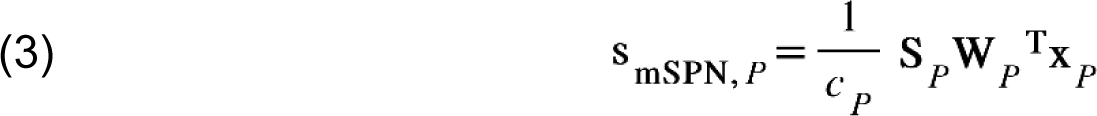

mSPNs collectively define action values (the value of performing various actions, encoded by the direct pathway) and inaction values (the value of refraining from those actions, encoded by the indirect pathway). (**Figures 1I** and S1J-L). Values (for each action/inaction *v_j,P_*, pathway *P*) are defined based on mSPN activity, internal coefficients β_*j,P*_, and priors α*_j,P_*:

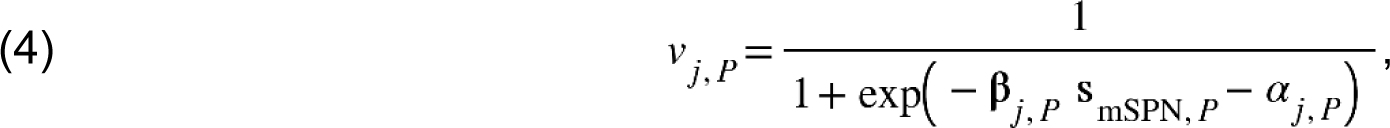

Based on these values, actions are either performed or refrained from over time (Figures S1M-P). We model this using a Merton process model where the first process to hit a threshold is enacted (direct pathway) or refrained from (indirect pathway). See Defining choice, STAR Methods.

In summary, our decision-space model has three requisite hypotheses: 1) sSPNs encode representations of cortical activity in sSPN subpopulations (corresponding to decision-dimensions), 2) dopamine is released by daSNC subpopulations to corresponding mSPN subpopulations (forming the decision-space), and 3) action values are defined by the activity of mSPN subpopulations after the decision-space is formed. Based on these hypotheses, we model how circuit elements physiologically interact during decision-making. This produces a convenient geometric interpretation (“decision-dimension”, “decision-space”) and an intuitive description of the functional processes of dimensionality reduction and action valuation. It is also convenient for connecting circuit activity to observed patterns in choice. For instance, we can form a heuristic linking the dimensionality of the decision-space with choice: we might expect low-dimensional decision-spaces to be used during simple choices (e.g. between two rewards) and high-dimensional decision-spaces to be used during more difficult choices (e.g. between offers which each have benefits and costs). Using this heuristic, we can infer the decision-spaces of behaving rodents^148^ and humans^149^ by regressing sSPN activity on experimental parameters (for example, temperature or music volume), as we demonstrate using synthetic data (Figures S1Q,R, Inferring decision-space from SPN activity and choice, STAR Methods).

Next, we sought to define tests where the model might be supported or disproven by experimental data collected during decision-making. To do this, we considered predictions which set the model apart from others of basal ganglia function and which could be convincingly tested using tools available to today’s neuroscientists.

### Model validation

#### Choices made at different levels of sSPN activity

Changes to the activity of sSPNs, via their inhibition of daSNC neurons, should affect the priority assigned by dopamine to each decision-dimension. Thus, we would expect sSPN activity during a decision to influence the type of decision-spaces that are formed. This appears to be the case. Experimentally, optogenetic stimulation or inhibition of sSPNs during a rodent conflict decision-making task alters choice selection^11^ in a way indicative of changes to the decision-space (**Figures 2A,B**, **S2A, Effect of sSPN activity on decision-space** and **Effect of decision-space on choice, STAR Methods**). The inverse is also true: difficult choices, which the model predicts to be made using a high-dimensional decision-space, are made with different levels of sSPN activity both experimentally^11^ and in our model (**Figures 2C**, **S2B-D**).

#### Response of SPNs to rapid changes in cortical information, as occurs during reward prediction errors

SPN subpopulations encode data along decision-dimensions, so we should expect them to respond similarly to related stimuli, even if the stimuli arrive from different cortical neurons. An example of this might be two stimuli related to reward: the sound of chocolate milk poured in a glass, and the taste of chocolate milk. In reward prediction error theory, these two stimuli would be linked by a value function, where expected reward rises when the chocolate milk is poured, and then rises again if the milk is tasted, or falls if the milk is withheld^89,150^. In the decision-space model, a similar phenomenon should occur, but with a different interpretation. Information about reward, for instance, is unlikely to completely align with a decision-dimension but should align to some decision-dimensions more than others (**Figure 2D**). Thus, when in the example milk is poured, an sSPN subpopulation corresponding to reward-predominant information is innervated, and the same subpopulation further adjusts its activity when the taste of chocolate milk arrives.

In cases where a cue is tied to a reward or cost response and a prediction error is expected, a range of experimental studies show that some sSPNs shift their activities^15,16,62^. The decision-space model also expects separate SPNs to encode data along different decision-dimensions. Indeed, sSPN subpopulations have been found that track either reward prediction errors or cost prediction errors, but not both^14,15^.

The decision-space model also makes two hypotheses contrary to most prediction error theories. First, it is expected that in a Pavlovian task, the sSPNs of even fully conditioned animals should respond to an outcome as well as the cue for that outcome. This hypothesis has experimental support^14^. Additionally, the model makes the prediction, currently untested, that information along axes other than reward and cost will have mixed encoding in cortex but separate encoding in SPNs. For example, the physical proximity of two separate objects may be encoded by a single SPN. If so, moving the first object closer to the subject should lead to excitation of the SPN, and later moving the second object further away should lead to inhibition of the same neuron.

#### sSPN-mSPN synchrony depending on the decision-space

Because the decision-space is formed by the interaction of sSPN subpopulations with associated matrisomes via daSNC, the activities of sSPNs and mSPNs should be most synchronized when more subpopulations are used for the decision, that is, when a high-dimensional decision-space is formed (**Figure 2E**). So, we would expect high synchrony during difficult decisions. To test this, we analyzed SPN data from the Corticostriosomal Circuit Stress Experimental database^123^ using cross-correlation and a custom Granger causality-based tool. According to both analyses, functional connectivity scaled with task difficulty, as scored based on deliberation time (**Figures 2F**, **S2E-N, Connected SPNs through cross-correlation** and **Connected SPNs through Granger causality, STAR Methods**).

#### Dimensionality reduction from cortex to SPNs

We also found direct evidence of dimensionality reduction from cortex to SPNs. Cortical neurons have the most coordinated activities over time (measured as effective correlation^151^), then FSIs, followed by sSPNs and mSPNs **(Figure 2G**, **Analyzing neural dimensionality reduction, STAR Methods)**. This suggests that information is condensed from the cortex, where many neurons encode similar information, to SPNs, which encode mostly different information, as is the case in the decision-space model.

#### Alignment to other experimental work on SPNs

To gauge alignment of the decision-space with the experimental literature, we devised five tests that link sSPN activity to choice, each verifiable with the experimental literature, that might support or reject the decision-space model (**Figures S2O-X, Table S2**). We also constructed, from roles commonly assigned to sSPNs, four alternative models in which sSPNs encode 1) conflict, 2) subjective value, 3) prediction error, or 4) actions. We found that while the alternative models each can be used to interpret a subset of the experimental evidence, the decision-space model aligns to the breadth of the experimental evidence (**Figure 2H**, **Table S3**). A selection of experimental studies on GPi, LHb, and daSNC^60,61,72,132,134,135^ also align with the decision-space model (see **Tables S4-6**). Thus, the model offers a new lens through which to interpret their functions during decision-making.

#### Inability to form a high-dimensional decision-space in disorders

In an experimental study on chronic stress^123^, sSPNs were hyperactive during the cost-benefit conflict task in which they had to choose between two cost-reward options (**Figure 3A**). Meanwhile, the rodents were less adherent to reward level, a feature of choices made using a lower-dimensional decision-space (**Figures 3B,C**, **Effect of decision-space on choice, STAR Methods**). Thus, we wondered if post-stress sSPN hyperactivity causes altered choices due to differences in formation of the decision-space.

We found evidence to support this. Animals made decisions in the cost-benefit conflict task more quickly after stress, as if the task were less difficult (**Figures S3A-D, Defining decision difficulty by task, STAR Methods**). Meanwhile, after stress, choices involving both reward and cost no longer had more functionally connected sSPNs and mSPNs than the simple tasks. In fact, functional connectivity was similar across tasks and to the simple tasks for controls (**Figures S3E-I**). Thus, after stress, the rodents both showed both neural signatures and choices aligned with a low-dimensional decision-space.

The inability to form a high-dimensional decision-space can also explain a counterintuitive finding that stress causes rodents to prefer a reward-cost combination over a reward presented without cost (**Figure S3J, Changes to choice after adding cost to a reward offer, STAR Methods**). In classic economic theory, the addition of cost to a reward typically makes the reward less attractive^152,153^ and thus our observation cannot be readily explained. In contrast, the decision-space model offers a simple explanation: cost can increase offer attractiveness in instances where cost level causes a transition from a default low-dimensional decision-space to a higher-dimensional decision-space (**Figures 3D,E**), as we hypothesized is the case after stress in **Figures 3B,C**. In other cases, decisions are predicted to resemble those predicted by classic theory, for example in cases where either a one-dimensional (as in **Figure 3D**) or two-dimensional (**Figure 3E**) decision-space is used across cost levels.

Cortex→FSI connectivity is impacted by chronic stress^123^ (**Figures S3K-R, Analyzed cortex-FSI connectivity** and **Modeled cortex-FSI connectivity after stress, STAR Methods**), leading to hyperactive sSPNs. Our model suggests that this causes the formation of a lower-dimensional decision-space (**Figures 3F,G**), reflected by changes in the variance and mean of the SPN population (**Figure S4A-G**). Interestingly, the cortex→FSI connection is also impacted in Huntington’s disease and aged rodents^12^, raising the possibility that an inability to form a high-dimensional decision-space during difficult decisions is a feature of multiple health conditions. Supporting this hypothesis, a model where Huntington’s disease and aged subjects use a lower-dimensional decision-space produces action values that follow the trend of experimental choice^12^ (**Figure 3H**, **Effect of decision-space on choice, STAR Methods**).

#### Dopamine release to direct and indirect pathway sSPNs and mSPNs alters choice

sSPNs and mSPNs have been found to be divided approximately evenly between the direct and indirect pathways of the striatum^21,22^, with dopamine release differently affecting sSPNs versus mSPNs and also dSPNs versus iSPNs^26–28,154–156^. Various neuropsychiatric disorders are associated with disturbed sSPN versus mSPN^37,75,157^ and direct versus indirect pathway^158,159^ balances. We sought to explore, through the lens of the model, the functional roles of the different neural responses to dopamine and determine whether these functions might shed light on the experimentally established effects of dopamine on decision-making.

In the model, two decision-spaces are formed in parallel, one for determining whether to perform an action (direct pathway) and the other for determining whether to refrain from it (indirect pathway) (**Figure 4A,B**). Dopamine release leads simultaneously to a higher-dimensional direct pathway decision-space but a lower-dimensional indirect pathway decision-space (**Figure 4C-F**, **Modeling time-variant input, STAR Methods**). Thus, the circuit has a natural mechanism to separately analyze the plusses and minuses of a choice using different data, adding nuance to the decision-making process.

This balance between two decision-spaces offers an intuitive explanation for a range of experimental observations on the direct versus indirect pathway. For example, our model replicates the experimental observations that increased dopamine leads to riskier^160^ and quicker^161^ decisions and preference for nearby offers, in time or physical proximity, compared to distant offers^162,163^ (**Figures S5A-I, Effect of dopamine on action/inaction values** and **Effect of decision-dimensions on choice, STAR Methods**).

### The decision-space is dynamically reformed after prediction errors

Sudden shifts in activity along decision-dimensions, which we suggest above track prediction errors^61,89,140,164,165^ in certain instances, serve the important functional role of producing sudden shifts in the decision-space. Sudden changes to cortical input, for instance during a prediction error, affect the activities of each dsSPN, isSPN, dmSPN, and imSPN corresponding to the decision-dimension. Thus, a new direct pathway and indirect pathway decision-space is formed after the prediction error (**Figures 4G-I**). Continuing the earlier example, where stimuli associated with reward (sound, taste of milk) synapse to a single SPN (corresponding a reward-predominant decision-dimension), the sound of the milk poured into a glass leads to prioritization of the reward-predominant decision-dimension in direct pathway decision-space and de-prioritization of it in the indirect pathway. This way, the circuit uses the predictive stimulus (the sound) to focus on associated information (e.g. reward cues) in preparation for the expected event (choosing whether to drink milk). In this process, dopamine is released to form the new decision-space.

The circuit also has a functional mechanism to quickly transition away from a decision-space that is no longer optimal. We form our model here from experimental work demonstrating rebounds in daSNC activity^41^ and striatal dopamine release^128^ after sSPN optogenetic stimulation. We model this as changes to the connection strength between sSPN and daSNC depending on sSPN signal. This plasticity allows for rapid de-prioritization of a decision-dimension after a negative prediction error (e.g. less reward than expected), enabling a shift to a more helpful decision-space **(Figure 4J)**.

### A low-dimensional decision-space is optimal under disorder conditions

Reduced cortical signal-to-noise ratio (SNR) and disrupted cortical signaling have been observed in disorders^166^ like chronic stress^123^, which our analysis linked to reduced dimensionality of the decision-space (**Figure 3**). According to our model, this change in the decision-space is an important functional adaptation to poor quality of cortical input.

The more decision-dimensions that are used, the more sophisticated the rules for defining action values become. However, decision-dimensions are orthogonal (because they are principal components of cortical inputs), so noise is additive. Thus, the dimensionality of the decision-space can be optimized to process relevant information while limiting noise (**Figures 5A-E**, **Effect of cortical SNR on choice, STAR Methods**). Indeed, dopamine, which is released to mSPNs to form the decision-space, has been suggested to optimize SNR in the striatum^167,168^ and cortex^166,169,170^. Differential release based on cortical input quality, we hypothesize, is a mechanism behind the shifts in choice observed across disorders indicative of a low-dimensional decision-space. For example, after chronic stress, rodents ignore costs and focus only on rewards^123^, and in addiction, subjects ignores rewards other than the addictive substance^171^.

### Model predictions

#### Variation in the decision-space leads to disorder comorbidity

An untested prediction of the model is that the decision-space forms probabilistically depending on daSNC activity. Thus, a decision-space can take a variety of forms at a given activity, although the activity of the circuit produces a bias towards certain types of decision-space over others (**Figures 6A,B**, **S6A,B, Effects of sSPN, LHb, and daSNC activity on decision-space, STAR Methods**).

This prediction, if verified, may explain patterns of decision-making commonly seen in patients (**Figures 6C-E and S6C-E, Effect of decision-space on choice profiles, STAR Methods**). There are differences in circuit activity across individuals, including between control and disorder-group rodents^12,68,87,123,172–176^. So, different disorder phenotypes^167^ and individual differences in disorder presentation despite similar disorder diagnoses^79,80,178^ could arise in cases where there are slight differences in circuit activity, leading to similar decision-spaces formed between individuals, but at different rates. Daily variance in decision-making^84–86^ could arise from daily variance in circuit activity, causing daily variance in the decision-spaces formed most often. Further, differences in circuit activity may explain the inter-individual differences in the severity of psychiatric disorder symptoms^179–182^, including during decision-making^183,184^. Individual differences in disorder susceptibility^81–83^ could arise from reliance upon or avoidance of a decision-space that leads to extreme decision-making tendencies (e.g. extremely action-heavy, extremely risk-averse) when combined with abnormal action value rules in mSPNs (**Figures 6F and S6F**).

#### Different adaptation of the circuit in individuals vulnerable versus resilient to disorders

Another untested prediction of the model is that certain decision-spaces are preferred over others during certain tasks, whether as a response to a difficult decision (as in **Figure 2**) or the quality of cortical input (as in **Figure 5**). We might expect the circuit to adapt between trials as it adjusts to more frequently form a preferred decision-space (**Figures 7A-F**, **Effect of initial circuit activity on future trials, STAR Methods**).

**Figure 7.**
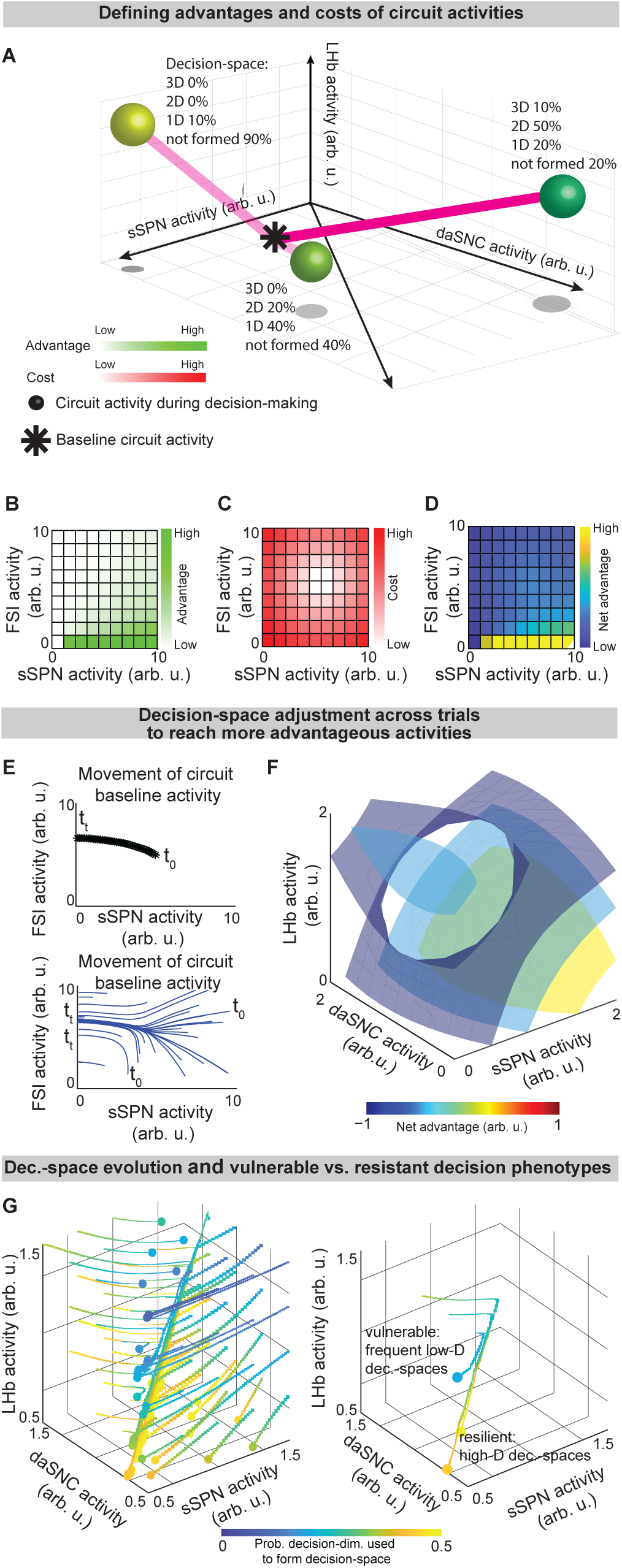
Circuit adapts between trials to form preferred decision-spaces, leading to disorder progression. **A**. Cartoon illustrating how the modeled circuit adjusts to facilitate construction of preferred decision-spaces (“advantage”) while limiting large changes in circuit activity during decision-making (“cost”). Possible advantages and costs (whose difference is “net advantage”) are shown at three circuit activities (balls). The decision-space is formed differently at each circuit activity (annotations), leading to differences in advantage. Between trials, the circuit adjusts its baseline activity (i.e. resting activity outside decision-making) in the direction of highest net advantage. **B-D**. Simulated advantages (**B**), costs (**C**), and net advantages (**D**) shown across sSPN and FSI activities. **E**. sSPN and FSI activity adjusts between trials to best form preferred decision-spaces. Due to this, the circuit adjusts its activity from an initial baseline activity (*t*_0_) to other activities associated with the required decision-spaces (*t_t_*). **F**. Similar to **D** but for sSPN, LHb, and daSNC activities, showing a simulated circuit where it is most advantageous to have high sSPN activity during simple choices. The trend in the continuous 3D decision-space is visualized using isosurfaces. **G**. Similar to **E** but for sSPN, LHb, and daSNC activities. Dots show ending circuit activities (i.e. *t_t_*) and beginnings of lines possible initial circuit activities (i.e. *t*_0_). Depending on initial activity, the circuit may increasingly use or disregard a decision-dimension when forming decision-space. Right panel shows trajectories from three starting circuit activities, two of which lead to the decision-dimension commonly being used to form decision-space (e.g. resilient subjects) and one which leads to it commonly being disregarded (e.g. vulnerable subjects).

We can use this prediction, if verified, to frame vulnerability or resilience to disorders as an adaptation that is favorable (e.g., to adeptly form a high-dimensional decision-space) or maladaptive (e.g., to only form a low-dimensional decision-space, regardless of decision context). Depending on its initial activity, a modeled circuit can adapt to reach very different activity, leading to disposition to either a high-or low-dimensional decision-space (**Figures 7G and S7A**). Thus, differences in the circuit before exposure to a traumatic event, for instance, may explain why two subjects that encounter the same traumatic event do not always develop the aberrant decision-making symptoms of PTSD^185,186^. It may also shed light on the neural processes underlying incubation of fear^187^ and incubation of craving^188–191^, where disorders progress over the span of weeks or months, even when the traumatic event or addictive substance does not reappear (**Figures S7B-D, Effect of altered advantage score on future trials, STAR Methods**).

## Discussion

Our findings suggest that the cortical-striatal-GPi-LHb-RMTg-daSNC circuit forms a mathematical space during decision-making which it uses to define action values. Our model, by incorporating circuit functions commonly observed within the brain like orthogonality^192–194^ and dimensionality reduction^99,125^ and fitting with the literature on regional functions observed in decision-making^14,15,17,36,61,72,132,132,134,135^, sheds light on: 1) the intersection of the indirect and direct pathways with sSPNs and mSPNs, 2) the functional roles of sSPNs and mSPNs; 3) the functional role of daSNC, and 4) the contribution of the circuit to neuropsychiatric disorders. Our model aligns with known patterns in decision-making, including SPN response to reward prediction errors^87,89,93,140,165,195^, selecting between two conflicting offers^11,196^, and fluctuations in subjective valuation^197^. Our model unifies these findings by explaining how neural processes give rise to mathematical computations which affect decisions. Additionally, our work offers a new perspective on other striatal modeling work (action selection^48^, proximity theory^198^).

### Impact of dopamine on striosomal circuit function

Experimental evidence indicates that the matrix and striosomal compartments have different dopaminergic dynamics^199,200^ which may arise due to the developmental characteristics of striosomes^201^. The dopaminergic mechanisms in our model are aligned with the finding that dopaminergic activation of the direct pathway in the striatum had opposing effects on up-states (where the neuron’s resting membrane potential is closer to an action potential) in the matrix and striosomes^156,199^. These differential dopamine dynamics within each compartment aligns with the differential roles the striosomes and matrix compartments have during decision-space “formation” Other experimental research demonstrates that dopamine is generally released faster in the striosomes than the matrix^202^ and more dopamine is released in the matrix compared to the striosomes^203,204^. This model also suggests that dopaminergic rebounds^41^ along with altered dopamine dynamics^128^ facilitates decision-space switching and adjustment by allowing new decision-dimensions to be integrated or removed from decision-space. Alterations to the decision-space may then lead to different action/inaction valuations potentially causing choice selection to change.

### Developing enhanced techniques for neuropsychiatric disorder diagnosis and treatment

Perturbing and understanding internal states^205–207^ are facets of behavioral research that may be critical to exploring decision-space. For example, shifts between observable states may be used for measuring shifts in decision-space. Linking quantifiable decision-making to internal decision-spaces and the expected circuit function could enhance the granularity of diagnostic methods/tools while allowing for accurate individualization of treatment plans. Mapping decision-space characteristics and shifts may improve the identification of individual differences in susceptibility to disorders by leading to the identification of mechanisms underlying the susceptible and resilient models currently being used^208–210^. Importantly, both the direct and indirect pathways and striosome-matrix balance are impaired in various neuropsychiatric disorders^37,42,75,157^ and clarifying the interaction between them may explain more disorder symptom manifestations as well as direct research towards potential treatment targets based on the expected aberrancies suggested by the model. Additionally, decision-space and the proposed function of each individual brain region during decision-making provided by our model could guide the development of algorithmic/artificial^211,212^ intelligence tools that will be beneficial for diagnosing neuropsychiatric disorders.

Overall, this model suggests mechanisms underlying individual differences in disorder susceptibility^81–83^, day-by-day variance in behavior/disorder symptom expression^84–86^, and similar decision-making behaviors observed in different disorders^177^. Our model provides a framework for linking biological processes to commonly observed decision patterns. Further, our model could also lead to the development of individualized biofeedback^213^ methods that can improve decision-making by identifying decision-spaces and the associated activity of brain regions, thereby potentially identifying the most important areas for improving decision-making. Overall, our model provides a theoretical foundation that unifies experimental and modeling work while also demonstrating how brain regions and pathways could physiologically function during decision-making.

### Limitations

Several observations would disprove the decision-space model: 1) sSPNs are not organized into subpopulations encoding different representations of cortical activity (each corresponding to a decision-dimension); 2) dopamine release from daSNC neurons is not the primary connector between sSPN and mSPN subpopulations encoding similar information (enabling the formation of decision-space); 3) the value of an action does not depend on the activities of multiple mSPN subpopulations, considered cumulatively (action values depend on the decision-space); or 4) direct and indirect pathway SPNs do not influence the value of actions and refraining from actions, respectively (direct and indirect pathway decision-spaces).

The model does not include certain brain regions such as the dorsolateral striatum, ventral striatum, dopamine in the ventral tegmental area, Globus Pallidus externus, Substantia Nigra reticula, subthalamic nucleus, and other basal ganglia regions. We did not consider neuronal molecular heterogeneity. Despite its focus on several brain regions, the model has demonstrated success in relating neural activity to decision-making.

## Acknowledgements

We greatly appreciate G. Schoenbaum, and Y. Shaham for their constructive criticism and discussions during model development. This project was supported by the NSF/CAREER (#2235858), NIH/NIDA (#R01DA058653), U-RISE T34GM145529 and G-RISE T32GM144919.

## Author Contributions

conceptualization: A.F., K.A.G.; data curation: D.W.B., L.D.D., L.R.I., S.B.D., A.G. Q.Z.; formal analysis: D.W.B., L.R.I., L.D.D., C.N.H., A.G., Q.Z., D.T., M.P.; funding acquisition: A.F., K.A.G.; investigation: A.F., D.W.B., L.R.I., L.D.D., C.N.H., Q.Z., S.B.H., A.Y.M., A.G., K.N., K.A.G.; methodology: A.F., D.W.B., S.B.D., L.R.I., A.A.S., N.F.R., R.J.I.; project administration: A.F., K.A.G.; software: A.F., D.W.B., L.R.I., L.D.D., C.N.H., Q.Z., A.G.; supervision: A.F. K.A.G.; validation: A.F., D.W.B., L.D.D., A.G., Q.Z., S.M.D., L.R.I.; visualization: D.W.B., S.B.H., P.V., S.A.B., A.Y.M., C.N.H., R.J.I., A.A.S., N.F.R.; writing – original draft: A.F., D.W.B., C.N.H., S.U.B., K.A.G.; writing – review & editing: A.F., D.W.B., C.N.H., K.A.G., T.M.M., L.E.O., M.P.

---

**Figure S1.**
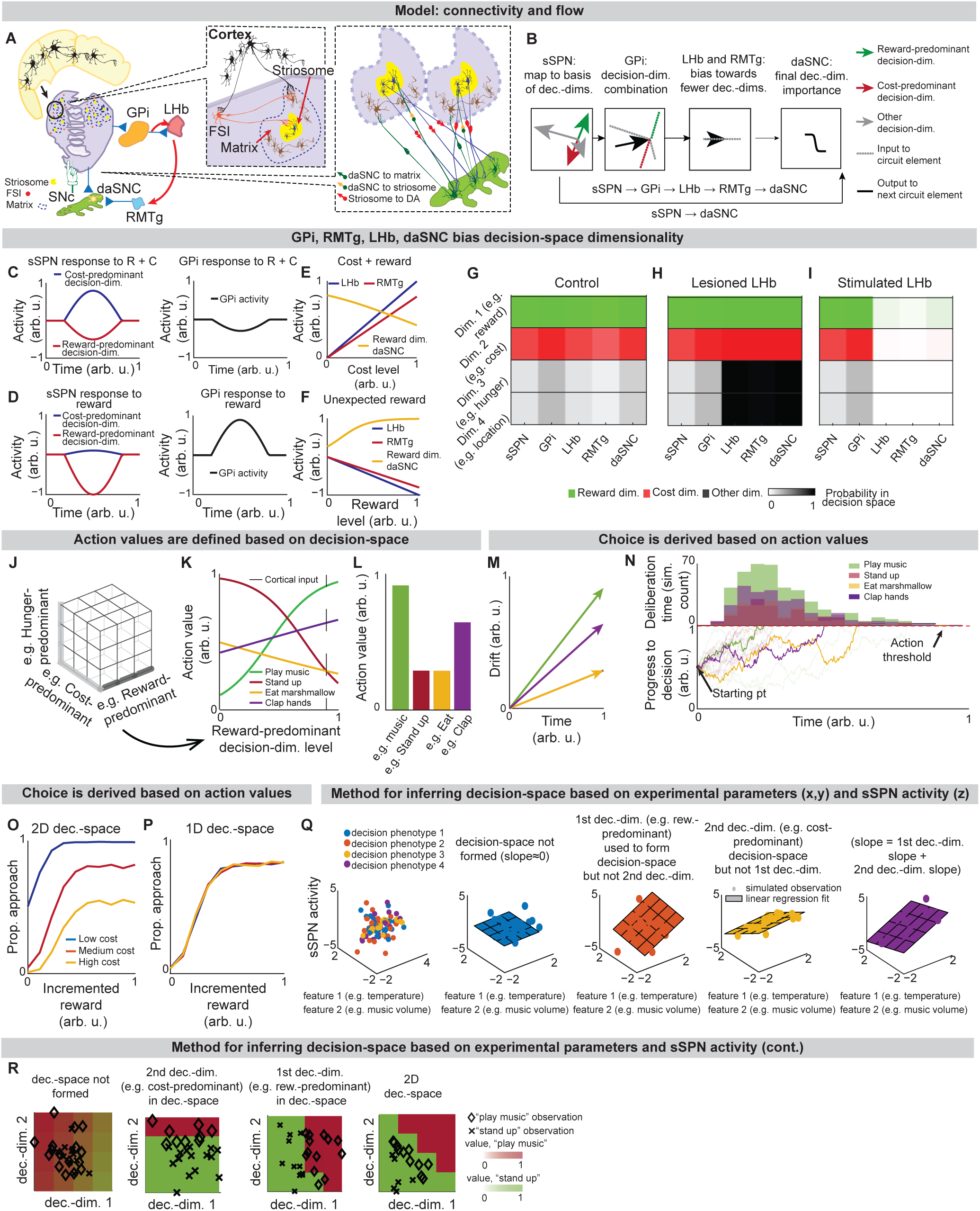
Action values are defined within a decision-space formed by the circuit, Related to Figure 1 (65/85) **A.** The circuit. Red = excitatory, blue = inhibitory connections. In this figure (and Figure 1), an instance of the model is described which allows for two convenient simplifications: 1) the representation of SPNs (both sSPNs and mSPNs) using one circuit element per decision-dimension, and 2) the treatment of the circuit using a feedforward model. See ***Instance 1: full connectivity and feedforward,* STAR Methods**. **B.** After FSI normalization, sSPNs map cortical activity to a basis of decision-dimensions. Mathematically, we represent, for a given pathway *P* (either direct pathway or indirect pathway), the mapping of cortical activity x_*P*_ to decision-dimensions as a linear transform via the matrix **W***P*, whose columns correspond to the first several (in our analysis, 4) principal components of cortical activity. During this process, there is divisive normalization by FSI activity *c_P_* and the potential for an overall shift in sSPN activity *b*_sSPN_ (eq. (1)). Next, GPi combines the sSPN signals into a single representation and RMTg and LHb bias this combined representation of signal along all decision-dimensions. We represent RMTg activity RMTg as the dot product of striosome to GPi weights **w**_GPi_ and the activities the sSPNs s_sSPN,*P*_ for each pathway *P*, after incorporation of additive shifts *z*_GPi_, *z*_LHb_, and *z*_RMTg_ (eq. (5)). Next, daSNC takes input directly from sSPNs and from RMTg to calculate the final importance of each decision-dimension. We represent daSNC activity daSNC_*i,P*_ corresponding to decision-dimension *i* and pathway *P* as the activity of the corresponding sSPN population s_sSPN,*i,P*_ multiplied by a connection weight *w*_sSPN→daSNC,*I,P*_, plus additive shifts applied individual to each daSNC element (*z*_daSNC,*i,P*_) and uniformly to all daSNC elements (RMTg), all passed through an activation function (eq. (2)). **C,D**. Example where sSPNs parse reward and cost (**C**), or reward (**D**) inputs along decision-dimensions (left) and then GPi combines the signals along the decision-dimensions into a single representation (right). **E,F.** Modeled responses of LHb, RMTg, and daSNC to a cortical signal encoding reward and cost (**E**), and a cortical signal encoding an unexpected reward (**F**). LHb and RMTg increase their activities proportionally to the cost and decrease their activities proportionally to the reward. daSNC change their activities inversely. The modeled daSNC response is due to the combination of cortical inputs projected onto them directly from sSPN and through GPi, LHb, and RMTg. **G-I**. Roles of the circuit elements in determining which dimensions form decision-space. Three scenarios are shown: control (*z*_LHb_= 0.5 in eq. (5)), lesioned LHb (*z*_LHb_ =− 5), and stimulated LHb (*z*_LHb_ = 5). The colors shown for each circuit element correspond to the decision-space that would be formed absent the influence of downstream circuit elements. **J-L**. Process by which action values are defined using decision-space. During a decision, a subset of decision-dimensions is selected, forming decision-space (**J**). This is represented mathematically through a diagonal matrix **S***P* whose elements are set probabilistically to either 1 (dimension in decision-space) or 0 (dimension not in decision-space) (eq. (3)). Otherwise, mSPN activity is formulated similarly to sSPN activity. Rules corresponding to retained decision-dimensions are used to define action values for action *j* and pathway *P* (**K,L**). This is represented mathematically as multiplication of a vector by mSPN activity , subtracted by a shift and run through an activation function (eq. (4)). **M,N.** Action (or inaction) values *v_j,P_* for each action *j* and pathway *P*are used as drift rates (**M**) in a Merton process model (**N**). Discrete Merton processes obtained as a constant time step discretization of eq. (6) are run for each action simultaneously. At the time the first of the *v*_*j*,direct_ processes reaches a defined threshold, the corresponding action is enacted unless its inaction process *v*_*j*,indirect_ has reached *h* first (eqs. (7),(8),(9)). Lines show the progress of processes towards a decision threshold for an example simulation. Histogram shows the decisions and deliberation times for the processes that reached the threshold first. **O,P**. Psychometric functions derived across multiple reward levels. In the modeled experiment, the subject is asked to evaluate the rewards and costs of two offers and either approach or avoid. Decision-space (**O**: 1D, **P**: 2D) affects choice patterns. **Q,R**. Demonstration of a method by which decision-space can be inferred from sSPN activity, showing the utility of the mathematical formation of the model in connecting experimental inputs, sSPN activity, and choice. The method demonstrated here could be used to design an experiment in which the decision-space model is tested, or it could be used to explain differences in choice between groups, for instance control and disorder. In the simulation, two environmental inputs (e.g. temperature, music volume) across 100 sessions are classified based on decision-making phenotypes, for example based on reaction time, heart rate, eye movement (colors). Using the method visualized here, the decision-making phenotype classes are assigned one of four decision-space reference labels: 1) where decision-space is not formed, 2) where only the first decision-dimension is used to form decision-space, 3) where only the second decision-dimension is used to form decision-space, and 4) where both decision-dimensions are used to form decision-space. Synthetic data is generated by forming an arbitrary set of cortex→sSPN weights, using these to form sSPN activity, and then adding i.i.d. Gaussian noise. A linear regression is used to derive estimated decision-dimensions and assign hypothesized decision-spaces to each label (**Q**). Choice is then examined with respect to the derived decision-dimensions. As expected, the regression slope (planes) corresponding to the 2D decision-space is roughly the sum of the regression slopes of the two 1D decision-spaces (**Q**), and decisions correlate with the dimensions hypothesized to be used to form decision-space when choices are plotted against hypothesized dimensions (**R**). In the colormaps in **R**, action values are interpolated from an example set of observations (diamonds and Xs) via logistic regression, and a boundary line is drawn where the action value of “play music” equals the action value of “stand up.” 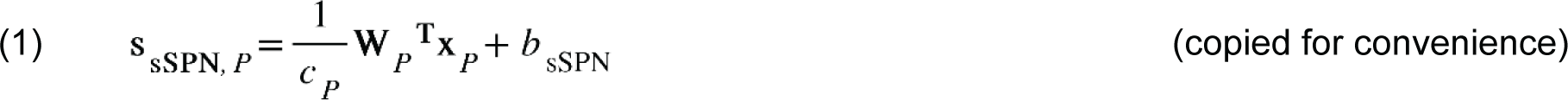

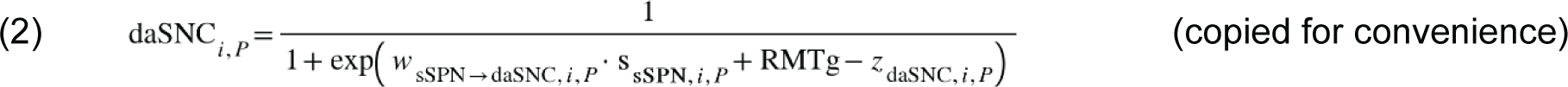

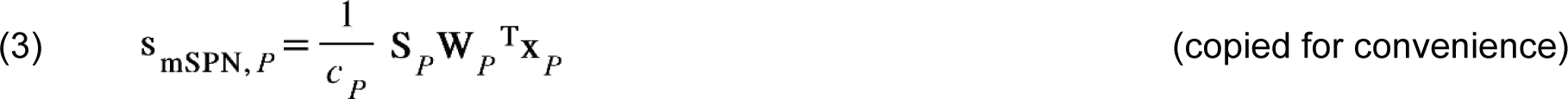

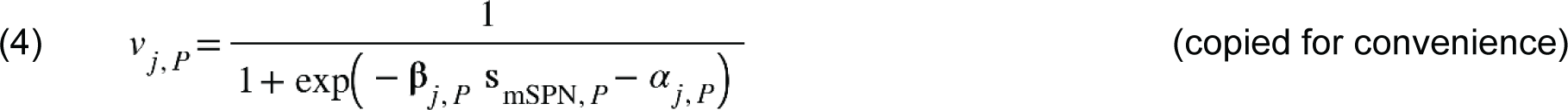

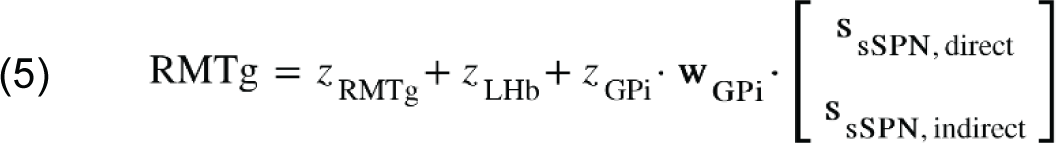

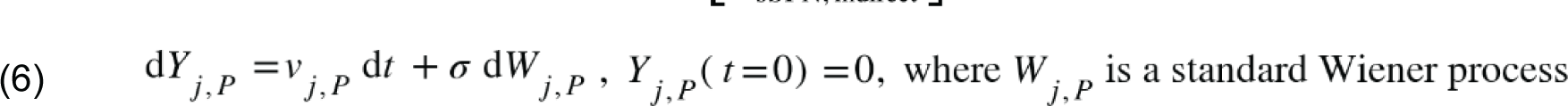

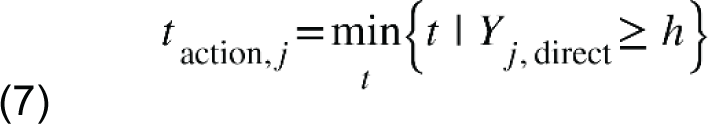

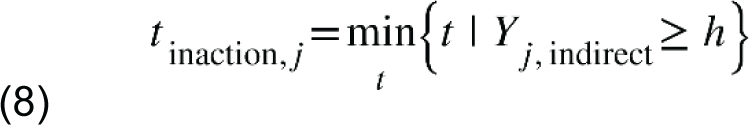

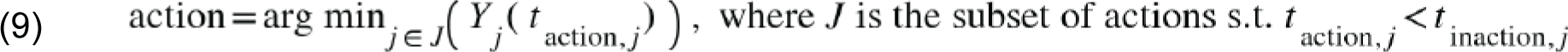

**Figure S2.**
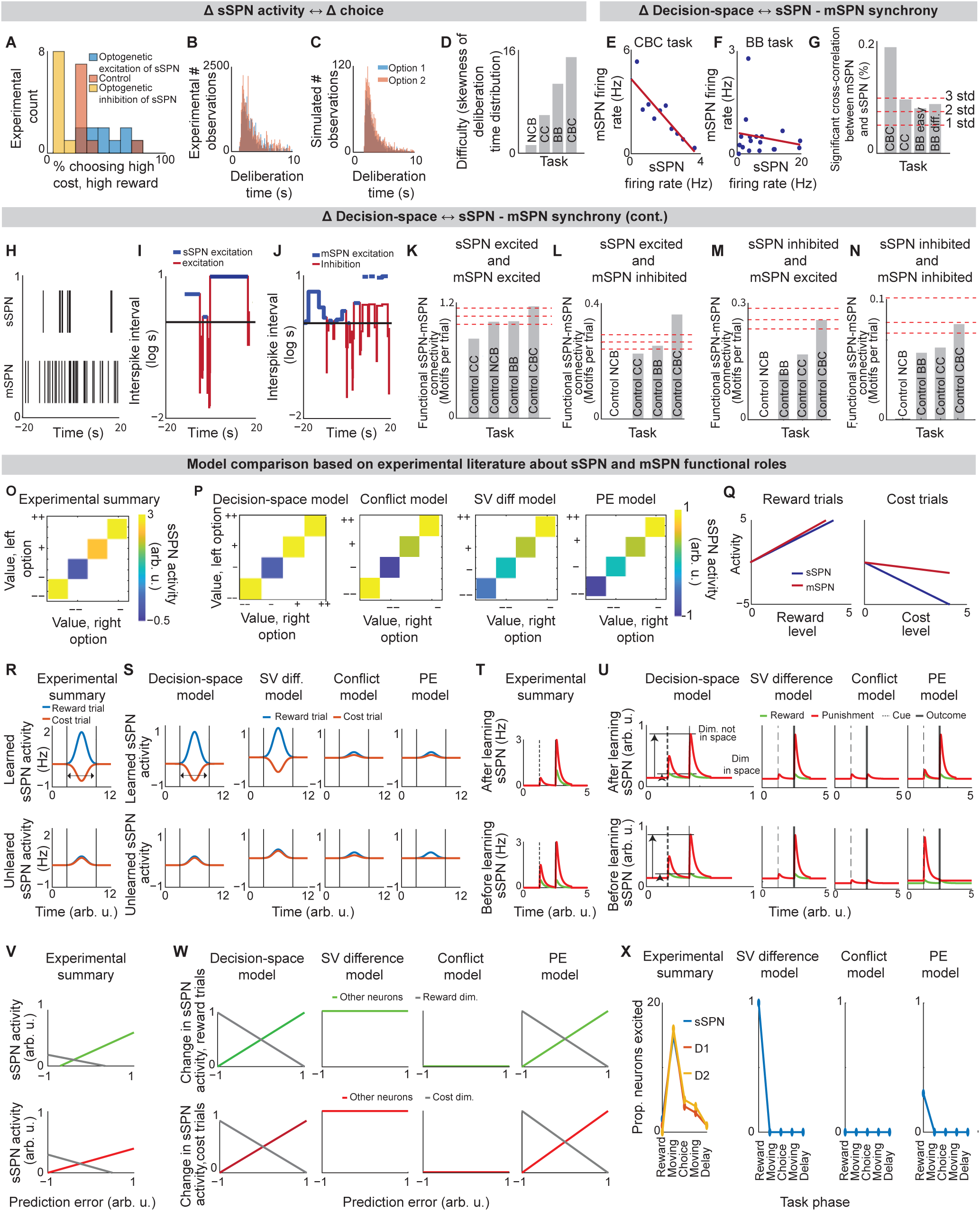
Decision-space model validation and comparison to alternative models, Related to Figure 2. (88/85) **A.** Summary of experimental results of optogenetic manipulation during a conflict task in Friedman et al. (2015). Resembles the model in Figure 2B. **B,C.** Experimental deliberation time distribution (6 animals, 35 sessions) (**B**), which is successfully modeled using the Merton process model (**C**). Distribution is for the benefit-benefit task (concentration 70%). Experimental data here and throughout the figure is analyzed from the Corticostriosomal Circuit Stress Experiment database. **D**. Skewness of the deliberation time distribution, which is used to estimate task difficulty. The CBC task had a deliberation time distribution that was more skewed than the non-conflict tasks. Tasks: NCB = non-conflict cost-benefit (4 rats, 27 sessions, 1250 trials), CC = cost-cost (7 rats, 25 sessions, 1852 trials), BB = benefit-benefit (7 rats, 128 sessions, 7762 trials), CBC = cost-benefit conflict (8 rats, 69 sessions, 3921 trials). **E,F**. Analysis of relationship between sSPNs and mSPNs during decision-making. sSPN and mSPN neurons are significantly more correlated in tasks that require integration of reward and cost versus only reward or only cost. **E** shows representative examples from the cost-benefit conflict task (CBC, both reward and cost) and **F** shows the benefit-benefit task (BB, only reward), respectively. **G.** The CBC task had significantly more correlated (Pearson’s r^2^ > 0.4 and significance < 0.05) pairs than the tasks which required integration of only reward or only cost. Confidence intervals (dashed red lines, 1,2,3 standard deviations) are estimated based on shuffled data. NCB = non-conflict cost-benefit (sSPNs = 14, mSPNs = 260), CC = cost-cost (sSPNs =46 , mSPNs = 400), CBC = cost-benefit conflict (sSPNs = 84, mSPNs =717), BB = benefit-benefit easy (sSPNs = 50, mSPNs =515 , chocolate milk concentration <50), BB = benefit-benefit difficult (sSPNs =33, mSPNs = 731, chocolate milk concentration >=50) **H-J.** Process by which we identify functionally connected sSPN and mSPN neurons. Spiking times (**H**) are used to find inter-spike intervals (**I**,**J**) for sSPN (top rows) and mSPN (bottom rows) that were recorded simultaneously during decision-making. Intervals above the median interval (black line) are classified as inhibition (blue) and below median are classified as excitation (red). Functional connections are determined based on whether excitation or inhibition of sSPN followed by excitation or inhibition of mSPN. **K-N.** Significantly more sSPN and mSPN neurons were functionally connected during decisions that required integration of reward and cost (CBC task) than the other tasks for all types of connections: sSPN excited and mSPN excited (**K**), sSPN excited and mSPN inhibited (**L**), sSPN inhibited and mSPN excited (**M**), and sSPN inhibited and mSPN inhibited (**N**). Tasks: NCB = non-conflict cost-benefit (sSPNs = 14, mSPNs = 260), CC = cost-cost (sSPNs =46, mSPNs = 400), BB = benefit-benefit (sSPNs =83, mSPNs = 1246), CBC = cost-benefit conflict (sSPNs = 84, mSPNs =717) **O.** Mean sSPN activity should track decision-space dimensionality. Tasks in Friedman et al. (2015), plotted in order of difference between reward and cost, are assessed for likely decision-space dimensionality based on task difficulty (**Figure S2D**). Those with higher task difficulty are assigned lower sSPN activity per the model in Figure 2C. **P.** Three alternative models applied to the task in Friedman et al. (2015). The subjective value (SV) model assumes that sSPN encoded the relative action values of the two arms of the T-maze. The conflict model assumes that sSPN activity is inversely proportional to the amount of conflict in the task. The prediction error model compares expected value entering the task with reward or cost obtained on the maze. The conflict method resembles experimental results but not the SV difference model or the conflict model. **Q,R.** Summary of differences in sSPN activity across trials in the operant conditioning task in Friedman et al. (2020). mSPNs and sSPNs were active during reward trials, suggesting per the model in Figure 2C that decision-space was not formed (**Q**). sSPNs were less active during the cost trials and were differently active than mSPNs, suggesting formation of a low-dimensional decision-space per the models in Figures 2C **and E**, (**R**). **S.** Prediction of sSPN activity for the task in **R** for the decision-space model and three alternative models. sSPN activity scales with overall subjective value in the task, so the subjective value model successfully interprets the experimental results. **T,U.** Experimental results are summarized from Xiao et al. (2021) (**T**). Prediction of the decision-space model and three alternative models of sSPN subpopulation activity at the cue and during the outcome period of a Pavlovian conditioning task (**U**). Per the decision-space model, sSPN subpopulations should respond similarly to the cue and to the outcome because associated data is likely mapped along the same decision-dimension. The subjective value model expects more activity during the outcome period when reward is administered. The prediction error model expects more activity at the cue after learning. There was no conflict in the experimental setup. **V,W.** Experimental results are summarized from Bloem et al. (2022) (**V**). Predictions of the decision-space model and the three alternative models of sSPN activity during a probabilistic Bandit task (**W**). The decision-space model expects that sSPN activity will reveal prediction errors, as does the prediction error model. The subjective value model instead anticipates that activity will track overall value regardless of prediction error. There was no conflict anticipated by conflict model. **X.** In the value-guided choice task in Weglage et al. (2021), activities of all neuron types recorded (sSPN, dmSPN, imSPN) resembled one another over phases of the task and did not solely track subjective value, prediction error, or conflict. As shown in Figure 2E, the experimental finding is an expectation of the decision-space model when a high-dimensional decision-space is formed.

**Figure S3.**
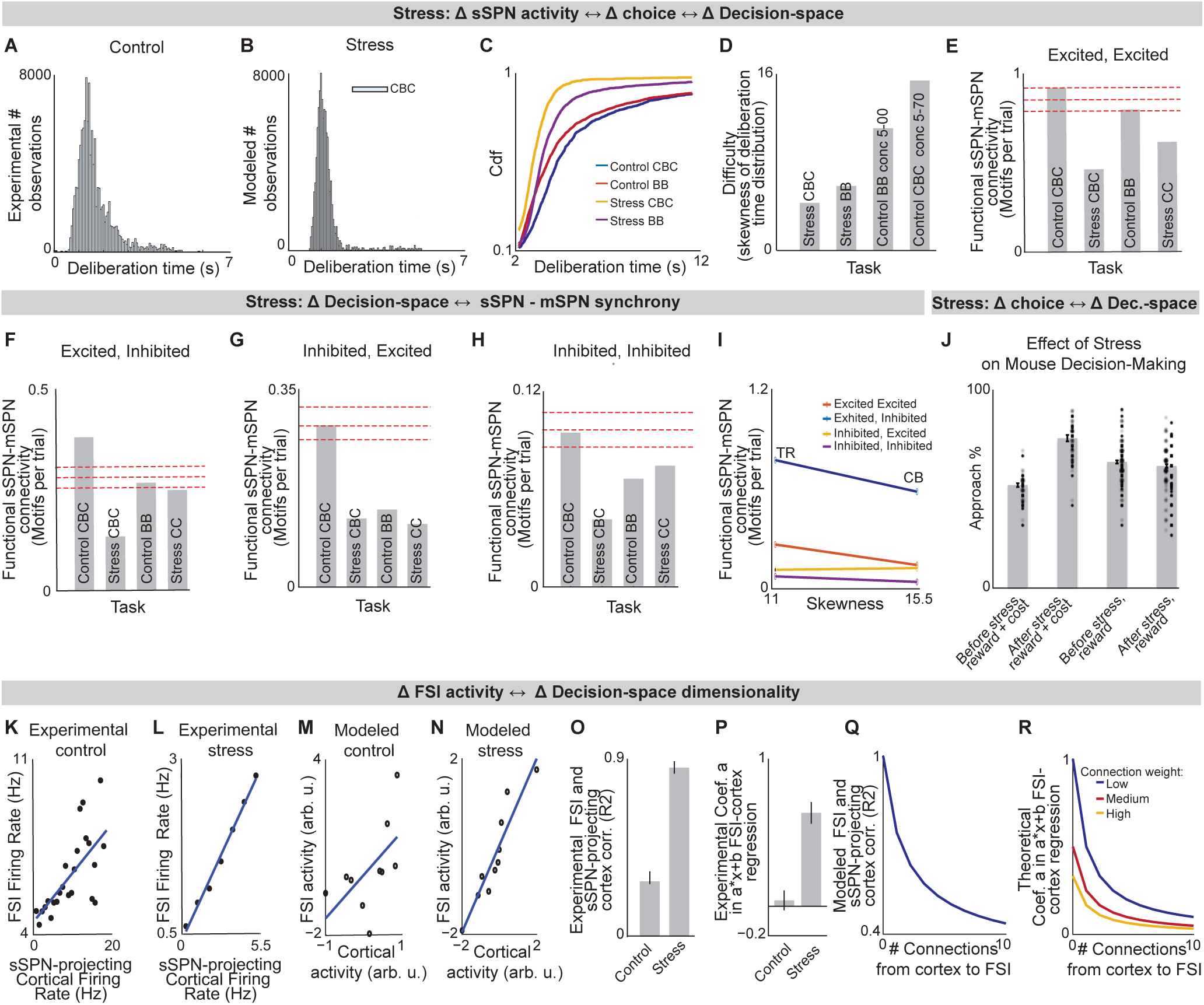
Analysis of neural and decision-making data shows that decision-space is changed after stress, Related to Figure 3. (84/85) **A,B.** Deliberation time distributions of rodents performing the cost-benefit conflict task before stress (**A**) are less skewed and have shorter deliberation time than after stress (**B,** control: 8 rats, 198 sessions, 11683 cells; stress: 5 rats, 138 sessions). **C,D.** Cumulative distribution functions (**C**) and distribution skewness (**D**) of deliberation time show that after chronic stress, the task involving integration of both reward and cost (CBC task) changes from producing the slowest (blue) to the quickest (yellow) decisions. CBC choice was slowest in control rats (p < 0.0001, KS-test). After stress, CBC choice was faster than in the other tasks (p<0.0001). Control CBC task: 69 sessions, 8 rodents; Control BB task: 128 sessions, 7 rodents; Stress CBC task: 34 sessions, 5 rodents; Stress BB task: 104 sessions, 5 rodents. **E-H.** sSPNs and mSPNs were more functionally connected during a difficult (CBC) task before stress than after, per all possible types of connection: sSPN excited, mSPN excited (**E**), sSPN excited, mSPN inhibited (**F**), sSPN inhibited, mSPN excited (**G**), and sSPN inhibited, mSPN inhibited (**H**). Significance levels, depicted by dashed lines, show one (bottom), two (middle), and three (top) STD for functional connections calculated from shuffled control data. Control cost-benefit conflict (CBC) task: 92 pairs, 8 rodents; Tasks: Control BB = benefit-benefit (sSPNs =83, mSPNs = 1246), Control CBC = cost-benefit conflict (sSPNs = 84, mSPNs =717), Stress CBC = cost-benefit conflict (sSPNs =41, mSPNs =898), Stress BB = benefit-benefit (sSPNs = 156, mSPNs = 2813). **I.** Deliberation time distribution skewness is no longer linked to functional connectivity after stress. **J.** Experimental data that inspires the model in Figure 3H. Mice that underwent chronic stress approached the lower-reward arm of the T-maze less when a small cost was added (“reward + cost” = cost-benefit conflict task, “reward” = benefit-benefit task). Dots = individual sessions, bar = mean across trials. There is significant difference (p<10^-19^, paired t-test) in choice for the CBC task before and after stress, and nonsignificant difference in choice for the BB task before and after stress (p=0.10). CBC, before stress: 17 rodents, 38 sessions; CBC, after stress: 13 rodents, 34 sessions; BB, before stress: 23 rodents, 114 sessions; BB, after stress: 14 rodents, 116 sessions. Our model interprets this result as due to differences in the decision-space. **K-N.** Representative examples of simultaneously recorded FSIs and prelimbic cortex neurons firing rates before (**K**, Pearson’s R=0.46) and after (**L**, R=0.99) chronic stress. After stress, there is less coordination between the connected pairs. This can be modeled as a reduction in the number of cortical neurons that synapse to each FSI (**M,N**). **O-R.** In general, the neuron pairs in rats that underwent chronic stress had significantly higher correlation (**O**, p<10^-18^) and significantly higher values of slope “a” in their a⋅x+b linear regression fits (**P**, p<10^-6^). Control: 7 rodents, 78 neuron pairs; Stress: 4 rodents, 37 neuron pairs. This suggests, per our modeling, that there are fewer connections from cortex to FSI after stress. Modeled squared Pearson correlation coefficient (**Q**) and slope “a” parameter in the a⋅x+b fits (**R**) are shown when the connection between cortex and FSI neurons are altered in two ways: 1) through a reduction in the number of connections, and 2) through a reduction in the strength of each connection (i.e. connection weight). This experimental evidence is aligned with a reduction in the number of connections, suggesting that FSI normalization is disrupted after stress, leading to higher sSPN activity and thus formation of lower-dimensional decision-spaces.

**Figure S4.**
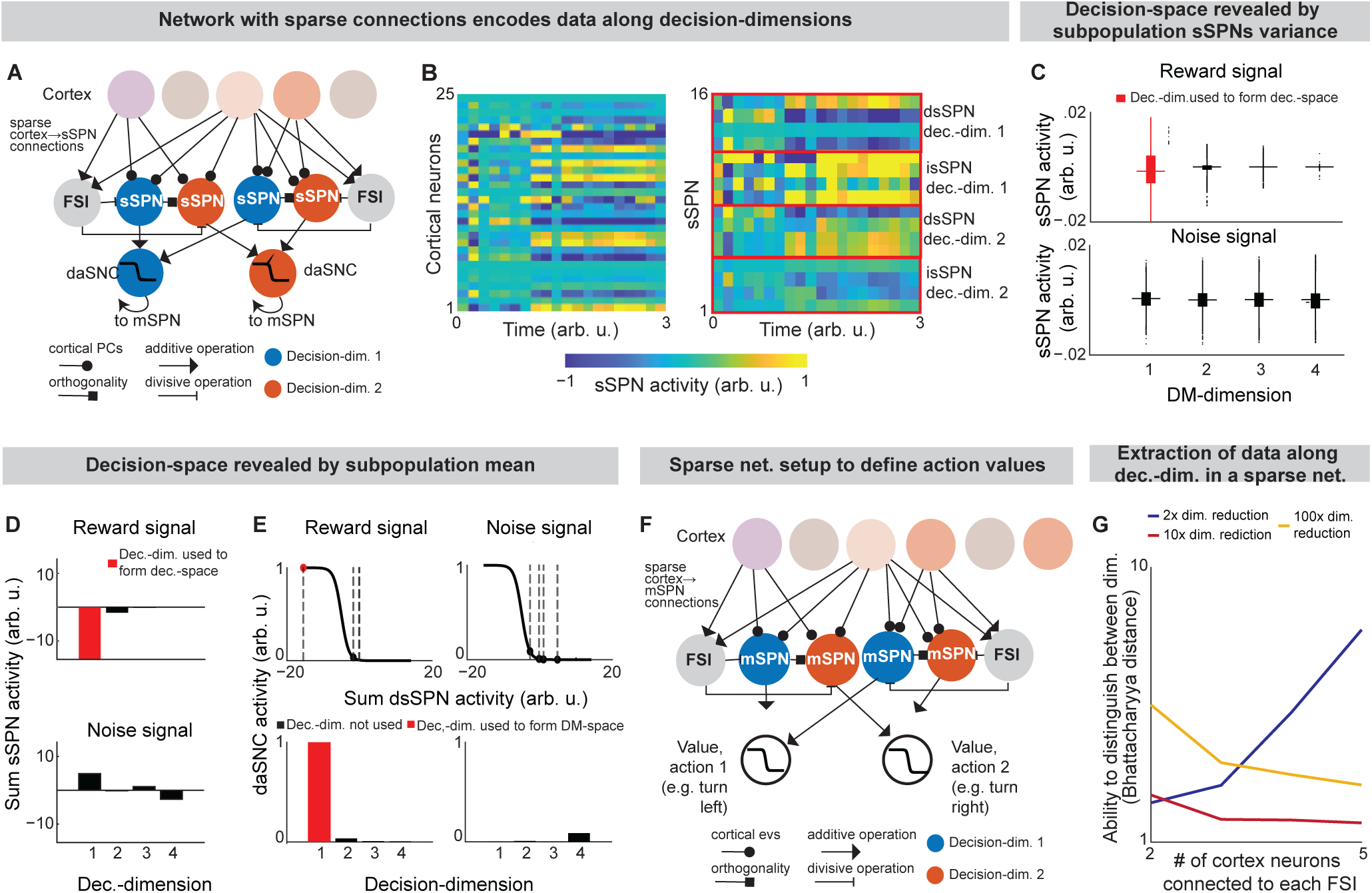
Mean and variance of SPN activities reveals decision-space, Related to Figures 2,3. **A,B**. In this figure, we demonstrate that SPNs can encode activity along decision-dimensions successfully even in large, sparse networks, as exist in the brain, by considering an instance of the model where there are sparse connections between cortical neurons (n=50), FSIs (n=10,000), and SPNs (40,000 sSPNs, 40,000 mSPNs). sSPNs each encode activity along a principal component of a randomly sampled set of cortical neurons *C* (**A**). So, when activity changes in *C* (for example, during the reward cue in **B,** left panel), sSPNs (**B,** right panel) that each encode data along an *i*th principal component respond somewhat similarly to one another. See ***Instance 2: sparse connectivity and feedforward*, STAR Methods**. **C.** Decision-space is revealed from the variance of sSPN activities, aligning to the experimental result in Figure 2F. Modeled sSPN subpopulations have greater variance when there is high-magnitude cortical signal along a corresponding decision-dimension (e.g., an sSPN subpopulation corresponding to a reward-predominant decision-dimension in response to a reward cue). Summaries are shown of the activities of 10,000 dsSPN (top row). Two types of cortical signal are passed to SPNs: one with reward (left column, matches the example shown in **Figure S4B**) and one with Gaussian white noise (right column). To produce the network, 10,000 groups of 4 randomly sampled cortex neurons (notated as the set *C*) are connected to an FSI and 4 dsSPNs (or isSPNs) and 4 dmSPNs (or imSPNs) for each pathway. The activity on a given sSPN which receives projection from *C*, sSPN*s,C*, is defined mathematically based on the weights from cortical neuron *q* to sSPN *s*, *w*_*q→s*_, the activity of the FSI which received projection from *C*, FSI*_C_*, and an additive shift that represents the relative activity of all sSPN neurons, (eq. (10)). **D.** Decision-space is revealed by the mean of sSPN activities, aligned with the experimental result in **Figure S2P**. An sSPN subpopulation has lower mean activity when there is high-magnitude cortical signal along a corresponding decision-dimension. **E.** Selection process by which daSNC neurons determine which decision-dimensions to include and to not include in decision-space, illustrating the construction of decision-space from sSPN activity. Lines are daSNC activation functions. Placement along the x-axis (emphasized by dashed lines) is the subpopulation average activity in **D**. Mathematically, daSNC neuron corresponding to decision-dimension *i* and pathway *P*, daSNC_*i,p*_, is computed as the weighted average of sSPNs corresponding to that decision-dimension and pathway, shifted by RMTg activity RMTg and a daSNC biasing factor *z*_daSNC,*i,P*_, all passed through an activation function (eq. (11)). **F.** Correlate to the sSPN-centered subnetwork described in **Figure S4A** for mSPN, illustrating how action values could be defined by a network with sparse connections. The activity of a given mSPN which receives projection from *C*, mSPN_*m,C*_, is defined similarly to an sSPN but for term representing dopamine signaling from the daSNC neuron corresponding to decision-dimension *i* and pathway *P* to an mSPN corresponding to the same decision-dimension and pathway, *d_i,P_* (eq. (12)). **G.** Bhattacharya distance between the distributions of SPNs encoding data along the reward-predominant and cost-predominant decision-dimensions. sSPN can correctly differentiate reward from cost signal despite sparse cortico-striatal connectivity. Lines show the averages of 1000 simulations. This result demonstrates the feasibility of data along decision-dimensions being encoded by neural populations with sparse connections. 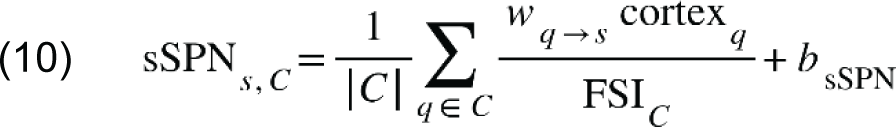

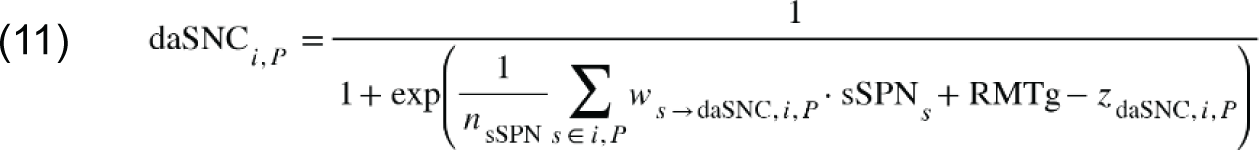

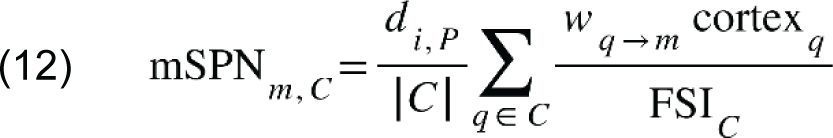

**Figure S5.**
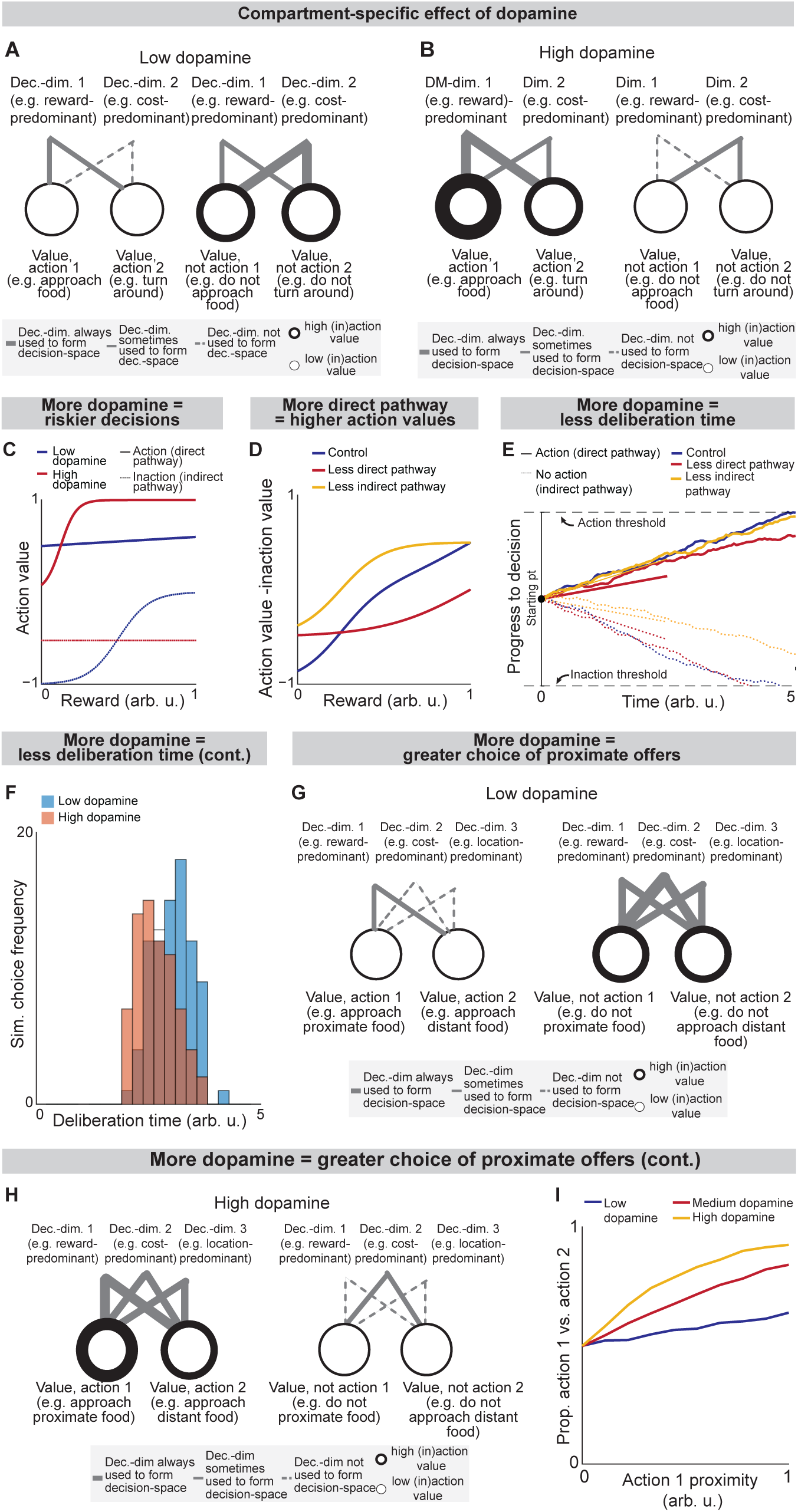
Roles of the direct and indirect pathways, Related to Figure 4. (80/85) In **Figure S5**, analysis is conducted using the instance of the model described in **Figure S1**. **A,B.** Effect of low (**A**) versus high (**B**) dopamine release on decision-space formed by the direct pathway (left panels) or indirect pathway (right panels). When dopamine release is low (**A**), low-dimensional direct pathway decision-spaces are constructed by dsSPNs and high-dimensional indirect pathway decision-spaces are constructed by isSPNs. The opposite happens when there is high dopamine (**B**). To analyze these effects, analysis in Figure 5 uses an instance of the model where the circuit elements interact dynamically, represented mathematically through a system of differential equations, where sSPN activity for a given decision-dimension *i* and pathway *P*, *s_i,P_*(*t*), respond to cortical input after normalization by FSI, *x_i,P_*(*t*), based on weights *w* between sSPNs, mSPNs, and daSNC elements, a decay factor *τ*, and a coefficient that controls sSPN→daSNC plasticity, *K* (eqs. (13),(14),(15),(16)). The weight of a decision-dimension in mSPN, S_*i,P*_(*t*), occurs dynamically depending on whether or not dopamine release is above a specified threshold (eq. (17)). See ***Instance 3: full connectivity and dynamics,* STAR Methods**. In **Figure S5**, we conduct computational analysis using ***Instance 3***, as defined in **Figure S1**. **C.** Modeled action values for the cost-benefit conflict task where an incremented reward (from a low reward of 0 to a large reward of 1 arb. u.) is accompanied by a constant cost (set to 0.25 arb. u.). Increased dopamine leads to increased action value of high-reward, high-cost options and decreased “inaction value,” thus leading to more approaches when there is high reward and high cost. **D.** The effects of the direct versus indirect pathways are examined in the cost-benefit conflict task used in Figures 2 **and 3**. Modeled action values change when dmSPN and dsSPN or imSPN and isSPN are inactivated during a task with incremented reward (from a low reward of 0 to a large reward of 1 arb. u.) with medium cost (set to 0.5 arb. u.). Inactivated dmSPNs and dsSPNs lead to lower action values and inactivated imSPNs and isSPNs lead to larger action values. **E,F.** Modeled deliberation time for the cost-benefit conflict task is shorter when there is more dopamine. Choice is modeled from action values calculated in the cost-benefit conflict task (here, reward = 1 arb. u., cost = 0.5 arb. u.). The method to derive choice from action values (upward-sloping drifts) and inaction values (downward-sloping drifts) is shown in **E** and modeled deliberation times are shown in **F**. **G-I.** The model predicts that dopamine biases actions that contain rewards in physical and/or conceptual proximity. Decision-dimensions important for some decisions but not all will be used to derive action values when there is high dopamine (**G**) but not when there is low dopamine (**H**). If these decision-dimensions correspond to information about location, for instance, then additional dopamine may lead to the incorporation of spatial information in decisions, leading to actions containing the same location information having more similar action values (**I**).

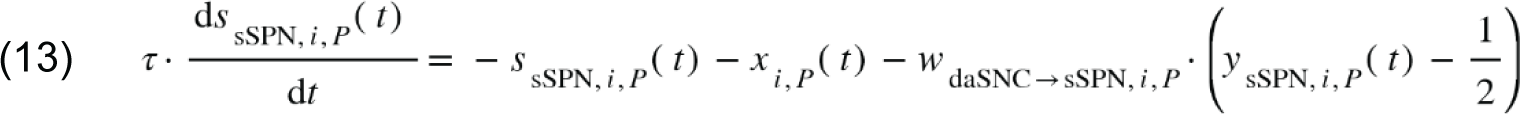

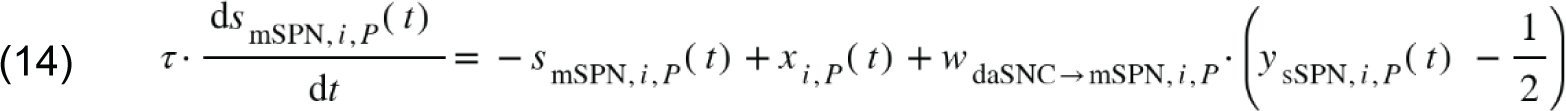

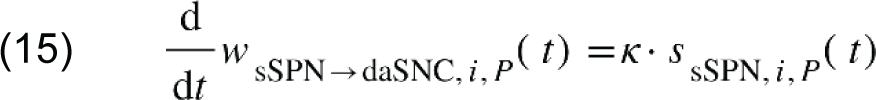

where:

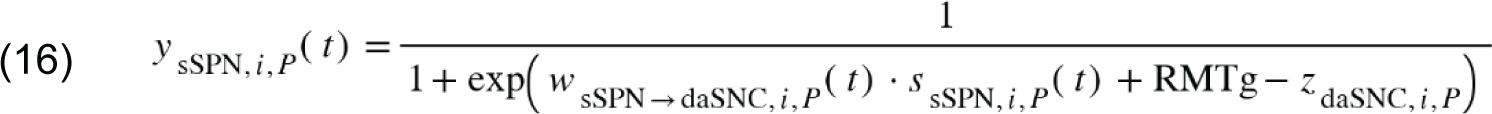

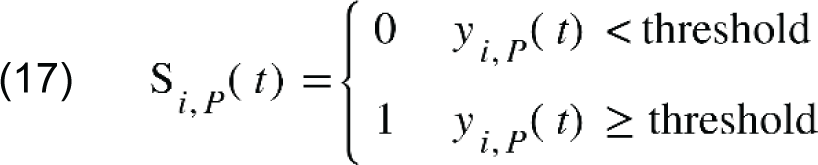

**Figure S6.**
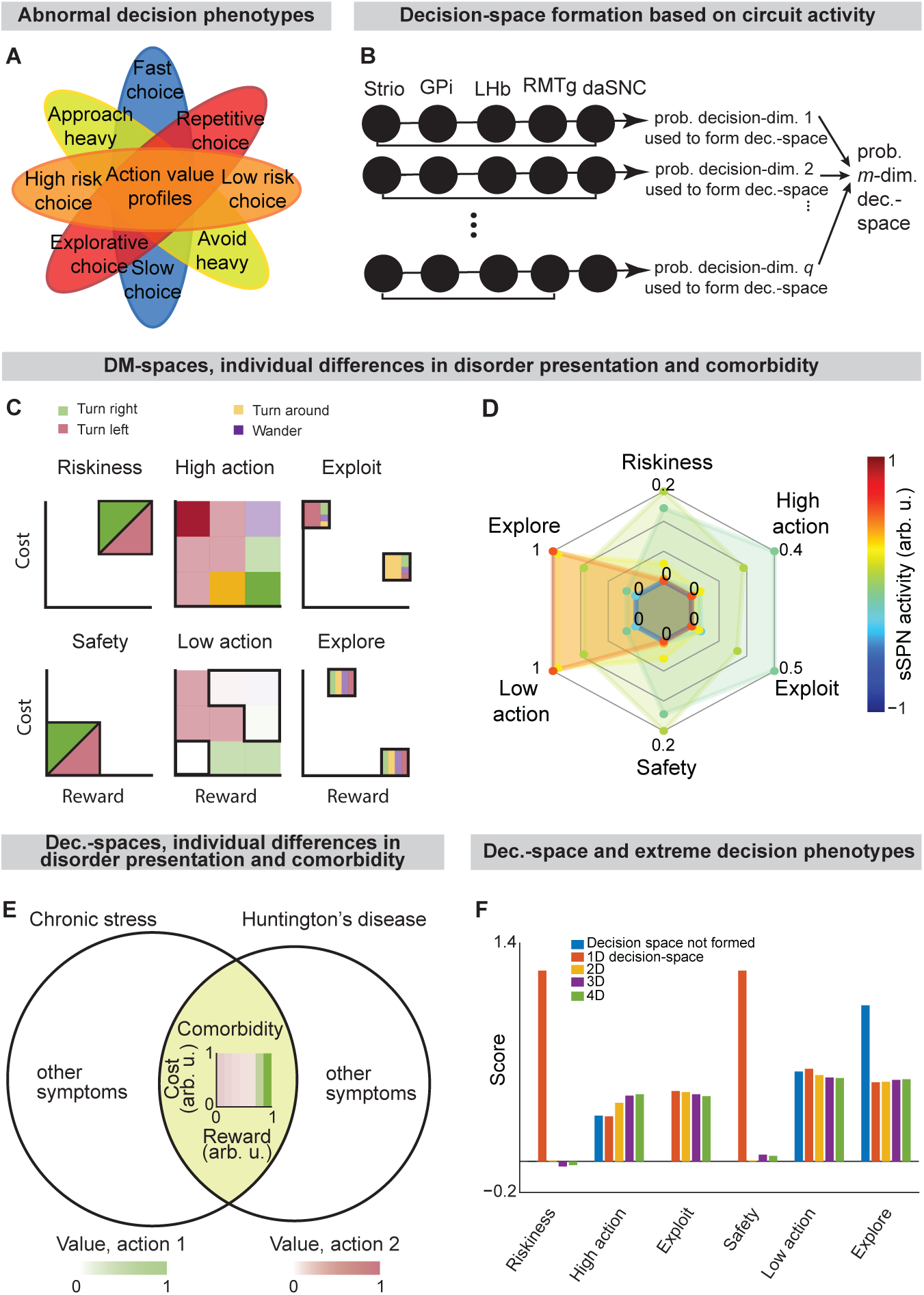
Individual decision-making differences can be explained by differences in decision-space, Related to Figure 6. (44/85) **A.** Decision-making symptoms observed in disorders. **B.** Schematic of the computational model for simulating decision-space based on circuit activity in the analyses plotted in Figures 6A**,B**. daSNC activity daSNC_1_ = daSNC_2_ = … daSPN_*q*_ = *d* is calculated for each of *q* elements, each corresponding to a decision-dimension. Decision-space dimensionality is determined from the Poisson binomial distribution of individual decision-dimension probabilities, per eq. (18). See **Defining mSPN activity and decision-space, STAR Methods**. **C.** Illustration of the method by which we score subjective valuations along six axes (riskiness, safety, high action, low action, exploit, explore) for a modeled T-maze task. A grid of action values for a range of reward and cost combinations is formed. The “riskiness” and “safety” axes are calculated based on the action values for the high-reward high-cost combinations and low-reward low-cost combinations, respectively. “High action” and “low action” axes are calculated as the proportion of the reward/cost grid with especially high (sum of all action values > 0.5) or low (< 0.2) action value, respectively. The “exploit” and “explore” axes are determined as the proportion of the reward/cost grid with especially high (> 0.5) or low (< 0.25) Gini coefficients between the action values. **D.** Extension of Figure 6C. sSPN activity is modified, leading to decision-spaces formed at different rates. These different decision-spaces lead to different action valuations. Thus, in a disorder which affects sSPN activity during decision-making, differences in decision-space may be responsible for differences in decision-making. **E.** Huntington’s disease and chronic stress both have decision-making signatures of low-dimensional decision-spaces. **F.** Extension of Figure 6F, showing that scores along the six subjective valuation axes (mean taken over 1000 simulations) are different depending on decision-space. Thus, disorders that affect decision-space formation may lead to shifts in decision-making. 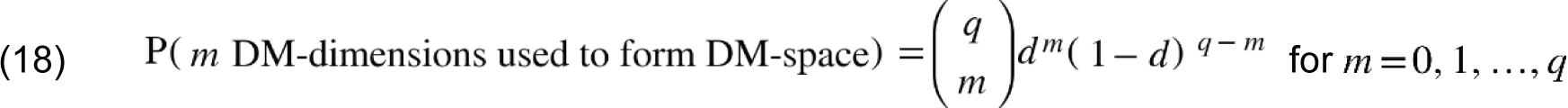

**Figure S7.**
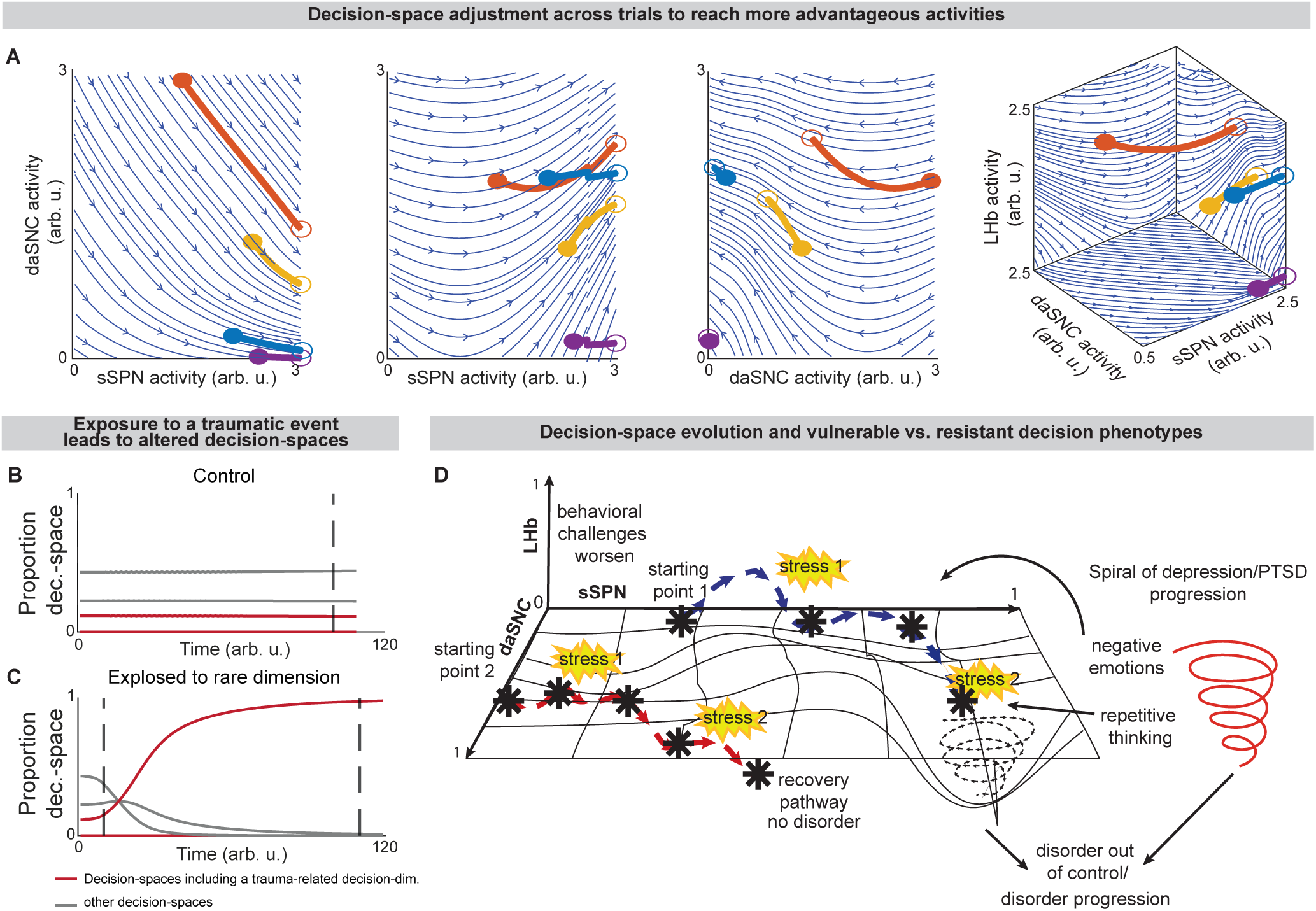
Disorder development and decision-space formation, Related to Figure 7. **A**. Simulations of trajectories of circuit activity between trials. Four trajectories of circuit activity movement (thick lines) are plotted above streamlines (blue lines) for a modeled circuit that adjusts to facilitate simple choices. Filled circles are various initial circuit activities (*t*_0_). Empty circles are the ending circuit activities (i.e. *t*_t_). The left three panels are two-dimensional slices of the plot on the right. Here and in Figure 7, the modeled circuit adjusts between trials to improve its ability to form preferred decision-spaces (advantage) while avoiding large changes to activity during decisions (cost). Advantage is represented mathematically as a weighted sum of probabilities that the possible decision-spaces form, given the circuit elements {*X*_1_,*X*_2_, … *X_n_*} take a specified set of activities *x*_1_,*x*_2_, …, *x_n_* (eq. (19)). Cost is represented as the distance between the circuit activity used during decision-making and a “baseline” circuit activity that the circuit takes outside of decision-making (eq. (20)). Net advantage is the difference between advantage and weighted cost (eq. (21)).The circuit adjusts between trials in the direction of maximal change in net advantage (eq. (22)).See ***Movement of Circuit Activity Across Multiple Trials*, STAR Methods**. **B,C**. A possible explanation for the observation that in certain psychiatric disorders (post-traumatic stress disorder and substance use disorder) exposure to a traumatic event or drug can lead to increasingly altered choice after an extended period of nonexposure or abstinence. This can be modeled as adaptation in the circuit to process a rare decision-dimension that was necessary to process, for example, the traumatic event or drug. In the simulation, the circuit’s preference for various decision-spaces is updated over time based on their success in making choices according to environmental stimuli. In the disorder resilient scenario (**B**), the circuit is first exposed to the traumatic event or drug at the dashed line. The “red” decision-space is not formed as often as are other decision-spaces (gray lines). In contrast, even short periods of exposure to the traumatic event or drug can impact future decision-spaces, resulting in vulnerability (**C**). Circuit activity adjusts until, when the traumatic event or drug reappears, the “red” decision-space forms frequently. This result may explain incubation of fear (in post-traumatic stress disorder) or craving (in a substance use disorder), where symptoms emerge only after a period of weeks after exposure. Differences in response post-incubation lead to modeled vulnerability or resilience. **D**. Circuit activity morphs over time in response to decision-making needs. 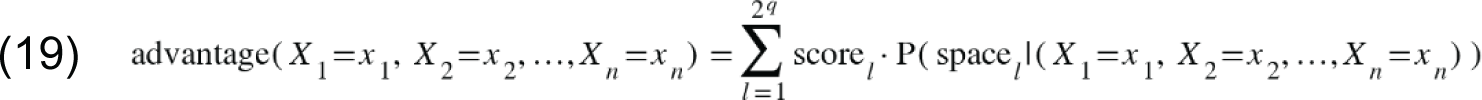

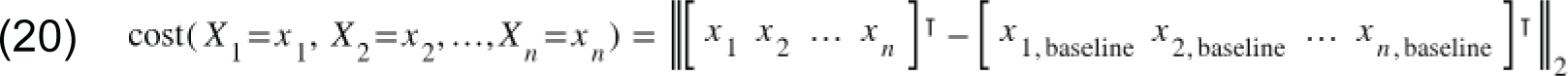

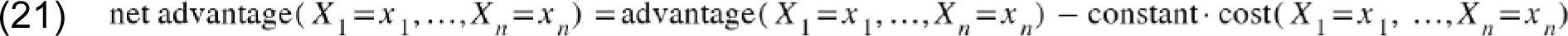

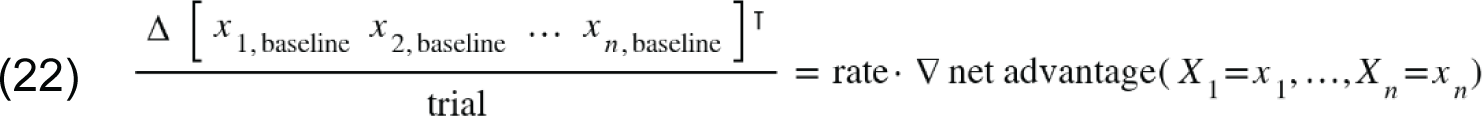

**Table S1:**
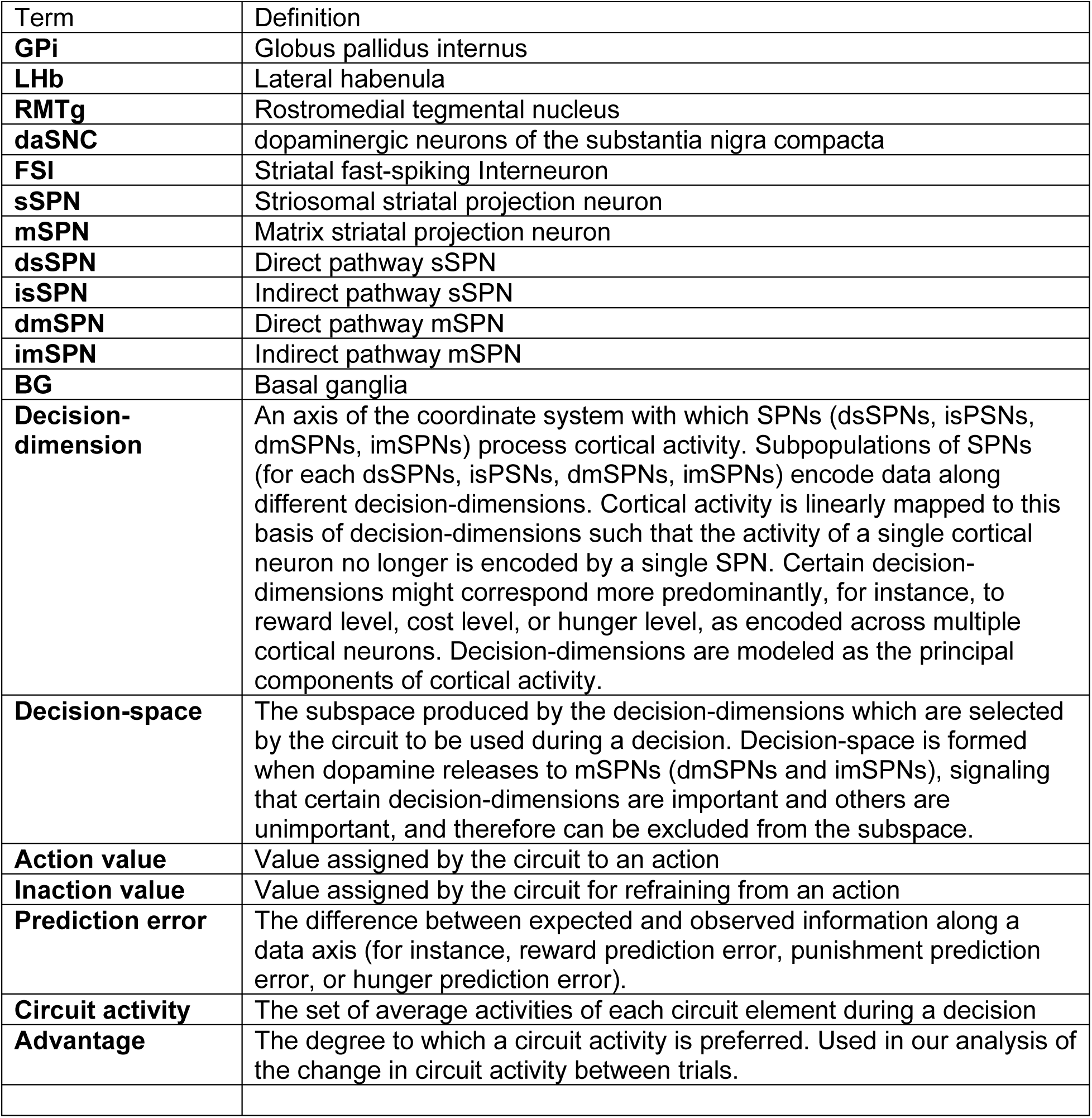
Terminology.

**Table S2:**
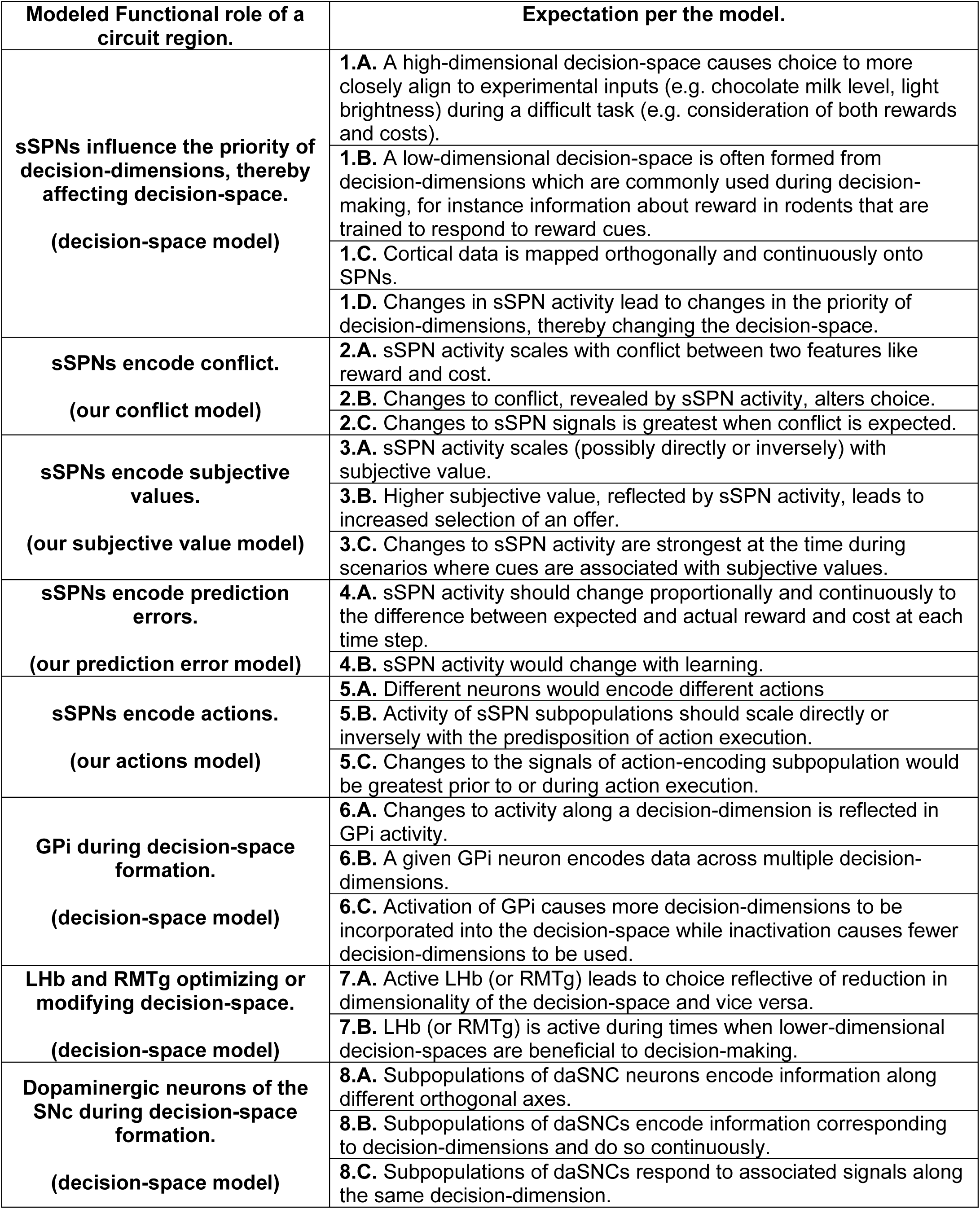
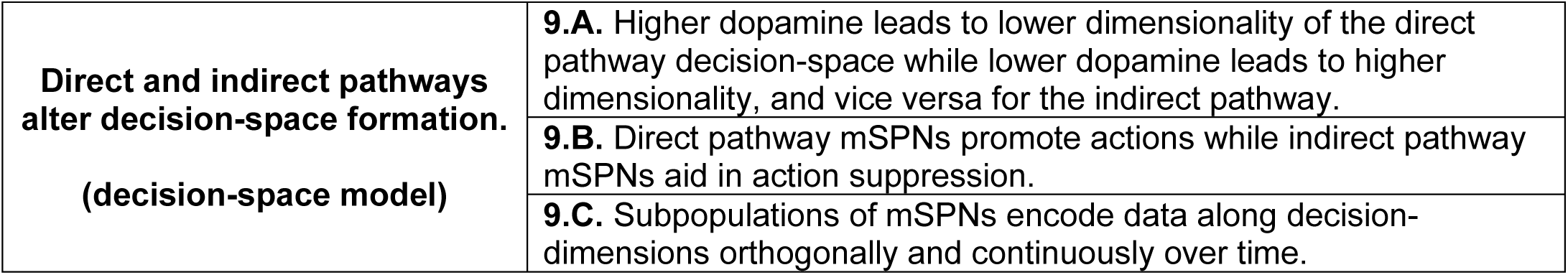
Criteria used in tables S2-7 to compare our decision-space model against other possible models of circuit function.

**Table S3:**
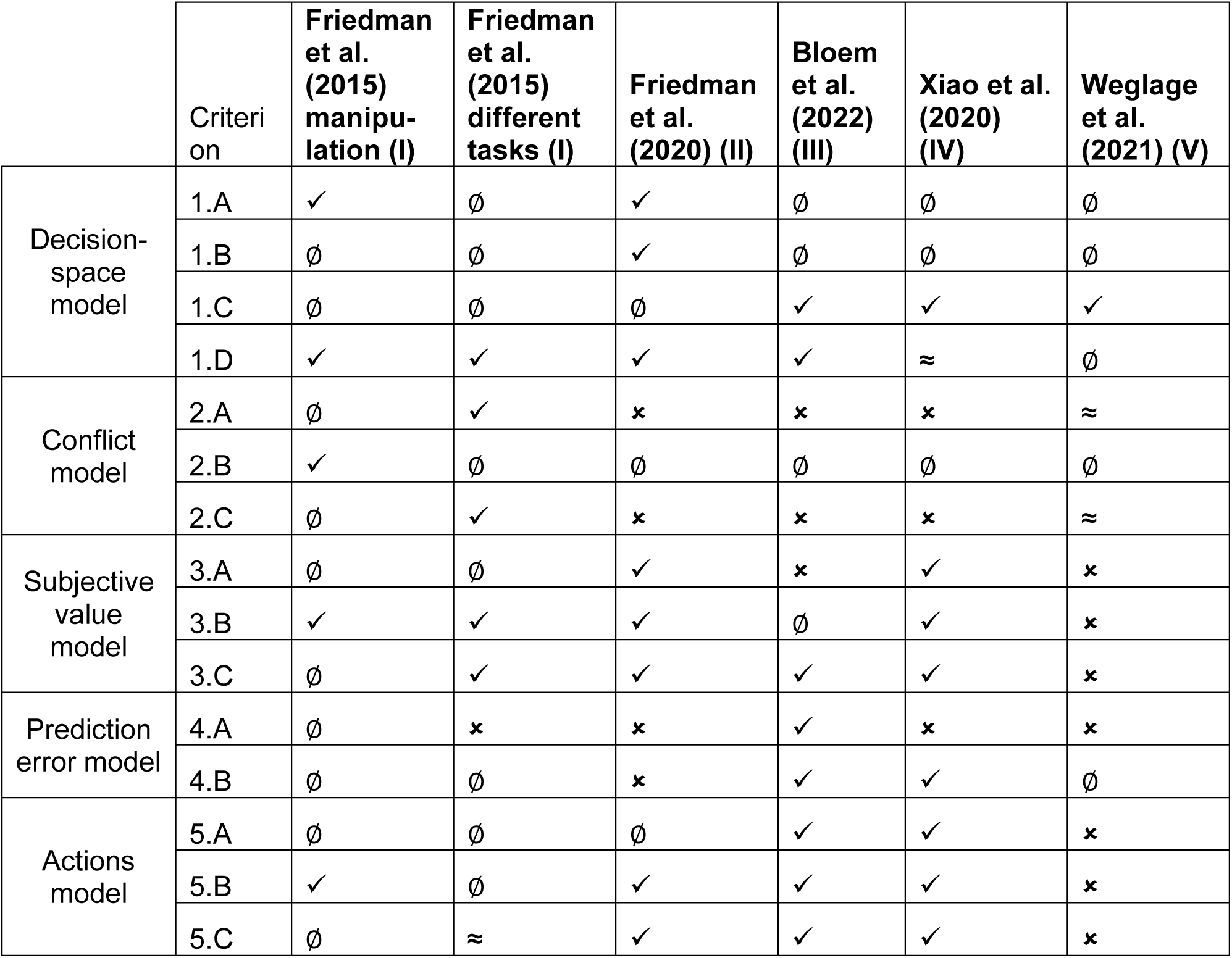
Testing alignment of the decision-space model and other models to a selection of the experimental sSPN literature. ✓ --aligned with criterion ≈ -- somewhat aligned to criterion ⌋ -- not aligned with criterion ∅ -- experiment does not test criterion **(I)** In Friedman et al. (2015), optogenetic manipulation led to less consistent choices and sSPN activity was different between tasks of different difficulty, suggesting a relationship between sSPN activity and the decision-space. **(II)** In Friedman et al. (2020), sSPN activity was different between tasks where attention to a different cue was necessary for successful completion, suggesting a relationship between sSPN activity and the decision-space. **(III)** In Bloem et al. (2022), sSPNs were found to encode reward or cost prediction errors during probabilistic cue association decision-making. This could also be interpreted through the decision-space model as a mapping of multiple cues to the same sSPN neurons which correspond to either a reward-related or a cost-related decision-dimension. **(IV)** In Xiao et al. (2020), subsets of sSPN neurons responded to reward, punishment, or both types of information. This could be interpreted through the decision-space model as subpopulations each corresponding to either a reward-related, punishment-related, or other decision-dimension. **(V)** In Weglage et al. (2021), it was shown that A) dSPN, iSPN, and sSPN activity roughly resembled one other over time in a multiphase decision-making task, and B) SPN activity was predictive of actions within each context, but not between contexts. This could be interpreted through the decision-space model as A) mapping of cortical activity to a basis of decision-dimensions somewhat similarly in sSPNs versus mSPNs when the decision-space is high-dimensional, and B) SPNs encoding data along decision-dimensions (which would be different between tasks) rather than actions.

**Table S4:** Testing the alignment of the decision-space model to a selection of the experimental literature on GPi. ✓ --aligned with criterion ≈ -- somewhat aligned to criterion ⌋ -- not aligned with criterion ∅ -- experiment does not test criterion **(I)** In Weglage et al. (2022), it was found that a specific subpopulation of LHb-projecting GPi neurons is influential to exploration and choice selection. Through the lens of the decision-space model, this could be interpreted as a GPi→LHb connection that facilitates switching between a high-versus low-dimensional decision-space. **(II)** In Munte et al. (2017), it was found that the GPi responds to reward signals in patients undergoing deep brain stimulation treatment. This could be interpreted through the decision-space model as adjustment of GPi activity based on its input from an sSPN subpopulation corresponding to a reward-related decision-dimension. **(D)** In Stephenson-Jones et al. (2016), LHb-projecting GPi neurons were excited by punishment predicting cues and the punishment itself but were inhibited by rewards and their associated cues. This could be interpreted through the lens of the decision-space model as adjustment of GPi activity based on its input from two sSPN subpopulations, one corresponding to a reward-related decision-dimension and the other to a cost-related decision-dimension.

| Criterion | Weglage et al. (2022)<br>(I) | Munte et al.<br>(2017) (II) | Stephenson-Jones et. al<br>(2016) (III) |
| --- | --- | --- | --- |
| 6.A | ∅ | ✓ | ∅ |
| 6.B | ∅ | ∅ | ✓ |
| 6.C | ✓ | ∅ | ✓ |

**Table S5:** Testing the alignment of the decision-space model to a selection of the experimental literature on LHb and RMTg. ✓ -- aligned with criterion ≈ -- somewhat aligned to criterion ⌋ -- not aligned with criterion ∅ -- experiment does not test criterion **(I)** In Matsumoto & Hikosaka (2007), it was found that LHb activation leads to suppression of dopaminergic signaling among daSNC neurons, lending support to the decision-space model. **(II)** In Lee & Hikosaka (2022), LHb was found to alter its activity depending on situational context. This could be interpreted, through the lens of the decision-space model, as a shift towards a higher- or lower-dimensional decision-space based on context. **(III)** In Stopper & Floresco (2014), LHb inactivation led subjects to change their choice during a probabilistic discounting task to accept a large, risky reward over a smaller, safe reward. This lends support to the functional role of LHb in the decision-space model which is to influence the types of informational input used during a decision. **(IV)** In Vento et al. (2017), RMTg selectively altered decisions, primarily in response to cost. This could be interpreted, through the lens of the decision-space model, as RMTg influencing the decision-space when it receives contextual inputs.

| Criterion | Matsumoto & Hikosaka (2007) (I) | Lee & Hikosaka (2022) (II) | Stopper & Floresco (2014) (III) | Vento et al. (2017) (IV) |
| --- | --- | --- | --- | --- |
| 7.A | ∅ | ∅ | ≈ | ✓ |
| 7.B | ✓ | ✓ | ∅ | ∅ |

**Table S6:** Testing the alignment of the decision-space model to a selection of the experimental literature on daSNC. ✓ -- aligned with criterion ≈ -- somewhat aligned to criterion ⌋ -- not aligned with criterion ∅ -- experiment does not test criterion **(I)** In Fiorillo et al. (2003), daSNC neurons responded differently depending on the likelihood of a cue predicting a reward outcome. Through the lens of the decision-space model, this supports the mapping of cue-related information about reward, for instance, to a reward-predominant decision-dimension, which then impacts dopamine release. **(II)** In Matsumoto & Hikosaka (2009), two dopamine populations responded very differently to rewarding or aversive stimuli and a third group was non-responsive. This supports the tenet of the decision-space model that different daSNC subpopulations correspond to different decision-dimensions, some of which might be related to reward information, some to cost information, and some to neither reward nor cost information. **(III)** In Gan et al. (2010), it was found that the activities of recorded daSNC neurons showed more resemblance to reward levels than to overall utility. This supports the tenet of the decision-space model that different daSNC subpopulations correspond to different decision-dimensions, some of which are related to reward information, and that daSNC neurons encode data along decision-dimensions, not an overall value function. **(IV)** In Bromberg-Martin et al. (2010), subpopulations of dopamine neurons that encoded value were excited by rewarding information while salience neurons were excited by both rewarding and aversive cues. This supports the tenet of the decision-space model that different daSNC subpopulations correspond to different decision-dimensions, some of which are related to reward information and some to other information. **(V)** In Kim et al. 2020, it was found that dopamine changes in response to altered proximity to reward. This supports the architecture of the decision-space model, where changes to reward information are captured in an sSPN subpopulation related to reward, then passed to a corresponding daSNC subpopulation.

| Criterion | Fiorillo et al. (2003) (I) | Matsumoto & Hikosaka (2009) (II) | Gan et al. (2010) (III) | Bromberg-Martin et al. (2010) (IV) | Kim et al. (2020) (V) |
| --- | --- | --- | --- | --- | --- |
| 8.A | ✓ | ✓ | ✓ | ✓ | ✓ |
| 8.B | ✓ | ✓ | ✓ | ✓ | ✓ |
| 8.C | ✓ | ✓ | ✓ | ∅ | ∅ |

**Table S7:** Testing the alignment of the decision-space model to a selection of the experimental literature on the direct versus indirect pathways (A). ✓ -- aligned with criterion ≈ -- somewhat aligned to criterion ⌋ -- not aligned with criterion ∅ -- experiment does not test criterion **(I)** In Parker et al. (2018), it was found that the direct and indirect pathway typically are coactivated, and dopamine depletion differentially alters direct versus indirect pathway dynamics. This supports the decision-space model, in which two decision-spaces form during a decision, one in each pathway, and are affected differently by dopamine. **(II)** Peak et al. (2020) demonstrated that inhibition of dSPNs during learning leads to blunted action associations, while inhibition of iSPNs leads to a reduced ability to switch actions based on context. This supports the decision-space model, in which the direct pathway is involved with performing actions and the indirect pathway is involved in refraining from actions. **(III)** In Maltese et al. (2021), increased dopamine release led to increased activation of dSPNs and decreased activation of iSPNs, and decreased dopamine release led to the opposite change. This supports the decision-space model, in which dopamine excites dmSPNs and inhibits imSPNs. **(IV)** Barbera et al. (2016) found that striatal neurons fire in clusters, supporting the tenet of the decision-space model that proximate clusters of sSPNs encode data along decision-dimensions.

| Criterion | Parker et al.<br>(2018) (I) | Peak et al.<br>(2020) (II) | Maltese et al.<br>(2021) (III) | Barbera et al.<br>(2016) (IV) |
| --- | --- | --- | --- | --- |
| 9.A | ✓ | ∅ | ✓ | ✓ |
| 9.B | ∅ | ✓ | ∅ | ✓ |
| 9.C | ∅ | ∅ | ∅ | ✓ |

