## Supplementary material for "Model of a striatal circuit exploring biological mechanisms underlying decision-making during normal and disordered states": Methods

### RESOURCE AVAILABILITY

#### *Lead Contact*

#### *Materials availability*

Code used to construct, analyze, and test the model is deposited to https://github.com/dirkbeck/DM_space_model.

Code used to analyze neural data from the Corticostriosomal Circuit Stress Experiment database is deposited to https://github.com/dirkbeck/DM_space_model.

Data from the Corticostriosomal Circuit Stress Experiment data, prepared for use in the current paper, is deposited to https://doi.org/10.7910/DVN/SMKW0I.

#### *Data and code availability*

- This paper analyzes existing, publicly available data. These accession numbers for the datasets are listed in the key resources table.
- All original code has been deposited at https://github.com/dirkbeck/DM_space_model and is publicly available as of the date of publication. DOIs are listed in the key resources table.
- Any additional information required to reanalyze the data reported in this paper is available from the lead contact upon request.

### METHOD DETAILS

#### *Outline.*

- ***Decision-dimensions and decision-space.*** Explanation of the foundational concept of the model.
- ***Analyzed instances of the model.*** We conduct our analysis using three instances of the conceptual model. In the following sections, we formally define the model in each instance, then detail the methods behind our related analyses.
    - ***Instance 1: full connectivity and feedforward.*** Related to **Figures 1-4,6, S1-3,6**. Used to link neural activity, the decision-space, and choice.
    - ***Modeled Circuit Manipulation using Instance 1.***
    - ***Instance 2: sparse connectivity and feedforward.*** Related to **Figure S4**. Used to demonstrate how a large network might encode the decision-space.
    - ***Modeling SPN encoding of data, using Instance 2.***
    - ***Instance 3: full connectivity and dynamics.*** Related to **Figure 5, S5**. Used to demonstrate how the decision-space might form over time.
    - ***Modeling time-variant input, using Instance 3.***

- ***Movement of Circuit Activity Across Multiple Trials.*** An extension of the model to view possible changes of the circuit between trials in the context of decision-space. Related to **Figure 7, S7**.
- ***Rationale for the computational framework.*** Reasoning behind our modeling strategies.
- ***Inferring decision-space from SPN activity and choice.*** A method we designed in which decision-space can be inferred from experimental data. Related to **Figure S1Q,R**.
- ***Testing the Model Through Analysis of Neural Data.*** Analysis of neural data which supports our model. Related to **Figures 2F,G, S2A-N,S3A-P.**

***Decision-dimensions and decision-space.***

The physiologies of the circuit elements produce two abstractions which we use, for convenience, throughout or work:

- A *decision-dimension* is an axis of the coordinate system with which SPNs (dsSPNs, isSPNs, dmSPNs, imSPNs; see **Table S1** for anatomical definitions) encode data projected from the cortex. A decision-dimension is equivalent to a principal component of cortical activity. In our analysis, separate groups of SPNs encode data along the first, second, third, and fourth principal components. (Arbitrarily, we do not consider principal components beyond the first four). Each of the dsSPN/isSPN/dmSPN/imSPN subgroups have neurons corresponding to each of the four principal components.
- The *decision-space* is the mathematical space formed by mSPNs (both dmSPNs and imSPNs) after dopamine signaling from daSNC. Modeled dopamine signaling determines whether to include or exclude neurons encoding each decision-dimension during a decision. Thus, from the mathematical space formed by all decision-dimensions, a mathematical subspace (i.e. the “decision-space”) is formed with which to define action values.

We use the prefix “decision” because the circuit uses decision-space, formed from a basis of decision-dimensions, to define action values during decision-making.

***Analyzed Instances of the Model.***

The general case of the model (although not formally used for analysis) is a dynamic network of cortical neurons, FSIs, dsSPNs, isSPNs, GPi, LHb, daSNCs, dmSPNs, imSPNs, and mSPN-projecting neurons which encode action values.

We conduct our analyses using three instances of this general case, which are each equivalent to the general case under the specific conditions we outline. The three instances, tailored to our various analyses, each allow for a different mathematical simplification. This allows us to conceptually and formally define the instances individually in a way that is intuitive and relates directly to our analyses.

1) Instance 1 has full cortex→FSI→SPN connectivity and constant activity in each circuit element throughout the decision. In this instance, the model can be defined equivalently

using a smaller set of network elements and a feedforward network. See ***Instance 1: full connectivity and feedforward.***

2) Instance 2 has constant activity in each circuit element throughout the decision. In this instance, the model can be defined equivalently using a feedforward network. See ***Instance 2: sparse connectivity and feedforward.***
3) Instance 3 has full cortex→FSI→SPN connectivity. In this instance, the model can be defined equivalently using a smaller set of network elements. See ***Instance 3: full connectivity and dynamics.***

#### ***Instance 1: full connectivity and feedforward.***

In this section, we describe the instance of the model where each cortical neuron projects to each FSI, each FSI projects to each SPN (for dsSPN, isSPN, dmSPN, imSPN), and each cortical neuron projects to each SPN. Additionally, cortex input to the system does not change over time, and the activities of other circuit elements do not decay over time.

This instance of the model leads to a convenient formation of the model as a circuit of fewer elements (one FSI, one dsSPN, isSPN, dmSPN, and imSPN per the four decision-dimensions), and no time component. In this section, we frame this instance mathematically and then describe our related analysis.

##### **Input: cortical activity.**

During a decision, a vector of cortical input $\mathbf{x}_P \in \mathbb{R}^{p \times 1}$ enters each pathway $P$ in the network ( $\mathbf{x}_{\text{direct}}$ to direct pathway SPNs and $\mathbf{x}_{\text{indirect}}$ to indirect pathway SPNs). The elements of $\mathbf{x}_P$ are the activities of $p$ cortical neurons. Each neuron encodes a different sensory input.

##### **Outputs.**

We use this instance of the model to examine: 1) the activities of the circuit elements depending on the activities of other circuit elements (**Figures 1A-C, S1A-I**); 2) the activation/inactivation of mSPNs by dopamine (i.e. decision-space, **Figures 1D-I**); 3) action values given decision-space (**Figures S1J-L**), and 4) choice given action values (**Figures S1M-P**).

1) The activities of circuit elements during a decision are related to each other based on anatomically realistic connections (eqs. (1),(2),(3),(5),(23)).
2) A decision-space is formed probabilistically. The probability a given decision-dimension $i$ being used during a decision is equivalent to the activity of daSNC (see eq. (2)), which ranges from 0 to 1. Probabilities are realized in the connection from daSNC to mSPN (see eq. (3)), when each decision-dimension is probabilistically assigned a weight (in most analyses, either 0 or 1). Decision-space is defined as the space formed from the basis of decision-dimensions that were not assigned a weight of 0.

3) Action value is derived based on mSPN activity during a decision.
4) Choice is derived from action values. Action values are treated as Merton processes[214] using eq. (6). Several possible actions are assigned action values and the corresponding process that hits the threshold first is enacted.

**Defining FSI activity.**

FSI activity, $c_P$, is set to the magnitude of $\mathbf{x}_P$ for each pathway, multiplied by a weight of cortex→FSI connection $a_{\mathrm{FSI}}$, plus an additive shift $b_{\mathrm{FSI}}$:

(23) $$c_P = a_{FSI} \cdot \left\| \mathbf{x}_P \right\| + b_{FSI}$$

where:

- $c_P$ is relative activities of FSIs that project to SPNs of pathway $P$ (activity arb. u.)
- $a_{\mathrm{FSI}}$ is the weight of cortex→FSI connection. Similar for both $P$. (dimensionless)
- $\mathbf{x}_P$ is the activities of cortical neurons that project to SPNs of pathway $P$. (activity arb. u.)
- $b_{\mathrm{FSI}}$ affects the relative activity of all sSPN neurons. Similar for both $P$. (activity arb. u.)

In the current instance of the model, there are 2 FSIs, one that receives input from $\mathbf{x}_{\mathrm{direct}}$ and projects to dsSPNs and dmSPNs, and the other than receives input from $\mathbf{x}_{\mathrm{indirect}}$ and projects to isSPNs and imSPNs.

For use in our analysis, see https://github.com/dirkbeck/DM_space_model/blob/main/algorithmic_model.m.

**Defining sSPN activity.**

To get the activities of sSPNs in each pathway, $\mathbf{x}_P$ is normalized via division by $c_P$ and multiplied by $\mathbf{W}_P \in \mathbb{R}^{p \times q}$, which linearly transforms and reduces cortical input from the $p$-dimensional coordinate space of cortex to the smaller $q$-dimensional coordinate space of sSPN. In the sSPN coordinate space, each coordinate is a principal component of a training set of historical cortical input across $n$ time steps $\mathbf{X}_P \in \mathbb{R}^{n \times p}$ (uncorrelated, for simplicity). For each pathway, $\mathbf{W}_P$ contains the truncated first $q$ columns (corresponding to the first $q$ principal components) of $\mathbf{W}_{\mathrm{full},P} \in \mathbb{R}^{p \times p}$ after the decomposition $\mathbf{X}_P \mathbf{X}_P^{\top} = \mathbf{W}_{\mathrm{full},P} \mathbf{\Lambda} \mathbf{W}_{\mathrm{full},P}^{\top}$ is made to obtain the full principal component matrix. Experimental work has revealed dimensionality reduction the order of ~100 times from cortex to SPNs[114], so $q \ll p$. Note that in our analysis using the current instance of our model, $\mathbf{X}_P$

is not explicitly generated because we specify the inputs to the system in terms of the coordinate space of decision-dimensions.

For each pathway, the components of an sSPN activity vector $\mathbf{s}_{\mathrm{sSPN},P} \in \mathbb{R}^{q \times 1}$ each correspond to the activity of an sSPN circuit element. A constant $b_{\mathrm{sSPN}}$, used in analyses where modeled sSPN activity is stimulated or inhibited, adjusts overall sSPN activity:

(1) $$\mathbf{s}_{\mathrm{sSPN},P} = \frac{1}{c_P}\mathbf{W}_P^{\mathrm{T}}\mathbf{x}_P + b_{\mathrm{sSPN}}$$ (copied from **Results** for convenience)

where:

- $c_P$ is the relative activity of the FSI projecting to SPNs of pathway $P$ (activity arb. u.)
- $\mathbf{W}_P$ is a matrix of weights from cortical neurons to SPNs of pathway $P$. Each column is equivalent to a principal component of cortical activity. (dimensionless)
- $\mathbf{x}_P$ is the activities of cortical neurons that project to SPNs of pathway $P$. (activity arb. u.)
- $b_{\mathrm{sSPN}}$ affects the relative activity of all sSPN neurons (activity arb. u.)

In the current instance of the model, activities are defined based on a feedforward network, so the simplification is made that sSPN activities are not affected by daSNC activities.

In the current instance, there is one sSPN per decision-dimension per pathway. So, there are $q$ dsSPNs and $q$ isSPNs. The dsSPNs receive input from $\mathbf{x}_{\mathrm{direct}}$ and $c_{\mathrm{direct}}$. The isSPNs receive input from $\mathbf{x}_{\mathrm{indirect}}$ and $c_{\mathrm{indirect}}$.

For use in our analysis, see https://github.com/dirkbeck/DM_space_model/blob/main/algorithmic_model.m.

**Defining GPi, LHb, and RMTg activities.**

The GPi→LHb→RMTg→daSNC pathway performs a series of operations which influence RMTg activity $\mathrm{RMTg}$, which is an input to daSNC activity in eq. (2). Weights $\mathbf{w}_{\mathrm{GPi},P} \in \mathbb{R}^{2q \times 1}$, not necessarily positive, are combined with the activities of the $q$ dsSPNs and $q$ isSPNs, forming scalar representations of dsSPN or isSPN activity. $z_{\mathrm{GPi}}$, $z_{\mathrm{LHb}}$, and $z_{\mathrm{RMTg}}$ terms reflecting the activities of those circuit elements are combined with this scalar representation:

(5) $$\mathrm{RMTg} = z_{\mathrm{RMTg}} + z_{\mathrm{LHb}} + z_{\mathrm{GPi}} \cdot \mathbf{w}_{\mathrm{GPi}} \cdot \begin{bmatrix} \mathbf{s}_{\mathrm{sSPN,direct}} \\ \mathbf{s}_{\mathrm{sSPN,indirect}} \end{bmatrix}$$ (copied from **Figure S1** for convenience)

where:

- $z_{\mathrm{RMTg}}$ is an additive shift that affects relative RMTg activity (activity arb. u.)
- $z_{\mathrm{LHb}}$ is an additive shift that affects relative LHb activity (activity arb. u.)
- $z_{\mathrm{GPi}}$ is a coefficient that affects relative GPi activity (activity arb. u.)
- $\mathbf{w}_{\mathrm{GPi}}$ is the weights of connection from sSPNs of pathway $P$ to the GPi neuron (dimensionless)
- $\mathbf{s}_{\mathbf{sSPN},P}$ is the activity of sSPNs corresponding to decision-dimension $i$ and pathway $P$ (activity arb. u.)

This pathway contains one GPi element, one LHb element, and one RMTg element, as visualized in **Figure 1A**. All sSPN elements project to GPi. GPi activity is an input to LHb, which after the $z_{\mathrm{LHb}}$ addition, is an input to RMTg, which itself has a $z_{\mathrm{RMTg}}$ addition. For simplicity, these series of operations are presented together in eq. (5).

For use in our analysis, see https://github.com/dirkbeck/DM_space_model/blob/main/algorithmic_model.m.

**Defining daSNC activity.**

daSNC neurons incorporate the output of the GPi→LHb→RMTg→daSNC pathway with direct inputs from sSPN elements. There are $q$ sSPN elements of each pathway and $q$ daSNC elements corresponding to each pathway. For each pathway, the $i$ th sSPN element connects to the $i$ th daSNC element, but not to other daSNC elements (see **Figure 1A**). These connections have weights $w_{\mathrm{sSPN}\to\mathrm{daSNC},i,P}$ for $i = 1, 2, \ldots, q$. RMTg, on the other hand, connects to each daSNC element. The output of the $i$ th daSNC element, constrained to between 0 and 1 via a logistic function, captures the importance of a single decision-dimension:

(2) $$\mathrm{daSNC}_{i,P} = \frac{1}{1 + \exp\left( w_{\mathrm{sSPN}\to\mathrm{daSNC},i,P} \cdot \mathrm{s}_{\mathbf{sSPN},i,P} + \mathrm{RMTg} - z_{\mathrm{daSNC},i,P} \right)}$$

(copied from **Results** for convenience)

where:

- $w_{\mathrm{sSPN}\to\mathrm{daSNC},i,P}$ is the weight of connection from the sSPN corresponding to decision-dimension $i$ and pathway $P$ to the daSNC corresponding to decision-dimension $i$ and pathway $P$ . The weight is fixed in this instance of the model. (dimensionless)

- $\mathbf{s}_{\mathrm{sSPN}, i, P}$ is the activity of sSPNs corresponding to decision-dimension $i$ and pathway $P$ (activity arb. u.)
- $z_{\mathrm{daSNC}, i, P}$ is an additive shift applied to the daSNC neuron corresponding to decision-dimension $i$ and pathway $P$ (activity arb. u.)
- $\mathrm{RMTg}$ is RMTg activity, as defined in eq. (5). (activity arb. u.)

This pathway is modeled using one neuron per decision-dimension per pathway. So, there are $q$ daSNC neurons that each receive projection from a dsSPN, and $q$ daSNC neurons that each receive projection from an isSPN. daSNC elements also receive input from RMTg.

For use in our analysis, see https://github.com/dirkbeck/DM_space_model/blob/main/algorithmic_model.m.

**Defining mSPN activity and decision-space.**

In each pathway, decision-space is formed probabilistically. The conversion from daSNC activity to realization of decision-space occurs in the connections from daSNC to mSPN. There are $q$ daSNC elements corresponding to each pathway and $q$ mSPN elements, and, in each pathway, the $i$ th daSNC element connects to the $i$ th mSPN element, but not to other mSPN elements (see **Figure 1A**).

Like sSPNs, mSPNs encodes the cortical input normalized by an FSI and is transformed to a coordinate space of the first $q$ principal components. The difference is that for each of dmSPNs and imSPNs, a diagonal matrix $\mathbf{S}_P \in \mathbb{R}^{q \times q}$ is multiplied by the cortical input after transformation:

(3) $$\mathbf{s}_{\mathbf{mSPN}, P} = \frac{1}{c_P} \mathbf{S}_P \mathbf{W}_P^{\mathrm{T}} \mathbf{x}_P$$ (copied from **Results** for convenience)

where:

- $c_P$ is the relative activity of the FSI projecting to SPNs of pathway $P$ (activity arb. u.)
- $\mathbf{S}_P$ is a diagonal matrix that applies dopamine release (via daSNC activity) to mSPN activity in pathway $P$. (dimensionless)
- $\mathbf{W}_P$ is a matrix of weights from cortical neurons to SPNs of pathway $P$. Each column is equivalent to a principal component of cortical activity. (dimensionless)
- $\mathbf{x}_P$ is the activities of cortical neurons that project to SPNs of pathway $P$. (activity arb. u.)

In the current instance, there is one mSPN per decision-dimension per pathway. So, there are $q$ dmSPNs and $q$ imSPNs. The dmSPNs receive input from $\mathbf{x}_{\mathrm{direct}}$ and $c_{\mathrm{direct}}$. The imSPNs receive input from $\mathbf{x}_{\mathrm{indirect}}$ and $c_{\mathrm{indirect}}$.

The diagonal elements of $\mathbf{S}_P$ are set probabilistically to either 1 (dimension in decision-space) or 0 (dimension not in decision-space) such that $\mathrm{P}\left(\mathbf{S}_{\mathrm{P,ii}}=1\right)=\mathrm{daSNC}_{i,\mathrm{P}}$.

Thus, in the portions of our analysis where we set the activities of the $q$ daSNC elements to be equal, the decision-dimensions each have the same probability of being included in decision-space, i.e. $\mathrm{daSNC}_1=\mathrm{daSNC}_2=\ldots=\mathrm{daSPN}_q=d$. In this case, we treat the probability of a certain decision-space dimensionality forming as a binomial distribution:

(18) $\mathrm{P}(m \text{ DM-dimensions used to form DM-space}) = \binom{q}{m} d^m(1-d)^{q-m}$ for $m=0,1,\ldots,q$

(copied from **Figure S6** for convenience)

where:

- $q$ is the number of possible decision-dimensions
- $d$ is the (equal) probability that each decision-dimension is used to form decision-space

**Defining action value.**

Action value (or, in the indirect pathway, inaction value) $v_{j,P}$ for each of $k$ potential actions is defined based on the activities of dmSPNs (or imSPNs). During this process, elements of a coefficient matrix $\boldsymbol{\beta}_P \in \mathbb{R}^{k\times q}$ are applied to mSPN activities for each decision-dimension, action, and pathway. Bias $\alpha_{j,P}$ is subtracted. Below, $\boldsymbol{\beta}_{j,P}$ is used to indicate row $j$ of $\boldsymbol{\beta}_P$.

(4) $v_{j,P}=\dfrac{1}{1+\exp\left(-\boldsymbol{\beta}_{j,P}\,\mathbf{s}_{\mathrm{mSPN},P}-\alpha_{j,P}\right)}$ (copied from **Results** for convenience)

where:

- $\boldsymbol{\beta}_{j,P}$ is a matrix of weights from dmSPNs to downstream action value encoding neurons for the direct pathway, or imSPNs to downstream inaction value encoding neurons for the indirect pathway. (dimensionless)
- $\mathbf{s}_{\mathrm{mSPN},P}$ is the activity of sSPNs corresponding to decision-dimension $i$ and pathway $P$ (activity arb. u.)
- $\alpha_{j,P}$ is an additive shift corresponding to the neuron encoding action $j$ for the direct pathway or inaction $j$ for the indirect pathway. (activity arb. u.)

There is one neuron encoding each $v_{j,P}$. So, there are $k$ neurons encoding action values and $k$ neurons encoding inaction values. Each of these neurons receives projection from all mSPNs of the corresponding pathway.

**Defining choice.**

$k$ Merton process[214] are run to determine whether each action should be taken, and another $k$ to determine whether each action should be refrained from. Progress to choice for each action (or inaction), $Y_{j,P}$, is related to its corresponding action (or inaction) value $v_{j,P}$ and an uncorrelated Brownian component $\mathrm{d}W_{j,P}$ scaled by a coefficient $\sigma$.

(6) $$\mathrm{d}Y_{j,P} = v_{j,P}\,\mathrm{d}t + \sigma\,\mathrm{d}W_{j,P}\ ,\ Y_{j,P}(t=0) = 0,\ \text{where } W_{j,P} \text{ is a standard Wiener process}$$

(copied from **Figure S1** for convenience)

where:

- $Y_{j,P}$ is the progress to enaction of action $j$ in the direct pathway, and progress to refraining from action $j$ in the indirect pathway.
- $v_{j,P}$ is the action (or inaction) value corresponding to action $j$ and pathway $P$. (activity arb. u.)
- $\sigma$ is the coefficient of noise.

The time it would take to enact action $j$, $t_{\mathrm{action},j}$, is defined as the first hit time of a threshold $h$ for process $j$ of the direct pathway:

(7) $$t_{\mathrm{action},j} = \min_t \left\{ t \mid Y_{j,\mathrm{direct}} \geq h \right\}$$ (copied from **Figure S1** for convenience)

The time it takes to exclude action $j$ from consideration, $t_{\mathrm{inaction},j}$, is calculated similarly using the indirect pathway:

(8) $$t_{\mathrm{inaction},j} = \min_t \left\{ t \mid Y_{j,\mathrm{indirect}} \geq h \right\}$$ (copied from **Figure S1** for convenience)

The enacted action is the first to reach $h$, given that the corresponding inaction process has not first reached $h$:

(9) $\text{action} = \arg\min_{j \in J}\left( Y_j(t_{\text{action},j}) \right)$, where $J$ is the subset of actions s.t. $t_{\text{action},j} < t_{\text{inaction},j}$

(copied from **Figure S1** for convenience)

where:

- $Y_{j,P}$ is the progress to enaction of action $j$ in the direct pathway, and progress to refraining from action $j$ in the indirect pathway.
- $t$ is time (s)
- $h$ is a threshold at which an action is considered taken (progress to decision arb. u.)

In our analysis, we run simulations using a constant time step discretization of eq. (6).

For code, see https://github.com/dirkbeck/DM_space_model/blob/main/weiner_process_model.m.

***Modeled Circuit Manipulation using Instance 1.***

To get a sense of the functional role of the circuit elements, we conducted sensitivity analyses by changing parameters in the model individually and determining their effect on the activities of other circuit elements, decision-space formation, action values, and/or choice.

**Common parameters.**

The values specified here, arbitrarily chosen, are used in the analyses in ***Instance 1*** unless otherwise indicated:

- throughout, $k = 4$
- in eq. (23): $a_{\text{FSI}} = 1$
- in eq. (23): $b_{\text{FSI}} = 0.5$
- in eq. (1): $b_{\text{sSPN}} = 0$
- in the inputs to eq., $q = 4$
- in eq. (5): $z_{\text{GPi}} = 1$
- in eq. (5): $z_{\text{LHb}} = 0.5$
- in eq. (5): $z_{\text{RMTg}} = 0.5$
- in eq. (2): $w_{\text{sSPN} \rightarrow \text{daSNC}, i, P} = 1$ for all $i$ and $P$

- in eq. (2): $z_{\text{daSNC},i,P}$ for all $i$ and $P$
- in eq. (4): $\boldsymbol{\beta}_{\text{direct}} = \begin{pmatrix} 1 & -1 & 0 & 0 \\ -1 & 1 & 0 & 0 \\ 0 & 0 & 0 & 0 \\ 0 & 0 & 0 & 0 \end{pmatrix}$,

  whose rows correspond to, for example: turning left, turning right, turning around, wandering; and whose columns correspond to, for example: a reward-predominant decision-dimension 1, a cost-predominant decision-dimension 2, a hunger-predominant decision-dimension 3, and a location-predominant decision-dimension 4. The coefficients model a T-maze where a choice is made to turn right or left based on relative values of cost and reward.

- in eq. (4): $\alpha_{j,P} = -3$ for all $j$ and $P$
- in eq. (6): $\sigma = 1$
- in eq. (7), (8): $h = 2$

**Effect of reward/costs on LHb/RMTg/daSNC activity.**

In **Figures S1E,F,** we modeled the effect of incrementing reward or cost on the activities of LHb, RMTg, and daSNC.

The inputs enter the model circuit in two ways: 1) reward and cost are mapped to decision-dimensions; and 2) cost level leads to changes in LHb and RMTg activities, similar to what has been demonstrated in experimental work[135,136,215–217]. The modeled LHb and RMTg responses to cost are proportional to cost level with an arbitrary coefficient (set to 1 for LHb and 0.9 for RMTg for the purposes of plotting).

The modeled results show that the mean activity of a daSNC subpopulation encoding reward-predominant data responds positively to increases in reward and negatively to decreases in reward, similar to experimental evidence[218]. LHb and RMTg respond negative linearly to reward level and positive linearly to cost level, similar to experimental evidence[135,144]. Sudden changes in reward or cost level, therefore, lead to shifts in activities that track changes to expectations of future reward or cost value, including reward or cost currently received, i.e. reward or cost prediction error.

For code, see https://github.com/dirkbeck/DM_space_model/blob/main/model_overview/GPi_LHb_RMTg_DA_model.m.

**Effect of LHb/RMTg/daSNC activity on decision-space.**

In **Figures S1G-I**, we modeled the effect of incrementing GPi, LHb, RMTg, or daSNC activity on the type of decision-space formed during a decision.

In the plotted analysis, we altered $z_{\mathrm{GPi}}$ in eq. (5), $z_{\mathrm{LHb}}$ in eq. (5), $z_{\mathrm{RMTg}}$ in eq. (5), and $z_{\mathrm{daSNC},i,P}$ (uniform change for all $i$, a single pathway is considered) in eq. (5) such that they took 10 values incremented from 0 to 1. Parameters not altered took default values (see **Common parameters**).

We also examined the role of each component in decision-space formation through the perspective of a series of steps, each carried out by a different circuit element. For this analysis, we substituted eq. (5) into eq. (2) and altered each parameter in turn. The plots illustrate the value of $\mathrm{daSNC}_{i,P}$ if the other parameters were set to 1 ($z_{\mathrm{GPi}}$) or 0 ($z_{\mathrm{RMTg}}$, $z_{\mathrm{daSNC},i,P}$). $b_{\mathrm{LHb}}$ is set to 0.5 (control), -5 (lesioned LHb), or 5 (stimulated LHb).

See **Table S4** for alignment to the experimental literature.

For code, see https://github.com/dirkbeck/DM_space_model/blob/main/model_overview/GPi_LHb_RMTg_DA_model.m.

**Effect of sSPN activity on decision-space.**

In the analysis plotted in **Figure 2A,** we incremented $b_{\mathrm{sSPN}}$ in eq. (1) and, for each increment, recorded $\mathrm{daSNC}_i$ in eq. (2). Then, using the approach in eq. (6), we converted the probability that one decision-dimension is used in the formation of decision-space to the probability that a decision-spaces of a certain dimensionality is formed.

For code, see https://github.com/dirkbeck/DM_space_model/blob/main/model_tests/friedman2015optogeneticmanipulation.m.

**Effect of decision-space on choice.**

In the analysis plotted in **Figure 2B**, we changed which decision-space was formed by mSPNs and measured choice.

The excitation group was modeled using a non-dimensional decision-space (dopamine→mSPN weights of 0 reward-predominant decision-dimension, 0 cost-predominant dimension). The control group was modeled using a 1D direct pathway decision-space (dopamine→mSPN weights of 0.5 reward-predominant dimension, 0 cost-predominant dimension). The inhibition group was modeled using a 2D direct pathway decision-space (dopamine→mSPN weights of 1 reward-predominant dimension, 1 cost-predominant dimension). The modeled T-maze task was a choice between reward=2, cost=1 (high reward, high cost) and reward=1, cost = 0.5 (low reward, low cost). 20 simulations were run per modeled subject for 100 subjects. Other parameters for forming decision-space and calculating action value are set to their defaults (see **Common parameters**). For simplicity, the indirect pathway is not modeled in this analysis.

For code, see
https://github.com/dirkbeck/DM_space_model/blob/main/model_tests/friedman2015optogenetic
manipulation.m.

In the analysis in **Figures 3B,C**, we modeled changes to decision-space and choice after stress.

Here, modeled control rodents made decisions using a 2D direct pathway decision-space formed from reward-predominant and cost-predominant decision-dimensions. This correspond mathematically to a truncation of $\boldsymbol{\beta}_{\text{direct}}$ (see eq. (4) and **Common parameters**) to two columns. The first subset of modeled stress-group rodents made decisions without forming direct pathway decision-space. This corresponds to an elimination of $\boldsymbol{\beta}_{\text{direct}}$ such that action value is defined purely based on priors ($\alpha_{j,\text{direct}}$ in eq. (4)). The second subset made decisions without forming direct pathway decision-space until they reached a critical threshold, beyond which they formed a 1D direct pathway decision-space with a reward-predominant dimension. Action values are derived for the three groups across multiple reward and cost combinations (**Figure 3C**) via eqs. (3) and (4). Then choices are modeled using eqs. (6), (7), and (9) across 2000 simulations per group for each reward concentration (each incremented from 0 to 1 arbitrary units, 7 increments). Cost concentration is set to 0.5 arbitrary units (set at this level to resemble the steepness of increase in the experimental psychometric function). Default parameters are used for action value formation and the Merton process model. For simplicity, the indirect pathway is not modeled in this analysis. **Figure 3B** plots the averages of the simulations.

For code, see
https://github.com/dirkbeck/DM_space_model/blob/main/disorder_hypotheses/Friedman2017_lowD_space.m.

In the analysis plotted in **Figures 3D,E**, we modeled the effect on choice of shifts in decision-space after a small cost is added to a reward (experimental data is plotted in **Figure S3J**).

Rodents in the only-reward task were modeled as forming a lower-dimensional direct pathway decision-space (decision-dimension 1 weight = 0.5, decision-dimension 2 weight = 0.2) while animals in the reward-and-cost task formed a higher-dimensional direct pathway decision-space (decision-dimension 1 weight = 1, decision-dimension 2 weight = 0.5).

To do this, we truncated $\boldsymbol{\beta}_{\text{direct}}$ (see eq. (4) and **Common parameters**) to two columns or derived action value purely based on priors ($\alpha_{j,\text{direct}}$ in eq. (4)). A cortical input of reward = 0.7, cost = 0.3 is shown in the plots. For simplicity, the indirect pathway is not modeled in this analysis.

For code, see
https://github.com/dirkbeck/DM_space_model/blob/main/disorder_hypotheses/altered_choice_after_space_transition.m.

In the analysis plotted in **Figure 3H**, we modeled changes to choice after aging in young and old groups.

Here, we truncated $\boldsymbol{\beta}_{\text{direct}}$ (see eq. (4) and **Common parameters**) to two decision-dimensions, the first corresponding to a reward-predominant decision-dimension and the second corresponding to a cost-predominant decision-dimension. In the current analysis, the first row of $\boldsymbol{\beta}_{\text{direct}}$ corresponded to licking while the second row corresponds to performing a different action, e.g. movement. The licking action was assigned a larger prior, $\alpha_{1,\text{direct}}=0$, $\alpha_{2,\text{direct}}=-3$, due to the strong association developed in the rodents between the experimental apparatus and licking. For the modeled "learned, young" group, no decision-space is formed during the reward-cue task and a decision-space using only a cost-predominant decision-dimension is formed during the cost-cue task (i.e. $\mathbf{s}_{\text{mSPN, reward, direct}}=\begin{bmatrix}0\\0\end{bmatrix}$, $\mathbf{s}_{\text{mSPN, cost, direct}}=\begin{bmatrix}0\\1\end{bmatrix}$ in eq. (3)). For the modeled "learned, old" group, no decision-space is formed during the reward-cue task and a decision-space involving a cost-predominant decision-dimension is partially formed during the cost task ( $\mathbf{s}_{\text{mSPN, reward, direct}}=\begin{bmatrix}0\\0\end{bmatrix}$, $\mathbf{s}_{\text{mSPN, cost, direct}}=\begin{bmatrix}0\\0.5\end{bmatrix}$). For the "not learned" group, a decision-space involving a cost-predominant decision-dimension is partially formed during both tasks ( $\mathbf{s}_{\text{mSPN, reward, direct}}=\begin{bmatrix}0\\0.5\end{bmatrix}$, $\mathbf{s}_{\text{mSPN, cost, direct}}=\begin{bmatrix}0\\0.5\end{bmatrix}$). For simplicity, the indirect pathway is not modeled in this analysis.

For code, see https://github.com/dirkbeck/DM_space_model/blob/main/disorder_hypotheses/Friedman2020_lowD_space.m.

**Effect of FSI activity on decision-space.**

In the analysis plotted in **Figure 3F**, we incremented FSI activity $a_{\text{FSI}}$ in eq. (23) and determined the response of $\text{daSNC}_i$ in eq. (2) (a single pathway is considered). The activity parameters related to other circuit elements were held constant (see **Common parameters**).

For code, see https://github.com/dirkbeck/DM_space_model/blob/main/disorder_hypotheses/space_dimensionality_vs_FSI.m.

**Effect of dopamine on action/inaction values.**

In the analyses plotted in **Figures S5C-F**, we measured the effect of high versus low dopamine on action and inaction values across a range of cortical inputs to the system.

We modeled a cost-benefit conflict task with increasing reward (scale of 0 to 1 arbitrary units, 100 increments) and constant cost (set to 0.25 arbitrary units). Experimental work has shown that dopamine increases direct pathway activity while decreasing indirect pathway activity and vice versa[26]. Therefore, we set coefficients relating to overall activity of the pathways oppositely: in the low dopamine case, the direct pathway coefficient was 0.1 arbitrary unit and the indirect pathway coefficient 5 arbitrary units; and in the high dopamine case, the indirect pathway coefficient was 5 arbitrary units and the direct pathway coefficient 0.1 arbitrary unit. These coefficients were multiplied by $\boldsymbol{\beta}_{\mathrm{direct}}$ or $\boldsymbol{\beta}_{\mathrm{indirect}}$ in eq. (4), increasing or decreasing the overall sensitivity of action value on data along cortical principal components. In the model, changes to dopamine also involved a change in decision-space: due to their opposite effects on mSPN activity, dopamine biases the direct pathway towards forming higher-dimensional decision-spaces and the indirect pathway towards forming lower-dimensional decision-spaces. For the purpose of this analysis, eq. (3) is reframed to incorporate the effects of dopamine in scaling action value score ($A$, set to either 5 or 0.1 arbitrary units in our analysis) and changing decision-space ($B$, set to 1 arbitrary unit when dopamine is high and 0 when dopamine is low). Here, individual elements are referenced through subscripts based on their $j$ th row and column corresponding to the reward or cost dimension.

(24) $$v_{j,\mathrm{direct}} = \frac{1}{1 + \exp\left( -A \cdot \left( \beta_{j,\mathrm{reward,direct}} + B \cdot \beta_{j,\mathrm{cost,direct}} \right) - \alpha_{j,\mathrm{direct}} \right)}$$

(25) $$v_{j,\mathrm{indirect}} = \frac{1}{1 + \exp\left( -A \cdot \left( (I - B) \cdot \beta_{j,\mathrm{reward,indirect}} + \beta_{j,\mathrm{cost,indirect}} \right) - \alpha_{j,\mathrm{indirect}} \right)}$$

where:

- $A$ is the multiplicative effect of dopamine released to mSPNs (dimensionless)
- $B$ is the effect of dopamine on decision-space (dimensionless)
- $\beta_j$ is the connection weight from mSPN to an action value neuron. Each $\beta_{j,\mathrm{reward}}$ or $\beta_{j,\mathrm{cost}}$ corresponds to an element of the connection weight matrix $\boldsymbol{\beta}_P$.
- $\mathrm{prior}_j$ is an additive shift corresponding to the neuron encoding action $j$ (activity arb. u.)

The plot in **Figure S5C** compares the high dopamine and low dopamine cases. The plot in **Figure S5D** shows a similar analysis but for changes in parameters: cost is fixed at 0.5 arbitrary units, and $A$ is set to either 2 arbitrary units (corresponding to the pathway not disconnected) or 0 (corresponding to the pathway disconnected). The plots in **Figures S5E,F** show progress to action in the case where reward = 1 arbitrary unit and cost = 0.25 arbitrary unit for low versus high dopamine. Deliberation time distributions are formed by aggregating the deliberation times across the 100 simulations. For parameters used for subjective valuation and deliberation time simulation, see **Common parameters**.

For code, see https://github.com/dirkbeck/DM_space_model/blob/main/dynamic_model_and_neural_net/direct_vs_indirect_pathway_SV.m.

**Effect of decision-dimensions on choice.**

In the analysis plotted in **Figure S5I**, we altered the connections between mSPN and action/inaction encoding neurons on choice.

A modeled approach/avoid experiment is conducted by offering an option with reward = 1 arbitrary unit, cost = 1 arbitrary unit, and varying (10 values incremented from [0, 2] arbitrary units) physically proximity to another reward. An additional column is added to $\boldsymbol{\beta}_{\text{direct}}$ and $\boldsymbol{\beta}_{\text{indirect}}$ in eq. (4) to reflect the fact that additional proximity to the other reward increases approach rate: $\boldsymbol{\beta}_{\text{direct}}=\boldsymbol{\beta}_{\text{indirect}}=\begin{bmatrix} 1 & -1 & 1 \\ -1 & 1 & -1 \end{bmatrix}$, where the upper row corresponds to approaching, the bottom row corresponds to not approaching, and the columns correspond to, from left to right, a reward-predominant decision-dimension, a cost-predominant decision-dimension, and a location-predominant decision-dimension. Reward is assigned a greater relative importance than cost or location (score of 3 arbitrary units versus 1 versus 1) in sSPNs, while cost is assigned a greater relative importance than reward or location (score of 3 arbitrary units versus 1 versus 1). daSNC activity is incremented by changing $z_{\text{daSNC},i,P}$ in eq. (2) for all $i$ and both pathways. Action and inaction values are calculated, and then choice is formed by averaging the results of 1000 Merton process simulations.

For code, see https://github.com/dirkbeck/DM_space_model/blob/main/dynamic_model_and_neural_net/direct_vs_indirect_pathway_proximity_theory.m.

**Effect of cortical SNR on choice.**

In the analysis plotted in **Figures 5A-E**, cortical signal to noise ratio (SNR) is altered and the effect on choice is simulated.

Merton process simulations (see **Defining Choice**) are run across ten increments of reward and cost from -1 to 1 arbitrary units for a modeled cost-benefit conflict task. Parameters related to action value are set to their defaults and the T-maze task is used (see **Common parameters**). Here, “turn right” corresponds to receiving the reward and cost combination, while other actions correspond to receiving no reward and no cost. For simplicity, only the direct pathway is used to influence choice. 100 simulations are run for each of 100 reward and cost combinations, and for each combination, choice is averaged.

The above process is replicated with changes to two sets of parameters. First, the effect of changes to decision-space were considered. A different $\mathbf{S}$ in eq. (3) was used depending on specified decision-space: $\mathbf{S}=\begin{bmatrix} 1 & 0 & 0 \\ 0 & 0 & \vdots \\ 0 & \cdots & 0 \end{bmatrix}$ for 1D decision-spaces, and $\mathbf{S}=\begin{bmatrix} \mathrm{I}_2 & & 0 \\ & & \vdots \\ 0 & \cdots & 0 \end{bmatrix}$ for 2D decision-spaces. Second, changes to cortical noise were considered by adding i.i.d. Gaussian noise to $\mathbf{x}_{\text{direct}}$ (with mean 0 and standard deviation $\sigma$) at every time step, then recalculating action values in eq. (4) based on the mSPN activities calculated at that time step. Simulations were run for $\sigma = 1, 2, \ldots, 10$. Default parameters were used for calculating action value (see **Common parameters**).

Examples of single simulations at each level of reward and cost are shown for the $\sigma = 1$ (high cortical SNR) and $\sigma = 5$ (low cortical SNR) cases in **Figures 5A-D**. In **Figure 5E,** expected value is averaged across the 100 simulations for each noise level. Expected value here is defined as reward minus 0.75*cost (to add preference for reward compared to cost, coefficient is arbitrary) achieved across reward and cost levels. In the plot, SNR is set to the inverse of $\sigma$.

For code, see https://github.com/dirkbeck/DM_space_model/blob/main/dynamic_model_and_neural_net/cortical_snr.m.

**Effects of sSPN, LHb, and daSNC activity on decision-space.**

In **Figures 6A,B**, 100 values of $b_{\text{sSPN}}$ (eq. (1)), $z_{\text{LHb}}$ (eq. (5)), and $z_{\text{daSNC}, i, P}$ (for all $i$ and a single pathway, eq. (2)) are selected uniformly at random from a range of 0 to 10 arbitrary units. Each of the 100 points are used to derive $\text{daSNC}_i$ in eq. (2) and converted to decision-spaces via eq. (18).

For code, see https://github.com/dirkbeck/DM_space_model/blob/main/day_to_day_space_sampling/sampling_space_based_on_activity.m.

**Effect of decision-space on choice profiles.**

In **Figures 6C,F, S6D,F**, we form decision-spaces using various decision-dimensions across incremented cortical reward and cost inputs, then classified the action values formed across those reward/cost inputs using a scoring system.

The scoring system, visualized in **Figure S6C**, is as follows:

Scores for “explore,” “riskiness,” “high action,” “exploit,” “safety,” “low action” are calculated by incrementing reward and cost on [-1 1] (arbitrary units) scales (9 increments are used for each of reward and cost in **Figures 6C, S6D,** 6 increments in **Figures 6F, S6F**). The notation used here treats $v_{r,c,j}$ as the action value of the $j$th action at a certain reward and cost increment and $v_{r,c}$ as the set of those action values. In the plotted analysis, $k=4$ actions are assigned action values.

- Explore. The tendency to pursue multiple actions simultaneously. Scored as the area of the region of reward and cost combinations with a Gini coefficient less than 0.25.

(26) $$\text{explore}=\sum_{\text{r}=-1}^{1}\sum_{\text{c}=-1}^{1}\left[\text{gini}\left(v_{r,c}\right)<0.25\right]$$

(27) $$\text{gini}\left(v_{r,c}\right)=\frac{\sum_{i=1}^{k}\sum_{j=1}^{k}\left|v_{r,c,i}-v_{r,c,j}\right|}{2k\sum_{j=1}^{k}v_{r,c,j}}$$

- Exploit. The tendency to pursue only one action. Scored at the area of the region of reward and cost combinations with a Gini coefficient greater than 0.5.

(28) $$\text{exploit}=\sum_{\text{r}=-1}^{1}\sum_{\text{c}=-1}^{1}\left[\text{gini}\left(v_{r,c}\right)>0.5\right]$$

- Riskiness. The combined value of actions when reward and cost are high. Scored by examining the combinations where both reward and cost are greater than 0.

(29) $$\text{riskiness}=\sum_{\text{r}=0}^{1}\sum_{\text{c}=0}^{1}\sum_{j=1}^{k}v_{r,c,j}$$

- Safety. The combined value of actions when reward and cost are low. Scored by examining the combinations where both reward and cost are less than 0.

(30) $$\text{safety}=\sum_{\text{r}=-1}^{0}\sum_{\text{c}=-1}^{0}\sum_{j=1}^{k}v_{r,c,j}$$

- High action. How often actions will have high action values. Scored as the area of the region of reward and cost combinations that have combined action value greater than 0.5.

(31) $$\text{high action}=\sum_{\text{r}=-1}^{1}\sum_{\text{c}=-1}^{1}\left[\sum_{j=1}^{k}v_{r,c,j}>0.5\right]$$

- Low action. How often actions will have low action values. Scored as the area of the region of reward and cost combinations that have combined action value less than 0.2.

(32) $$\text{high action}=\sum_{\text{r}=-1}^{1}\sum_{\text{c}=-1}^{1}\left[\sum_{j=1}^{k}v_{r,c,j}<0.2\right]$$

In the analyses plotted in **Figure 6C** and **Figure S6D** and the examples in **Figure 6D**, action value scores, as measured by the scoring definitions above*,* are compared when different decision-spaces are constructed but cortical input and system parameters are unchanged. The underlying action values across reward and cost levels resembles those from other analyses (see **Common parameters**) except for an addition of normal random noise (mean = 0, standard deviation = 1) to every element of $\boldsymbol{\beta}_{\text{direct}}$ (see eq. (4)).

The analysis in **Figure S6D** is similar, except for here, a weighted average is taken of action value scores, as measured by the scoring definitions above, between scenarios where different decision-spaces are constructed. A different $\mathbf{S}$ in eq. (3) is used depending on the specified dimensionality of direct pathway decision-space: $\mathbf{S}=\begin{bmatrix} 1 & 0 & 0 \\ 0 & 0 & \vdots \\ 0 & \cdots & 0 \end{bmatrix}$ for 1D, $\mathbf{S}=\begin{bmatrix} \mathbf{I}_2 & & 0 \\ & & \vdots \\ 0 & \cdots & 0 \end{bmatrix}$ for 2D, $\mathbf{S}=\begin{bmatrix} \mathbf{I}_3 & & 0 \\ & & \vdots \\ 0 & \cdots & 0 \end{bmatrix}$ for 3D, and $\mathbf{S}=\mathbf{I}_4$ for 4D. A weighted average of the five decision-spaces is calculated for three levels of sSPN activity (-1, 0, and 1).

For code, see https://github.com/dirkbeck/DM_space_model/blob/main/day_to_day_space_sampling/subjective_value_scores_by_space.m.

In the analyses plotted in **Figures 6F, S6F**, for each of 1000 simulations, uncorrelated Gaussian white noise (mean = 0, standard deviation = 1) is added to every element of $\boldsymbol{\beta}_{\text{direct}}$ (see eq. (4)) and 6 by 6 grids of action values across reward and cost combinations are scored by the “explore,” “riskiness,” “high action,” “exploit,” “safety,” and “low action” metrics. Scores for each metric are compared across simulations and between decision-space groups. Observations that score in the top 10% by a metric are considered outliers. Outlier proportion is plotted in **Figure 6F**. The means across the simulations of each score are plotted in **Figure S6F**.

For code, see https://github.com/dirkbeck/DM_space_model/blob/main/day_to_day_space_sampling/subjective_value_score_extremes.m.

**Effect of decision-space on sSPN-mSPN correlation**

In the analysis plotted in **Figure 2E,** sSPN-mSPN correlation is compared across decision-spaces with different dimensionality.

It is assumed in the plotted examples that 1D decision-spaces are only formed from the first decision-dimension, 2D decision-spaces are only formed from the first and the second, and 3D decision-spaces are only formed from the first, second, and third. The analysis assumes a comparison of SPNs of the same pathway (that is, either dsSPN-dmSPN or isSPN-imSPN). For this analysis, eigenvalues of cortical activity are set to 2, 1, 0.5, 0.2, and 0.1, respectively. Weighted averages of example signals (left panel) and correlation for different decision-spaces (right panel) are formed using the identity that eigenvalues of principal components are equivalent to their variances.

For code, see https://github.com/dirkbeck/DM_space_model/blob/main/model_tests/ctx_sSPN_mSPN_coordinated_activity.m.

***Instance 2: sparse connectivity and feedforward.***

In this section, we describe the instance of the model where cortex input to the system does not change over time and the activities of other circuit elements do not decay over time.

This instance of the model leads to a convenient formation of the model as a circuit with no time component. In this section, we frame this instance mathematically and then describe our related analysis examining the process of formation of the decision-space over time.

To focus on the portions of the circuit we analyze using this instance, we define here the subset of the circuit involving cortex, FSI, sSPN, daSNC, and mSPN.

For code, see https://github.com/dirkbeck/DM_space_model/blob/main/dynamic_model_and_neural_net/neural_network_model.m.

**Cortical input.**

A set of 4 cortical neurons, notated as $C$, is sampled at random from a population of 50 cortical neurons. Each neuron in $C$ projects to one FSI and each of $q$ SPNs, which each correspond to a decision-dimension. In our analysis, $q$ is set to 4. This process is repeated 10,000 times per each pathway, forming 10,000 groups of 4 cortical neurons, 1 FSI, 4 dsSPNs (or isSPNs), and 4 dmSPNs (or imSPNs) for each pathway.

**Defining FSI activity.**

FSI activity is defined as a weighted sum of the activities of connected cortical neurons:

(33) $$\mathrm{FSI}_C = \sum_{q \in C} w_{\mathrm{cortex} \rightarrow \mathrm{FSI}} \, \mathrm{cortex}_q + b_{\mathrm{FSI}}$$

where:

- $C$ is a randomly sampled subset of cortical neurons.
- $\mathrm{FSI}_C$ is the activity of the FSI which receives projection from the cortical neurons in $C$ (activity arb. u.)
- $w_{\mathrm{cortex} \to \mathrm{FSI}}$ is the connection weight between cortical neurons and FSIs (dimensionless)
- $\mathrm{cortex}_q$ is the activity of cortical neuron $q$ (activity arb. u.)
- $b_{\mathrm{FSI}}$ affects the relative activity of all sSPN neurons (activity arb. u.)

In the current instance of the model, there are 10,000 FSIs that project to each of dSPNs and iSPNs. Each cortical neuron in $C$ projects to $\mathrm{FSI}_C$.

**Defining sSPN activity.**

sSPN activity is defined as a weighted sum of the activities of connected cortical neurons, divided by a weighted sum of the activities of connected FSIs, plus an additive shift $b_{\mathrm{sSPN}}$ applied to all sSPNs:

(10) $$\mathrm{sSPN}_{s,C} = \frac{1}{|C|} \sum_{q \in C} \frac{w_{q \to s}\, \mathrm{cortex}_q}{\mathrm{FSI}_C} + b_{\mathrm{sSPN}}$$ (copied from **Figure S4** for convenience)

where:

- $C$ is a randomly sampled subset of cortical neurons.
- $\mathrm{sSPN}_{s,C}$ is the activity of an sSPN $s$ that receives projection from cortical neurons in $C$. (activity arb. u.)
- $w_{q \to s}$ is the connection weight between cortical neuron $q$ and sSPN $s$. The weight is equivalent to one of the first four principal components of the cortical activity of the four connected cortical neurons. sSPNs are separated into equal populations that correspond to the first, second, third, or fourth principal component. (dimensionless)
- $\mathrm{FSI}_C$ is the activity of the FSI which receives projection from the cortical neurons in $C$ (activity arb. u.)
- $\mathrm{cortex}_q$ is the activity of cortical neuron $q$ (activity arb. u.)
- $b_{\mathrm{sSPN}}$ represents the relative activity of all sSPN neurons (activity arb. u.)

In the current instance of the model, there are 40,000 neurons for each of dsSPNs and isSPNs. Activities are defined based on a feedforward network, so the simplification is made that sSPN activities are not affected by daSNC activities. All cortical neurons in $C$ project to $\mathrm{sSPN}_{s,C}$ for all $s$, and similarly, $\mathrm{FSI}_C$ projects to $\mathrm{sSPN}_{s,C}$ for all $s$.

**Defining daSNC activity.**

The activity of the daSNC element corresponding to decision-dimension $i$ and pathway $P$ is defined as the weight from sSPNs corresponding to decision-dimension $i$ and pathway $P$:

(11) $$\mathrm{daSNC}_{i,P} = \frac{1}{1+\exp\left(\frac{1}{n_{\mathrm{sSPN}}}\sum_{s\in i,P} w_{s\rightarrow \mathrm{daSNC},i,P}\cdot \mathrm{sSPN}_s + \mathrm{RMTg} - z_{\mathrm{daSNC},i,P}\right)}$$

(copied from **Figure S4** for convenience)

where:

- $\mathrm{daSNC}_{i,p}$ is the activity the daSNC neuron corresponding to decision-dimension $i$ and pathway $P$. (activity arb. u.)
- $n_{\mathrm{sSPN}}$ is the count of sSPNs in each of the direct/indirect pathways
- $w_{s\rightarrow \mathrm{daSNC},i,P}$ is the connection weight from sSPN $s$ to the daSNC neuron corresponding to decision-dimension $i$ and pathway $P$. The weight is fixed in this instance of the model. (dimensionless)
- $\mathrm{sSPN}_s$ is the activity of SPN $s$ (activity arb. u.)
- $\mathrm{RMTg}$ is RMTg activity. (activity arb. u.)
- $z_{\mathrm{daSNC},i,P}$ is the bias in the activity of a daSNC neuron corresponding to decision-dimension $i$ and pathway $P$. (activity arb. u.)

In the current instance of the model, there are $q$ daSNC neurons that receive projection from the 10,000 dsSPNs corresponding to each decision-dimension, and likewise $q$ daSNC neurons that receive projection from the 10,000 isSPNs corresponding to each decision-dimension. RMTg also projects to all daSNC neurons.

In our analysis using this instance of the model, we arbitrarily set $w_{\mathrm{sSPN}\rightarrow \mathrm{daSNC},i}=1$ arbitrary unit for both pathways and $z_{\mathrm{daSNC},i}=-5$ arbitrary units for all $i$ for the direct pathway, and $z_{\mathrm{daSNC},i}=5$ arbitrary units for all $i$ for the indirect pathway. For simplicity, $\mathrm{RMTg}$ is set to 0.

**Defining mSPN activity and decision-space.**

mSPN activity is defined as a weighted sum of the activities of connected cortical neurons, divided by a weighted sum of the activities of connected FSIs. Here, unlike in the definition of sSPN activity in eq. (10), a $d_{i,P}$ term is multiplied to incorporate the weighting of mSPNs by dopamine:

(12) $$\mathrm{mSPN}_{m,C} = \frac{d_{i,P}}{|C|} \sum_{q \in C} \frac{w_{q \to m}\, \mathrm{cortex}_q}{\mathrm{FSI}_C}$$ (copied from **Figure S4** for convenience)

where:

- $C$ is a randomly sampled subset of cortical neurons.
- $\mathrm{mSPN}_{m,C}$ is the activity of mSPN $m$ that receives projection from cortical neurons in $C$. (activity arb. u.)
- $d_{i,P}$, which takes the value 0 or 1, is dopamine signaling to mSPNs corresponding to decision-dimension $i$ and pathway $P$. $d_{i,P}$ is the realization of probabilistic weighting of decision-dimensions based on daSNC activity (see ***Conceptual Model***). (dimensionless)
- $w_{q \to m}$ is the connection weight between cortical neuron $q$ and mSPN $m$. The weight is equivalent to one of the first four principal components of the cortical activity of the four connected cortical neurons. As described in ***Conceptual Model,*** sSPNs are separated into equal populations that correspond to the first, second, third, or fourth principal component. (dimensionless)
- $\mathrm{FSI}_C$ is the activity of the FSI which receives projection from the cortical neurons in $C$ (activity arb. u.)
- $\mathrm{cortex}_q$ is the activity of cortical neuron $q$ (activity arb. u.)

In the current instance of the model, there are 40,000 neurons for each of dmSPNs and imSPNs All cortical neurons in $C$ project to $\mathrm{mSPN}_{m,C}$ for all $m$, and similarly, $\mathrm{FSI}_C$ projects to $\mathrm{mSPN}_{m,C}$ for all $m$.

***Modeling SPN encoding of data, using Instance 2.***

To explore the ability of SPNs to successfully encode data along decision-dimensions, even when cortex and SPNs are sparsely connected, we constructed networks with different degrees of dimensionality reduction (**Figure S4G**). A single pathway is considered. One type of network had 2 times dimensionality reduction (20 cortical neurons, 10 SPN), another had 10 times dimensionality reduction (100 cortical neurons, 10 SPN), and another had 100 times dimensionality reduction (1000 cortical neurons, 10 SPN), similar to what is found in the human brain[99,114].

For each type, we constructed modeled networks with cortex→SPN connections equal to principal components of cortical activity by simulating, for each analyzed pathway, a random symmetric positive definite matrix that is used as the cortical covariance matrix $\mathbf{\Sigma}_P$ (see **Defining sSPN activity**) via MATLAB's sprandsym() with density=1. Eigenvalues are arbitrarily specified as $\lambda_1 = 2$, $\lambda_2 = 2$, $\lambda_3 = 0.5$, $\lambda_4 = 0.2$, and $\lambda_5 = \lambda_6 = \ldots = \lambda_p = 0$, i.e. the first and second principal components are very important, the third somewhat important, the fourth slightly important, and the others unimportant. The weights $\mathbf{W}_P$ from cortical neurons to the $q$ SPN circuit elements per pathway are then derived as the first $q$ eigenvectors of $\mathbf{\Sigma}_P$.

We incremented the number of cortical neurons that connected to each SPN from 2 to 10. The network was connected sparsely based on the specified number of connections from randomly selected cortical neurons to each SPN. For simplicity in this analysis, we created a cortical signal that resembled the first principal component of cortical activity as a whole (regardless of connectivity to SPNs) and let the first cortical principal component to have large eigenvalue compared to the others (i.e. $\lambda_1 = 1$, $\lambda_2 = 0.1$, $\lambda_3 = \lambda_4 = \ldots = \lambda_p = 0$).

For each of the modeled networks (3 network types by 9 increments from 2 to 10), we simulated the process 1000 times. During each simulation, we calculated the ability of the network to discriminate between the large signal along the first cortical principal component and the absence of signal along the second cortical principal component, given its access to only a subset (2 to 10 cortical neurons) of the complete signal.

Then the SPNs encoding data along the first versus second decision-dimension were assessed in their ability to distinguish between signals along the first cortical principal component versus the second. This was quantified using the Bhattacharyya distance of the activity among SPNs encoding data along the first decision-dimension versus the second, assuming the subpopulations have mean activities $\mu_1$ and $\mu_2$ and standard deviations $\sigma_1$ and $\sigma_2$, respectively.

(34) $$D_B = \frac{1}{4}\frac{\left(\mu_1 - \mu_2\right)^2}{\sigma_1^2 + \sigma_2^2} + \frac{1}{2}\ln\left(\frac{\sigma_1^2 + \sigma_2^2}{2\sigma_1\sigma_2}\right)$$

where:

- $\mu_1$ is the mean activity of activities in the first subpopulation (activity arb. u.)
- $\mu_2$ is the mean activity of activities in the second subpopulation (activity arb. u.)
- $\sigma_1$ is the standard deviation of activities in the first subpopulation (activity arb. u.)
- $\sigma_2$ is the standard deviation of activities in the second subpopulation (activity arb. u.)

For the current analysis, $a_{\mathrm{FSI}}$ is arbitrarily set to 1 arbitrary unit and $b_{\mathrm{FSI}}$ is arbitrarily set to 0.

For code, see https://github.com/dirkbeck/DM_space_model/blob/main/dynamic_model_and_neural_net/dimension_discrimination_vs_sparsity.m.

***Instance 3: full connectivity and dynamics.***

In this section, we describe the instance of the model where each cortical neuron projects to each FSI, each FSI projects to each SPN (for dsSPN, isSPN, dmSPN, imSPN), and each cortical neuron projects to each SPN.

This instance of the model leads to a convenient formation of the model as a circuit of fewer elements (one FSI, one dsSPN, isSPN, dmSPN, and imSPN per the four decision-dimensions), with a time component. In this section, we frame this instance mathematically and then describe our related analysis.

To focus on the portions of the circuit we analyze using this instance, we define here the subset of the circuit involving cortex, sSPN, daSNC, and mSPN.

**Defining SPN and daSNC activity.**

Here, the activities of SPN and daSNC elements and the weights from sSPN to daSNC are represented as a system of differential equations. Because FSI activity is not measured in the related analyses, cortical activity to pathway $P$ after FSI normalization $x_{i,P}(t)$ is used as input to the system in the equations below.

For a diagram of the model with dynamics, see **Figure 4B**. Note that daSNC activity of 0 (i.e. average activity) leads to 0 change in SPN activity (due to the $1/2$ terms in eqs. (13) and (14).

(13) $$\tau \cdot \frac{\mathrm{d}s_{\mathrm{sSPN},i,P}(t)}{\mathrm{d}t} = -s_{\mathrm{sSPN},i,P}(t) - x_{i,P}(t) - w_{\mathrm{daSNC}\to\mathrm{sSPN},i,P} \cdot \left(y_{\mathrm{sSPN},i,P}(t) - \frac{1}{2}\right)$$

(copied from **Figure S5** for convenience)

(14) $$\tau \cdot \frac{\mathrm{d}s_{\mathrm{mSPN},i,P}(t)}{\mathrm{d}t} = -s_{\mathrm{mSPN},i,P}(t) + x_{i,P}(t) + w_{\mathrm{daSNC}\to\mathrm{mSPN},i,P} \cdot \left(y_{\mathrm{sSPN},i,P}(t) - \frac{1}{2}\right)$$

(copied from **Figure S5** for convenience)

(15) $$\frac{\mathrm{d}}{\mathrm{d}t} w_{\mathrm{sSPN} \rightarrow \mathrm{daSNC}, i, P}(t) = \kappa \cdot s_{\mathrm{sSPN}, i, P}(t)$$

(copied from **Figure S5** for convenience)

where:

(16) $$y_{\mathrm{sSPN}, i, P}(t) = \frac{1}{1 + \exp\left( w_{\mathrm{sSPN} \rightarrow \mathrm{daSNC}, i, P}(t) \cdot s_{\mathrm{sSPN}, i, P}(t) + \mathrm{RMTg} - z_{\mathrm{daSNC}, i, P} \right)}$$

(copied from **Figure S5** for convenience)

- $\tau$ is the time constant related to the decay rate of activity (dimensionless)
- $s_{i,P}(t)$ is the SPN activity (either dsSPN, isSPN, dmSPN, or imSPN) corresponding to decision-dimension $i$ and pathway $P$, as a function of time. (activity arb. u.)
- $t$ is time (seconds)
- $x_{i,P}(t)$ is the cortical activity input, after FSI normalization, to an SPN corresponding to decision-dimension $i$ and pathway $P$ (activity arb. u.)
- $w_{\mathrm{daSNC} \rightarrow \mathrm{sSPN}, i, P}$ is the connection weight from a daSNC neuron corresponding to decision-dimension $i$ and pathway $P$ to an sSPN (either dsSPN or isSPN) corresponding to decision-dimension $i$ and pathway $P$. (dimensionless)
- $y_{i,P}(t)$ is the activity of the daSNC neuron corresponding to decision-dimension $i$ and pathway $P$, as a function of time. (activity arb. u.)
- $w_{\mathrm{sSPN} \rightarrow \mathrm{daSNC}, i, P}(t)$ is the connection weight from an sSPN (either dsSPN or isSPN) to a daSNC neuron corresponding to decision-dimension $i$ and pathway $P$, as a function of time. (dimensionless)
- $\kappa$ is a coefficient that determines the rate at which the connection from sSPNs to daSNC neurons change depending on sSPN (dsSPN or isSPN) activity. (dimensionless)
- $z_{\mathrm{daSNC}, i, P}$ is the bias in the activity of a daSNC neuron corresponding to decision-dimension $i$ and pathway $P$. (activity arb. u.)
- $\mathrm{RMTg}$ is the output of RMTg, per eq. (5). (activity arb. u.)

In the current instance of the model, there are $q$ dsSPNs, $q$ isSPNs, $q$ dmSPNs, $q$ imSPNs, $q$ daSNC neurons that each receive projection from a dsSPN, and $q$ daSNC neurons that each receive projection from an isSPN. daSNC neurons corresponding to decision-dimension $i$ and pathway $P$ project to sSPNs and mSPNs of the same $i$ and $P$.

The sign (+ or -) of the $x_{i,P}(t)$ addition matches our interpretations of the experimental literature in **Table S2.** This aids in our examples in the analyses presented in **Figure 4**, where there is cortical data along reward-predominant and cost-predominant decision-dimensions. Note, though, that the direction of encoding is arbitrary and may be different for different decision-dimensions.

**Defining decision-space.**

A decision-dimension is used to form decision-space at times when daSNC activity corresponding to the decision-dimension exceeds a threshold:

(17) $$\mathrm{S}_{i,P}(t) = \begin{cases} 0 & y_{i,P}(t) < \text{threshold} \\ 1 & y_{i,P}(t) \geq \text{threshold} \end{cases}$$ (copied from **Figure S5** for convenience)

where:

- $y_{i,P}(t)$ is the activity of the daSNC neuron corresponding to decision-dimension $i$ and pathway $P$, as a function of time. (activity arb. u.)
- $\mathrm{S}_{i,P}(t)$ is the application of dopamine release (via daSNC activity) to mSPN activity corresponding to decision-dimension $i$ and pathway $P$. The notation here is used to match the notation in ***Instance 1***; $\mathrm{S}_{i,P}(t)$ is the $(i, i)$ element of the diagonal matrix $\mathbf{S}_P(t)$ whose elements correspond to the weights assigned to the decision-dimensions, similar to in eq. (3). (dimensionless)

**Defining action value.**

Action value is defined here like in ***Instance 1****,* except mSPN activity (for dmSPN and imSPN) is defined as a function of time:

(35) $$v_{j,P} = \frac{1}{1 + \exp\left( -\boldsymbol{\beta}_{j,P}\, \mathbf{s}_{\mathrm{mSPN},P}(t) - \alpha_{j,P} \right)}$$

where:

- $\boldsymbol{\beta}_{j,P}$ is a matrix of weights from dmSPNs to downstream action value encoding neurons for the direct pathway, or imSPNs to downstream inaction value encoding neurons for the indirect pathway. (dimensionless)

- $s_{sSPN,P}$ is the activity of sSPNs corresponding to decision-dimension $i$ and pathway $P$ (activity arb. u.)
- $t$ is time (seconds)
- $\alpha_{j,P}$ is an additive shift corresponding to the neuron encoding action $j$ for the direct pathway or inaction $j$ for the indirect pathway. (activity arb. u.)

As in the other instances of the model, there is one neuron encoding each $v_{j,P}$. So, there are $k$ neurons encoding action values and $k$ neurons encoding inaction values. Each of these neurons receives projection from all mSPNs of the corresponding pathway.

***Modeling time-variant input, using Instance 3.***

In **Figures 4C-J,** simulated responses of dsSPN, isSPN, dmSPN, and imSPN are plotted to various cortical inputs using the forward Euler method with step size 0.001s.

In **Figures 4C-F,** the cortical input to sSPN elements corresponding to each of the four-example direct pathway decision-dimensions is represented by a vector of length 5001 (corresponding to 0s to 5s with increments 0.001s). Four input vectors are used, one corresponding to each plotted decision-dimension. Each has a signal with Gaussian white noise added at every time increment:

(36) $2 + x_1,\ 1 + x_2,\ -1 + x_3,\ -1 + x_4,$ where $x_1, \ldots, x_4$ are Gaussian white noise processes

$x_1$ corresponds to a reward-predominant dimension (shown in green in **Figures 4C,D**). A relatively large positive mean value (2 arbitrary units) is assigned to it as an example of an important decision-dimension to a decision. A cost-predominant decision-dimension ($x_2$) is specified to be important, but less so, and hunger-predominant and location-predominant decision-dimensions ($x_3$ and $x_4$) are assigned to be relatively unimportant.

In **Figure 4E**, the number of decision-dimensions used to form decision-space is averaged across time steps in the 5s simulation. Simulations are run across 100 evenly spaced increments of $z_{daSNC,i}$ (for each pathway, depending on the simulation) in eq. (16) from -1 to 1 arbitrary units.

In **Figure 4F**, the activities of dmSPN or imSPN are calculated at each of the 5001 time-steps. Then mSPN activities along the four decision-dimensions included in the simulation are converted to values of an example action. This process uses the conversion from mSPN activities to action values in eq. (35). $\boldsymbol{\beta}_j$ is set to 1 and $\alpha_j$ to 0 for the example action $j$. Once action values have been calculated across all time-steps, an average is taken to assess the average action value over time. Simulations are run across 100 evenly spaced increments of $z_{\text{daSNC}, i}$ (for each pathway, depending on the simulation) in eq. (16) from -1 to 1 arbitrary units.

**Figures 4G,H** show examples of the response of dsSPN, isSPN, dmSPN, and imSPN elements to different cortical inputs. In **Figure 4G**, a cortical input of 10 arbitrary units for 2.5s is followed by an input of 20 arbitrary units for 2.5s. In **Figure 4H**, a cortical input of 10 arbitrary units for 2.5s is followed by an input of 0 for 2.5s.

In the analysis shown in **Figure 4I,** cortical inputs of 10 arbitrary units for 2.5s are followed by cortical inputs with prediction errors incremented by 0.1 from -1 to arbitrary units. These prediction errors are relative to the original cortical input of 10 arbitrary units. For example, for the prediction error of -1, there is a signal of 0 for 2.5s, and for the prediction error of 1, there is a signal of 20 for 2.5s. To find the change in the activities of circuit elements, their activities at 2.5s are subtracted from their activities at 5s.

**Figure 4J** shows simulations for an example input with $\kappa = 0$ in eq. (16) versus $\kappa = 0.1$ arbitrary units. The cortical input is as follows: in decision-dimension 1 (e.g. reward-predominant), a cortical signal of 10 arbitrary units for the 0-1.25s timeframe and elsewhere a signal of 0; in decision-dimension 2 (e.g. cost-predominant), a cortical signal of 10 arbitrary units for the 1.25-2.5s timeframe and elsewhere a signal of 0; in decision-dimension 3, a cortical signal of 10 arbitrary units for the 2.5-3.75s timeframe and elsewhere a signal of 0; and in decision-dimension 4, a cortical signal of 10 arbitrary units for the 3.75-5s timeframe and elsewhere a signal of 0.

For code, see https://github.com/dirkbeck/DM_space_model/blob/main/dynamic_model_and_neural_net/sSPN_DA_mSPN_dynamic_interaction.m.

Parameters are set to common values, chosen arbitrarily:

- in eqs. (13), (14): $\tau = 2$
- in eq. (13): $w_{\text{daSNC} \to \text{sSPN, direct}} = 1$
- in eq. (13): $w_{\text{daSNC} \to \text{sSPN, indirect}} = -1$
- in eq (14): $w_{\text{daSNC} \to \text{mSPN, direct}} = 1$
- in eq. (14): $w_{\text{daSNC} \to \text{mSPN, indirect}} = -1$
- in eq. (15): $\kappa = 0.1$
- in eq. (16): $\text{RMTg} = 0$
- in eq. (16): $z_{\text{daSNC}, i, P} = 0$ for all $i$ and $P$
- in eq. (17): $\text{threshold} = 0.5$

Additionally, the following initial conditions, also chosen arbitrarily, are used across analyses:

- $s_{\text{sSPN}, i, \text{direct}}(0) = s_{\text{sSPN}, i, \text{indirect}}(0) = s_{\text{mSPN}, i, \text{direct}}(0) = s_{\text{mSPN}, i, \text{indirect}}(0) = 0$
- $w_{\text{sSPN} \to \text{SNc}, i, \text{direct}}(0) = w_{\text{sSPN} \to \text{SNc}, i, \text{indirect}}(0) = 1$

***Movement of Circuit Activity Across Multiple Trials.***

Here, we model changes in circuit activity between trials. We begin by forming advantage and cost functions that guide the realignment of the circuit. Using these, we explore how vulnerability versus resilience in disorder formation could be interpreted through the lens of the model.

**Defining advantage and cost of circuit activity.**

*Advantage* is defined here as the ability of the circuit to produce beneficial decision-spaces at a certain activity. The goal of the sSPN-GPi-LHb-RMTg-daSNC circuit in the model is to produce preferred decision-spaces for action valuation (**Figures 7A-D**). For instance, in a laboratory environment when an animal routinely makes a choice to approach depending on reward level, a one-dimensional direct pathway decision-space with a reward dimension may be helpful. During a decision it may make sense for this animal to reach a circuit activity where forming a one-dimensional direct pathway decision-space is probable.

We represent this logic mathematically as a function of an $n$-element circuit $\left\{X_1, X_2, \ldots, X_n\right\}$. In our analysis, we focus on either FSI and sSPN, holding the rest of the circuit elements fixed at default values (see **Common parameters**); or sSPN, LHb, and daSNC, holding the rest of the circuit elements fixed at default values. The advantage of a certain circuit activity is defined as a weighted sum of probabilities the direct pathway decision-space occurs and the benefit of forming each decision-space:

(19) $$\text{advantage}(X_1=x_1, X_2=x_2, \ldots, X_n=x_n) = \sum_{l=1}^{2^q} \text{score}_l \cdot \text{P}(\text{space}_l | (X_1=x_1, X_2=x_2, \ldots, X_n=x_n))$$

(copied from **Figure S7** for convenience)

where:

- $\left\{X_1, X_2, \ldots, X_n\right\}$ are elements of the circuit with activities $x_1, x_2, \ldots, x_n$. (activity arb. u.)
- $q$ is the count of decision-dimensions
- $\text{score}_l$ is a coefficient corresponding to the preference for a given direct pathway decision-space. (dimensionless)

The probability each decision-space forms is derived from probability its decision-dimensions individually are used during the decision ($\text{daSNC}_i$ in eq. (2), here notated as $d_i$):

(37) $$\text{probability of formation of a decision-space } l = d_1\left(1-d_1\right)d_2\left(1-d_2\right)\cdot\ldots\cdot d_m\left(1-d_m\right)$$

The rationale for this formation of advantage is theoretical. Elsewhere, we show that direct pathway decision-spaces of different dimensionality are beneficial (and may be used by rodents) for tasks of different difficulties (**Figure 2**). We also show that certain decision-spaces are beneficial with certain levels of cortical noise (**Figure 5**) and for obtaining different types of action values (**Figures 6C-F**). We represent this as an assignment of greater value to certain decision-spaces, given external and internal contexts and the task at hand.

*Cost*, here, is defined as the difference between the circuit activity and a baseline activity. For most scenarios, the circuit might be best served searching for the circuit activity with the highest advantage. However, there is an obvious counterexample: it could be that it is easiest to form preferred decision-spaces at extremely unusual circuit activity (e.g. very high sSPN, very high LHb, very high daSNC), and only slightly more difficult to form that decision-space at closer to average circuit activity (average sSPN, low LHb, average daSNC). It may be more advantageous for the circuit to shift to the latter activity.

Therefore, we introduce a cost function to form *net advantage*. We then use net advantage to define the circuit activities that are the most beneficial. The concept of baseline circuit activity is introduced here in order to define cost. This can be interpreted as the circuit activity outside of decision-making.

(20) $$\text{cost}(X_1=x_1, X_2=x_2, \ldots, X_n=x_n) = \left\| \begin{bmatrix} x_1 & x_2 & \cdots & x_n \end{bmatrix}^{\mathsf{T}} - \begin{bmatrix} x_{1,\text{baseline}} & x_{2,\text{baseline}} & \cdots & x_{n,\text{baseline}} \end{bmatrix}^{\mathsf{T}} \right\|_2$$

(copied from **Figure S7** for convenience)

where:

- $\left\{X_1, X_2, \ldots, X_n\right\}$ are elements of the circuit with activities $x_1, x_2, \ldots, x_n$. (activity arb. u.)
- Outside of decision-making, $\left\{X_1, X_2, \ldots, X_n\right\}$ have baseline activities $x_{1,\text{baseline}}, x_{2,\text{baseline}}, \ldots, x_{n,\text{baseline}}$. (activity arb. u.)

Net advantage is defined as advantage minus cost multiplied by a constant:

(21) $$\text{net advantage}(X_1=x_1, \ldots, X_n=x_n) = \text{advantage}(X_1=x_1, \ldots, X_n=x_n) - \text{constant}\cdot\text{cost}(X_1=x_1, \ldots, X_n=x_n)$$

(copied from **Figure S7** for convenience)

where:

- Functions for reward and cost are taken from eqs. (19) and (20), respectively.
- $\text{constant}$ alters the weight given to cost compared to reward. (dimensionless)

**Visualizing advantage, cost, and net advantage.**

**Figure 7B** shows an example of how a circuit forms advantage per eq. (19). The advantage scores $\text{score}_i$ are randomly generated (normal distribution, mean 0, standard deviation 1) for each of the 16 possible decision-spaces formed from four decision-dimensions. FSI and sSPN activity are incremented across a 10x10 grid of sSPN and FSI activities which each range from 0 to 1 arbitrary unit. The activities of other circuit elements are set to default values in ***Instance 1*** of the model (see **Common parameters**).

Cost in **Figure 7C** is formed using eq. (20). Distance from the circuit baseline point is measured as Euclidean distance in the two plotted decision-dimensions.

Net advantage in **Figure 7D** is calculated per eq. (21). For this example, the cost coefficient $\text{constant} = 1$ (dimensionless coefficient).

**Figure 7F** similarly shows net advantage but after incrementing sSPN, LHb, and daSNC activities. Similar to in the example in **Figure 7D**, a set of scores are independently randomly generated and $\text{constant}$ in eq. (21) is set arbitrarily to 1 (dimensionless coefficient).

For code, see https://github.com/dirkbeck/DM_space_model/blob/main/circuit_trajectories/advantage_cost_net_advantage_example.m.

**Defining direction of circuit movement.**

The direction of movement is defined as a search for the most optimal circuit activity to produce advantageous direct pathway decision-spaces. To understand how a circuit governed by eq. (21) might adjust over the course of multiple trials, we relate the adjustment of circuit activity to net advantage. We specify that the circuit adjusts so that it can more easily reach advantageous decision-spaces. This constitutes a movement in the baseline activity of eq. (20).

Thus, the circuit adapts in the direction of the gradient of net advantage.

(22) $$\frac{\Delta\begin{bmatrix} x_{1,\text{baseline}} & x_{2,\text{baseline}} & \cdots & x_{n,\text{baseline}} \end{bmatrix}^{\mathrm{T}}}{\text{trial}} = \text{rate} \cdot \nabla \text{ net advantage}(X_1 = x_1, \ldots, X_n = x_n)$$

(copied from **Figure S7** for convenience)

where:

- $\{X_1, X_2, \ldots, X_n\}$ are elements of the circuit with activities $x_1, x_2, \ldots, x_n$. (activity arb. u.)
- Outside of decision-making, $\{X_1, X_2, \ldots, X_n\}$ have baseline activities $x_{1,\text{baseline}}, x_{2,\text{baseline}}, \ldots, x_{n,\text{baseline}}$. (activity arb. u.)
- $\text{rate}$ is a coefficient that affects the speed of movement of the circuit per trial. ($\text{trial}^{-1}$)

**Effect of initial circuit activity on future trials.**

In **Figures 7E,G, S7A**, several examples illustrate trajectories of circuit movement, as defined in eq. (21).

In these analyses, we assume that decision-dimensions are assigned equal importance by sSPN, i.e. (1) in eq. $b_{\text{sSPN}} = 0$. The probability that an individual decision-dimension is used to form decision-space is calculated using eq. (2). Advantage scores $\text{score}_i$ are randomly generated (normal distribution, mean = 0, standard deviation = 1 arbitrary unit) for each of the 16 possible decision-spaces created from four dimensions. The value of $\text{constant}$ in eq. (21) is set to 1 arbitrary unit. For each increment of a 15x15 grid of sSPN and FSI (**Figure 7E**) or 15x15x15 grid of sSPN, LHb, and daSNC (**Figures 7G, S7A**), net advantage is calculated for the scenario where that activity is the circuit baseline, i.e. $X_1 = x_{1,\text{baseline}}, X_2 = x_{2,\text{baseline}}, \ldots, X_n = x_{n,\text{baseline}}$. The gradient of this grid is approximated via MATLAB's gradient() routine. Trajectories are integrated from the gradient via MATLAB's streamline(), which uses the forward Euler method.

In the upper panel of **Figure 7E**, circuit baseline points are shown for each increment of Euler's method. The bottom panel shows the trajectories of 20 randomly selected points on the [0, 5] arbitrary units range for sSPN and FSI.

In **Figure 7G**, starting points are selected to form a 5 by 5 by 5 grid incremented along sSPN, LHb, and daSNC axes. At each point in a trajectory line, the probability a dimension is used to form decision-space is calculated via eq. (2). Then, during plotting, these calculated probabilities are interpolated using MATLAB's patch() routine.

In **Figure S7A**, the gradient is plotted using MATLAB's streamslice(). Then the trajectories of five randomly selected are plotted.

For code, see https://github.com/dirkbeck/DM_space_model/blob/main/circuit_trajectories/sSPN_DA_LH_trajectories.m and https://github.com/dirkbeck/DM_space_model/blob/main/circuit_trajectories/sSNC_DA_LH_trajectories_examples2.m

**Effect of altered advantage score on future trials.**

In **Figures S7B,C**, we examine the effect of a change to advantage $\text{score}_l$ in eq. (19) as a modeled task is being performed.

The circuit adjusts over 100 trials. At the beginning, advantage score is set to 1 (dimensionless coefficient) for the non-D decision-space and 0 for every other decision-space. Over the first 20 and the last 10 time-steps (between the dashed lines on the plots), advantage score is incremented for the 4D decision-space but not others. Over time steps 21-90, advantage score is incremented for the non-D decision-space.

In the plot, the value of each decision-space is divided by the total value of all decision-spaces to form a ratio.

For code, see https://github.com/dirkbeck/DM_space_model/blob/main/circuit_trajectories/changing_space_value_between_trials.m.

***Rationale for the computational framework.***

During the formation of our computational model, we considered several alternative modeling techniques, including neural networks, Hidden Markov Models, Bayesian methods, State Space Models, biophysics models, drift diffusion models, and others.

Here, we outline justifications for the modeling methods we chose and suggest other modeling techniques that may achieve similar results.

**Modeling choices: dimensionality reduction**.

We model the decision-making axes used to make choices (decision-dimensions) as the principal components of cortical data to link the processes of the circuit to operations commonly performed in data science. The decision-space model could easily be modified in a way that would preserve its concept, for example by replacing the principal components with the independent components[219] of cortical data. However, modifications may come at the cost of computational convenience or interpretability.

**Modeling choices: choice from physiology.**

We sought to derive choice and deliberation time of an arbitrary number of potential actions. Drift diffusion models have been successful in other work involving the Basal Ganglia[220], but classically only model two possible outcomes, although extensions have been made for the multi-outcome case. We designed our model to have a similar framework to drift diffusion models but be suited to an unlimited number of possible actions. Our model successfully reproduces choice in the T-maze task (**Figures S2B,C**). Our goal could also be achieved through models of interacting populations[221], though in our objective, we are more considered with choice outcomes (choice, deliberation time, deliberation time distribution) than neural recruitment to produce it. Alternatively, we might find success by treating choice as a Bayesian process arising from circuit activity[222] or through an intermediate layer such as in a Hidden Markov Model[223], though deliberation time is not commonly derived using these classes of models.

For code, see https://github.com/dirkbeck/DM_space_model/blob/main/Cross%20Correlation%20Pattern%20Counts/Neural%20Network/createHistogramsByTaskTypeOrConcentration.m (**Figure S2B**) and https://github.com/dirkbeck/DM_space_model/blob/main/Cross%20Correlation%20Pattern%20Counts/Neural%20Network/newDirkCode.m (**Figure S2C**)

**Modeling choices: physiological connections.**

We modeled the overall activities of circuit elements using relative firing rates because it achieved our goal of linking the overall activities of circuit brain regions. While models that model the timing of neuron spikes have been used to precisely describe connections in portions of the circuit[224,225], the decision-space conceptual model does not require their added granularity.

**Reasoning behind the FSI model.**

In eq. (1), scaling of cortical data serves the important functional role of normalizing $\mathbf{x}_P$ so that the circuit can make sense of data coded across the wide range of firing rates that are observed in the cortex[117] (**Figures 1D,E**). Previous anatomical work and physiological analysis has shown that many cortical neurons synapse to one FSI[117,226]. The cortical to FSI connection is excitatory[227]. This would suggest that FSI surveys the cumulative activity of cortex. Strengthening this hypothesis, physiological analysis of connected FSI and cortical neurons in the current work shows that FSI activity scales linearly with cortical activity[12,228] (see **Figures S3K,L**).

In the model, the parameter $a_{\mathrm{FSI}}$ is related to the strength of connection between cortex and FSI. The parameter $b_{\mathrm{FSI}}$ is related to firing rate of FSI when $\mathbf{x}_P = \mathbf{0}$.

For the FSI and SPN relationship, anatomical and physiological work has shown an inhibitory effect of FSI on SPN[229]. The algebraic operator that best represents this inhibition is less clear: it could be thought of as a subtraction or as a division depending on the experimental evidence used to model[230,231].

A subtraction possibility:

(38) $$\mathbf{s}_{\mathrm{sSPN}}\left(a_{\mathrm{FSI}}\right) = \mathbf{x}_P - a_{\mathrm{FSI}}$$

A division possibility:

(39) $$\mathbf{s}_{\mathrm{sSPN}}\left(a_{\mathrm{FSI}}\right) = \frac{\mathbf{x}_P}{a_{\mathrm{FSI}}}$$

where:

- $a_{\mathrm{FSI}}$ is the weight of cortex→FSI connection (dimensionless)
- $\mathbf{x}_P$ is the activities of the cortical neurons in a given pathway $P$ (activity arb. u.)

The analytic relationship used here leads to a convenient interpretation from the perspective of data processing. It can also be thought of as approximating subtraction or division depending on the scale of $a_{\mathrm{FSI}}$. Below, we show a series approximations of eq. (1) as a function of connection strength from cortex, with input from cortex and other parameters fixed.

Substitution in terms of $a_{\mathrm{FSI}}$:

(40) $$\mathbf{s}_{\mathrm{sSPN}}\left(a_{\mathrm{FSI}}\right) = \frac{1}{a_{\mathrm{FSI}} \cdot \left\|\mathbf{x}_P\right\|_2 + b_{\mathrm{FSI}}} \mathbf{x}_P \mathbf{W} + b_{\mathrm{sSPN}}$$

Truncated Taylor series at $a_{\mathrm{FSI}} = 0$:

(41) $$\mathbf{s}_{\mathrm{sSPN}}\left(a_{\mathrm{FSI}}\right) = \mathbf{b}_{\mathrm{sSPN}} + \frac{\mathbf{x}_P \mathbf{W}}{b_{\mathrm{FSI}}} - \frac{\left\|\mathbf{x}_P\right\|_2 \mathbf{x}_P \mathbf{W}}{b_{\mathrm{FSI}}^2} \cdot a_{\mathrm{FSI}} + \mathcal{O}\left(a_{\mathrm{FSI}}^2\right)$$

Truncated Laurent series at $a_{\mathrm{FSI}} = \infty$ :

$$s_{\mathrm{sSPN}}(a_{\mathrm{FSI}}) = b_{\mathrm{sSPN}} + \frac{\mathbf{x}_P \mathbf{W}}{\left\|\mathbf{x}_P\right\|_2} \cdot \frac{1}{a_{\mathrm{FSI}}} - \mathcal{O}\left(\frac{1}{a_{\mathrm{FSI}}{}^2}\right) \quad (42)$$

Thus, when FSI is weakly connected to cortex, the formula somewhat resembles a subtractive operation, and when FSI is strongly connected to cortex, the formula somewhat resembles a division operation. The ratio between the $a_{\mathrm{FSI}}$ and $b_{\mathrm{FSI}}$ parameters is important here (the first series, before truncation, converges when $\frac{|b_{\mathrm{FSI}}|}{a_{\mathrm{FSI}}} > \left\|\mathbf{x}_P\right\|_2$ and the second, before truncation, when $\frac{|b_{\mathrm{FSI}}|}{a_{\mathrm{FSI}}} < \left\|\mathbf{x}_P\right\|_2$).

***Inferring decision-space from SPN activity and choice.***

The decision-space can be inferred from environmental or experimental inputs in conjunction with decision-making data (**Figure S1Q,R**). The method here requires that many decision-making experiments have been run with a similar apparatus but different parameters (for instance, light level), and that average sSPN activity has been measured during the decisions. The parameter data is stored in a matrix $\mathbf{X} \in \mathbb{R}^{n \times p}$, where $n$ here is a separate trial with separate inputs (rather than different inputs across time steps, as in the description of ***Instance 1***) and $p$ is the number of features of the experiment that may be encoded by cortex, for instance temperature, music volume, or light level (similar to the description in ***Instance 1***).

The process involves three steps:

**Labeling sessions.**

Recorded decisions are labeled using attributes of the process by which the decision was made. This might be achieved in a rodent task that measures response to music volume, for instance, by clustering sessions based on heart rate and distance traveled.

**Constraints based on SPN activity.**

The sessions are split by label. $\mathbf{X}$ is standardized such that each column has a mean of 0 and standard deviation 1. Then a linear regression is run on each labeled subset $\mathbf{X}_l$ to find coefficients $\mathbf{b}_l$ that map those observations to predicted SPN activities (averaged across SPN decision-dimensions) $\hat{\mathbf{y}}_l$ for each session in the subset (here, the subscript $l$ is used to indicate all elements of the subset):

$$\hat{\mathbf{y}}_l = \mathbf{X}_l \mathbf{b}_l + b_0 \quad (43)$$

where:

- $\hat{\mathbf{y}}_l$ is predicted SPN activities
- $\mathbf{X}_l$ is a subset of experimental parameter data (see *Labeled sessions)* corresponding to one label
- $\mathbf{b}_l$ and $b_0$ are the linear regression coefficients that map the experimental parameters (see **Labeled sessions***)* to sSPN activity

From the calculated $\mathbf{b}_l$, we can guess the principal axis dimensions that make up decision-space $\mathbf{w}_1, \mathbf{w}_2, \ldots, \mathbf{w}_q$ and the presence of each decision-dimension $e_{li} \in \{0, 1\}$ in each labeled subset. During this process, we consider $\mathbf{b}_l$ as the sum of the dimensions in decision-space in the subset:

(44) $$\mathbf{b}_l = \sum_{i=1}^{q} e_{li} \mathbf{w}_i$$

Using this framework, we can form a rule which we can use to assign labels to decision-dimensions and hypothesize $\mathbf{w}_1, \mathbf{w}_2, \ldots, \mathbf{w}_q$:

> If $\mathbf{b}_A$ has high correlation with $\mathbf{b}_B$, then one of $\mathbf{b}_A$ or $\mathbf{b}_B$ must correspond to a higher-dimensional decision-space that also includes the dimensions of the other.

For example, we might have data with labels A, B, C, and D and corresponding $\mathbf{b}_A, \mathbf{b}_B, \mathbf{b}_C, \mathbf{b}_D$, where $\mathbf{b}_A$ is correlated with $\mathbf{b}_D$, $\mathbf{b}_B$ is correlated with $\mathbf{b}_D$, and the other possible pairs uncorrelated. It follows from the rule above that $\mathbf{b}_D$ is at least a two-dimensional decision-space including dimensions from $\mathbf{b}_A$ and $\mathbf{b}_B$, and that $\mathbf{b}_A$ and $\mathbf{b}_B$ are at least one-dimensional decision-spaces. So, we would hypothesize that there are two SPN-encoded decision-dimensions used during the decision, $\mathbf{w}_1$ and $\mathbf{w}_2$, and that subsets A and D use $\mathbf{w}_1$, subsets B and D use $\mathbf{w}_2$, and subset C does not use either.

The logic to construct constraints I and II are as follows. When $e_{l1} = e_{l2} = \ldots = e_{lq} = 0$ (i.e. a non-D decision-space), $\hat{\mathbf{y}}_l = b_0$. The coefficients assigned to the non-D decision-space $\mathbf{b}_{l:\ \text{0D space}}$ should have low correlation with any of the axes $\mathbf{w}_1, \mathbf{w}_2, \ldots, \mathbf{w}_q$. Therefore, it is expected that we find one $\mathbf{b}_l$ that has low correlation with the other $\mathbf{b}_l$, and we can label this $\mathbf{b}_{l:\ \text{0D space}}$. Further, when $e_{li}$ for one $i$ is equal to 1 and for all other $i$ is equal to 0 (i.e. a 1D decision-space), $\hat{\mathbf{y}}_l = \mathbf{w}_{i\ \text{in space}} + b_0$. This decision-space will have high correlation with $\mathbf{w}_{i\ \text{in space}}$ but low correlation with other $\mathbf{w}_i$.

Therefore, it is expected that we find several $\mathbf{b}_{l:\ \mathrm{dim}\ i\ \mathrm{space}}$ that are not correlated with one another, or $\mathbf{b}_{l:\ 0\mathrm{D\ space}}$, but may be related through the multi-dimensional decision-spaces in C. Finally, when $e_{li}$ is equal to 1 for multiple $i$, $\mathbf{b}_l$ is calculated as a sum per eq. (44). Therefore, it is expected that some $\mathbf{b}_{l:\ \mathrm{dim}\ i\ \cap\ \mathrm{dim}\ j\ \mathrm{space}}$ are linear combinations of $\mathbf{b}_{l:\ \mathrm{dim}\ i\ \mathrm{space}}$ and $\mathbf{b}_{l:\ \mathrm{dim}\ j\ \mathrm{space}}$.

The example shown in **Figure S1R** uses simulated data with 2 experimentally observed features (for instance, temperature and music volume) and 100 observations. $\mathbf{X}$ was constructed as a matrix of i.i.d. Gaussian variables with mean 0 and standard deviation 1. The ground-truth principal component matrix $\mathbf{W}$ was constructed as a 2x2 matrix of i.i.d. Gaussian variables with mean 0 and standard deviation 1. Then, for each of the 100 observations, a decision-space was randomly assigned in a reference dataset. Each decision-dimension for each observation was treated as an i.i.d. uniform variable, and if the variable corresponding to the observation and the decision-dimension exceeded a value (here, 0.5), then the dimension was considered as incorporated in decision-space. The randomly generated decision-spaces were each assigned their own label. Then a simulated SPN activity was created for each of the 100 observations was created by multiplying the ground-truth $\mathbf{W}$ by the decision-space used in each observation, similar to in eq. (1). i.i.d. Gaussian noise (mean 0, standard deviation 0.5) was then added to create simulated SPN activity observations. Using this simulated data, we ran a linear regression for each labeled subset via MATLAB's fitlm() routine. The linear regression fits are plotted as surfaces in **Figure S1R**. The slopes, with respect to temperature and music volume, were, for label A, -0.15 and 0.48, respectively; for label B, -437.90 and -9.72, respectively; for label C, -9.01 and -274.51 respectively; and for label D, -155.45 and -199.83. Slopes for label are relatively small, so it is assigned to "decision-space not formed" (matching the reference dataset). Of the remaining labels, label D is closer to an additive combination of labels B and C than other permutations, so label D is assigned the "2D decision-space" (matching the reference dataset), while labels B and C are each considered 1D decision-spaces (matching the reference dataset).

For code, see https://github.com/dirkbeck/DM_space_model/blob/main/model_overview/dimensionality_from_SPN_activity.m

**Tests using choice.**

We can then combine the constraints in Step 2, developed using SPN activity, with an analysis of choice given the hypothesized SPN-encoded decision-dimensions. In Step 2, a set of $\mathbf{w}_1, \mathbf{w}_2, \ldots, \mathbf{w}_q$ are hypothesized. Here, choice at different levels of those $\mathbf{w}_i$ are compared across the labeled subsets. The hypothesis in Step 2 is supported if choice, for each labeled subset, is correlated with the decision-dimensions hypothesized to be used to form decision-space but not correlated with the decision-dimensions hypothesized not to be used to form decision-space.

An example is shown in **Figure S1R.** For the case where decision-space is not formed, choices do not correlate with any hypothesized SPN-encoded decision-dimension. For a 1D decision-space, choices correlate with one hypothesized SPN-encoded decision-dimension. For a 2D decision-space, choices correlate with two hypothesized SPN-encoded decision-dimensions.

In the plotted examples, we plot simulated choices in each of the four decision-spaces used in the analysis. Choices are simulated from reference dataset (i.e. absent noise added during simulation) average sSPN activities by session $y$ by treating $Z = \frac{1}{1 + \exp(-y + \text{i.i.d. Gaussian noise})}$ as a random variable, where $Z < 0.5$ corresponds to a "turn left" action and $Z \geq 0.5$ to a "turn right" action. The threshold of 0.5 is chosen because it represents the expected value of $Z$ at average sSPN activity ($y = 0$). In the plots, we interpolate possible subject values from choice at different combinations of the two decision-dimensions $\mathbf{w}_1$ and $\mathbf{w}_2$, as derived in Step 2. For each action for each labeled subset, a logistic regression is used to convert actions to value of action across the grid. As expected, observations of label A, assigned to "decision-space not formed" have little correlation with either of the putative decision-dimensions derived in Step 2; observations of labels B and C, assigned to 1D decision-spaces, are correlated with one putative decision-dimension but not the other; and observations of label D, assigned to the "2D decision-space," are correlated with both.

For code, see
https://github.com/dirkbeck/DM_space_model/blob/main/model_overview/dimensionality_from_SPN_activity.m

***Testing the Model Through Analysis of Neural Data.***

As a further test of the decision-space model, we examined the relationships between behavior and neural activity in tasks that required different reward versus cost dimensions. The analysis, new to the current work, was performed using the Corticostriosomal Circuit Stress Experiment database (published with Friedman et al., 2017). We found more functionally connected sSPN and mSPN in tasks that were difficult. We then analyzed cortex, FSI, sSPN, and mSPN during these tasks and found evidence of dimensionality reduction from cortical neurons to SPNs.

**Defining decision difficulty by task.**

We defined decision difficulty through deliberation time, calculated as the time between when the door opened during the T-maze task and when the animal made a movement between one end of the maze or the other. Deliberation time distributions were analyzed for each trial group (for control and stress: cost-benefit cost, benefit-benefit, cost-cost, non-conflict cost-benefit). Skewness was calculated using MATLAB's skewness() routine. Example distributions are shown in **Figures S2B, S3A,B** (6 animals, 35 sessions) and summaries across groups in **Figure S2D, S3C,D** (14 rats, 249 sessions).

For code, see
https://github.com/dirkbeck/DM_space_model/blob/main/Cross%20Correlation%20Pattern%20Counts/Neural%20Network/createHistogramsByTaskTypeOrConcentration.m (**Figure S2B**) and
https://github.com/dirkbeck/DM_space_model/blob/main/Cross%20Correlation%20Pattern%20Counts/Pattern%20Analysis/createSkewnessBarChartOfTRAndCBAcrossControlAndStress2.m
(**Figure 2SD**)

**Replicating experimental deliberation time.**

To replicate the deliberation time distributions with a computational model, we leveraged a diffusion model where two functions would both start at zero and would continue towards either a positive or negative pre-defined threshold that represented either performing an action (positive threshold) or not doing an action (negative threshold). The first threshold that was reached by either function was which decision we considered as selected. The x-axis was the progression of time as the functions ran. The deliberation time for this model was the amount of time (in seconds) it took for the first function to reach its threshold. To match the experimental distribution times analyzed in **Figure S2B**, we ran the drift diffusion process adjusting drift rate, the thresholds, and noise until the modeled deliberation time distribution had a similar median and skew to experimental data from **Figure S2B** (https://doi.org/10.7910/DVN/SMKW0I). An additional 1.5 seconds was uniformly added to modeled distribution times.

For code, see https://github.com/dirkbeck/DM_space_model/blob/main/Cross%20Correlation%20Pattern%20Counts/Neural%20Network/createHistogramsByTaskTypeOrConcentration.m.

**Connected SPNs through cross-correlation.**

We used the Corticostriosomal Circuit Stress Experiment database to identify sSPN and mSPN among recorded neurons in the striatum. For details, see the Supplemental Materials & Methods of Friedman et al. (2017). 14785 cells across 14 control animals were analyzed. The total number of identified striosomal and matrix neuron per task: NCB = non-conflict cost-benefit (sSPNs = 14, mSPNs = 260), CC = cost-cost (sSPNs =46 , mSPNs = 400), CBC = cost-benefit conflict (sSPNs = 84, mSPNs =717), BB = benefit-benefit easy (sSPNs = 50, mSPNs =515 , chocolate milk concentration <50), BB = benefit-benefit difficult (sSPNs =33, mSPNs = 731, chocolate milk concentration >=50).

We binned the firing rates of sSPN and mSPN during the -3s to 3s window across all tasks into 1-5 bins. We used these firing rates to determine cross-correlation (MATLAB's xcorr()) between sSPN and mSPN recorded in the same session. Correlated pairs were defined as paired sSPN and mSPN that had a linear regression fit with correlation squared (MATLAB's corrcoef()) > 0.5 and significance p<0.04. To obtain the percentage of significantly correlated neurons, we counted the number of pairs that met the threshold for correlation and divided by the total number of identified pairs. Examples are plotted in **Figures S2E,F** and counts in **Figure S2G.**

To determine significance threshold for the counts of correlated pairs, we shuffled the sSPN and mSPN pairs across the database. We then performed the process above on the shuffled data. Using these shuffled pairs, we formed a distribution of correlations that might happen by chance. The threshold of significance was set to the 3 standard deviation mark among this distribution of shuffled pairs.

To produce **Figures S2E,F**, first follow the steps in run_me.m to generate 'pairsTableControl.mat', 'pairsTableStress.mat', and 'pairsTableStress2.mat' are first generated.

Then run the following lines of code in MATLAB:

```
examplePairedNeuronPLot('twdb_control', 'PLSvsFSI', 54); % Figure S2E
examplePairedNeuronPLot('twdb_stress', 'PLSvsFSI', 32); % Figure S2F
```

For the code that calls the figure creation in **Figure S2G**, see https://github.com/dirkbeck/DM_space_model/blob/main/Cross%20Correlation%20Pattern%20Counts/Fig7A_triplet-example/updated_cross_correlation.m

The fitting occurs in **Figure S2G**: https://github.com/dirkbeck/DM_space_model/blob/main/Cross%20Correlation%20Pattern%20Counts/Fig7A_triplet-example/matrix_strio_plot_dynamics.m

**Connected SPNs through Granger causality.**

sSPN and mSPN pairs were identified similarly to the previous section. 14785 cells across 14 control animals were analyzed and 25758 cells across 9 stress animals. The total identified number of striosomal and matrix cells for each of the tasks are the following: Control CC = cost-cost (sSPNs =46 , mSPNs = 400), Control BB = benefit-benefit (sSPNs =83 , mSPNs = 1246), Control CBC = cost-benefit conflict (sSPNs = 84, mSPNs =717), Stress CBC = cost-benefit conflict (sSPNs =41 , mSPNs =898 ), Stress BB = benefit-benefit (sSPNs = 156, mSPNs = 2813).

Then firing rates were determined over the span of the session by binning each trial into bins of 100ms. Granger causality (MATLAB's gctest()) was run on these sSPN and mSPN firing rate pairs to determine whether each pair was functionally connected. In control animals, 92 connected pairs were identified for the cost-benefit conflict task, 77 connected pairs for the cost-cost task, 1 connected pair for the non-conflict cost-benefit task, and 116 pairs for the benefit-benefit task. In stress animals, 110 connected pairs were identified for the cost-benefit conflict task and 292 connected pairs for the benefit-benefit task.

Next, for each of the trials with a functionally connected sSPN and mSPN pair, the class of motif was determined as either sSPN excited / mSPN excited, sSPN excited / mSPN inhibited, sSPN inhibited / mSPN excited, or sSPN inhibited / mSPN inhibited. These classes were formed by identifying whether, separately, the neurons were excited, inhibited, or neither in each of 5 blocks per each trial ([-15s, -3s], [-3s 0s], [0s 2.5s], [2.5s, 4.5s], [4.5s 20s]). A motif was counted if in a certain block each neuron was either excited or inhibited.

Excited or inhibited classifications were determined through inter-spike interval analysis, plotted in **Figures S2H-J**. The median inter-spike interval for each neuron across the full trial was calculated. A neuron was considered inhibited during the periods of the trial when inter-spike exceeded median. Conversely, a neuron was considered inhibited during periods when inter-spike interval fell below median. Excited blocks were defined as blocks where excitation time exceeded inhibition time inhibited by 10%. Inhibited blocks were defined, oppositely, as blocks where inhibition time exceeded excitation time by 10%. Blocks that reached neither threshold were not classified.

For code, see https://github.com/dirkbeck/DM_space_model/blob/main/Cross%20Correlation%20Pattern%20Counts/Pattern%20Analysis/plotBins.m.

Patterns were counted across the 5 bins and added across trials within the same group (e.g. control cost-benefit conflict). The total count, as plotted in **Figure 2F**, was divided by the number of trials to form an average number of functionally connected neurons across the 5 bins of the trial.

For code, see https://github.com/dirkbeck/DM_space_model/blob/main/Cross%20Correlation%20Pattern%20Counts/Pattern%20Analysis/allMaps/Maps%20By%20Task%20Type/createLinearPatternCountGraphsByTaskType.m.

Significance was determined similarly to the section above. Pairs of neurons, not necessarily functionally connected, were shuffled. From a shuffled distribution, the 3 standard deviation mark was identified and considered to be the threshold of significance.

For code, see https://github.com/dirkbeck/DM_space_model/blob/main/Cross%20Correlation%20Pattern%20Counts/Pattern%20Analysis/createRandomPatterns.m

**Analyzing neural dimensionality reduction.**

In the analysis plotted in **Figure 2G,** using data from the Corticostriosomal Circuit Stress Experiment database, we analyzed firing rates of 1) PL neurons that project to striatum (i.e. sSPN-projecting cortex, 221 sessions, 2-47 neurons per session), 2) FSIs (96 sessions, 2-12 neurons per session), 3) sSPNs (27 sessions, 2-9 neurons per session), and 4) mSPN (13 sessions, 2-6 neurons per session). For methods of classification see Friedman et al. (2017). Spike data was converted to firing rates by separating the spikes into 5-10 bins.

Covariance between the neurons was determined from the firing rates of simultaneously recorded neurons over time. For each trial, the effective correlation[138] was calculated from the activities of the $p$ neurons over time $\mathbf{X}$:

(45) $$\text{effective correlation} = 1 - \left\| \text{Corr}(X) \right\|^{1/p}$$

Effective correlations were then averaged across trials. Confidence intervals were determined from standard error between the sessions.

Code is in the directory https://github.com/dirkbeck/DM_space_model/tree/main/neuron_pair_analysis.

neuronSynchronyPlot.m generates the plot, which takes input files 'sessionDataOf....mat'.

These input mat files are generated by the 'covAndBinCtMatrixOfNeuronsFromSameSession.m' function which takes input files 'sameSession....mat'.

These input files are generated by 'extractNeuronsFromSameSession.m' function.

**Changes to choice after adding cost to a reward offer.**

In the analysis plotted in **Figure S3J,** we sought to define choice patterns before and after stress when a small cost was added to a reward offer. To do this, we analyzed choice data from the CBC task before and after stress (i.e. reward and a small cost) and in the BB task before after stress (i.e. only reward). We recorded the approach percentage across sessions and then averaged these between groups. Data from 17 rodents across 38 tasks is used for Control CBC, data from 13 rodents across 24 tasks for Stress CBC, data from 23 rodents across 114 tasks for Control BB, and data from 14 rodents across 116 tasks for Stress BB.

For data, see https://github.com/dirkbeck/DM_space_model/blob/main/disorder_hypotheses/experimental_data_analysis_choice_before_after_stress.xlsx.

**Analyzed cortex-FSI connectivity.**

Using data from the Corticostriosomal Circuit Stress Experiment database (published with Friedman et al. (2017)), firing rates and striatum-projecting prelimbic cortex neurons (i.e. sSPN-projecting cortex) and FSI were analyzed. For details regarding neuron classification, see Friedman et al. (2017). Spike data was converted to firing rates by separating data into 5-10 bins. A linear regression of the form $a \cdot x + b$ was fit through the firing rates. Examples are plotted in **Figures S3K,L** and averages of the square of Pearson correlation coefficient across neuron pairs and the $a$ slope parameter from the regression fit are plotted in **Figures S3O,P**. 78 neuron pairs across 7 rodents were analyzed before stress and 37 neuron pairs were analyzed across 4 rodents after stress.

Code is in the directory https://github.com/dirkbeck/DM_space_model/tree/main/neuron_pair_analysis.

To produce **Figures, S3K, L,** go to the directory https://github.com/dirkbeck/DM_space_model/tree/main/neuron_pair_analysis and run examplePairedNeuronPLot('twdb_control', 'PLSvsFSI', 54), examplePairedNeuronPLot('twdb_stress', 'PLSvsFSI', 32), respectively.

To produce **Figures S3O, P,** go to the directory https://github.com/dirkbeck/DM_space_model/tree/main/neuron_pair_analysis and run plotFitParamOfNeuronTriplets.m.

**Modeled cortex-FSI connectivity after stress.**

Connected cortical neurons and FSI show increased correlation and reduced slopes between their firing rates after stress (**Figures S3K,L,O,P**). To understand how this impacts SPNs, we modeled two changes to the cortex→FSI connectivity that might produce the experimental results (**Figures S3Q,R**). A first factor, connection weight $w_i$, measured the strength of connection between the $i$th connected cortical neuron and the FSI, that is, how strongly FSI would respond to an increase in activity in the connected cortical neuron. A second factor, number of connections $c$, measured the number of connected cortical neurons to the FSI.

We solved for correlation between the neurons and slope using identities that link covariance, correlation, and slope. This process assumes that FSI receives input only from the cortex.

(46) $$\text{correlation}(\text{cortex } 1,\ \text{FSI}) = \frac{\text{cov}(\text{cortex } 1,\ \text{FSI})}{\sqrt{\text{var}(\text{cortex } 1) \cdot \text{var}(\text{FSI})}}$$

(47) $$\text{slope}(\text{cortex } 1,\ \text{FSI}) = \frac{\text{cov}(\text{cortex } 1,\ \text{FSI})}{\text{var}(\text{FSI})}$$

(48) $$\text{cov}(\text{cortex } 1,\ \text{FSI}) = w_1 \text{var}(\text{cortex } 1) + \sum_{i=2}^{c} w_i \cdot \text{cov}(\text{cortex } 1,\ \text{cortex } i)$$

(49) $$\text{var}(\text{FSI}) = \sum_{i=1}^{c} w_i^2 \text{var}(\text{cortex } i) + \sum_{i=1}^{c} \sum_{\substack{j=1 \\ j \neq i}}^{c} w_i w_j \text{cov}(\text{cortex } i,\ \text{cortex } j)$$

For simplicity, we assumed in our analysis that variance is equal across cortical neurons projecting to the FSI (i.e. $\text{var}(\text{cortex } 1) = \text{var}(\text{cortex } 2) = \ldots = \text{var}(\text{cortex } c)$ ) and that the covariance between those neurons is equal (i.e. $\text{cov}(\text{cortex } 1, \text{cortex } i) = \text{cov}(\text{cortex } 1, \text{cortex } 2) = \text{cov}(\text{cortex } 1, \text{cortex } 3) = \ldots = \text{cov}(\text{cortex } 1, \text{cortex } c)$ ). So, in the analysis plotted in **Figures S3Q,R**, we substitute uniform variance and covariance to obtain:

(50) $$\text{correlation}(\text{cortex } 1,\ \text{FSI}) = \sqrt{\frac{\text{var}(\text{cortex } 1) + (c-1)\cdot \text{cov}(\text{cortex } 1, \text{cortex } i)}{c \cdot \text{var}(\text{cortex } 1)}}$$

(51) $$\text{slope}(\text{cortex } 1,\ \text{FSI}) = \frac{1}{cw}$$

where:

- $c$ is the number of cortex neurons connected to each FSI
- $w$ is the weight of each cortex→FSI connection

Thus, changes in strength of connection may be more likely to affect slope, while changes in the quantity of connections, the covariance between cortical neurons, and the variance of cortical neurons may also affect the cortex to FSI correlation. In stress, correlation and slope are both larger, suggesting reduction in quantity of cortex to FSI connections.

For code, see https://github.com/dirkbeck/DM_space_model/blob/main/disorder_hypotheses/ctx_to_FSI_synchrony_analysis.m.

**Simulation illustrating the theoretical change.**

We then conducted a simulation to examine the case where cortical data and cortex to FSI connectivities were less uniform. To do this, we constructed a network with random unit normal connection weights from cortex to FSI. Then we randomly lesioned all but several of the connections to reflect the sparsity of the brain. The connection from the first cortical neuron to the FSI was always preserved.

In each of ten simulations, a cortical input to sSPN was generated as a random unit normal 100x1 vector. Activity of the first cortical neuron and the FSI was recorded. These are plotted in **Figures S3M,N** along with a linear regression fit.

For code, see https://github.com/dirkbeck/DM_space_model/blob/main/disorder_hypotheses/ctx_to_FSI_synchrony_analysis.m.

**KEY RESOURCES TABLE**

| REAGENT or RESOURCE | SOURCE | IDENTIFIER |
|---|---|---|
| Deposited data | | |
| Corticostriosomal Circuit Stress Experiment | Friedman et al. (2017) | https://data.mendeley.com/datasets/z9jd8xhj84/1 |
| Computational model of striosomal circuit in normal and disordered decision-making | This paper | https://doi.org/10.7910/DVN/SMKW0I |
| Overview of the model | This paper | https://github.com/dirkbeck/DM_space_model/tree/main/model_overview |
| Tests of the model | This paper | https://github.com/dirkbeck/DM_space_model/tree/main/model_tests |
| Disorder hypotheses | This paper | https://github.com/dirkbeck/DM_space_model/tree/main/disorder_hypotheses |
| Instances 2 (sparse connectivity) and 3 (dynamics) | This paper | https://github.com/dirkbeck/DM_space_model/tree/main/dynamic_model_and_neural_net |
| Day to day differences and disorder comorbidity | This paper | https://github.com/dirkbeck/DM_space_model/tree/main/day_to_day_space_sampling |
| Circuit adjustment between trials | This paper | https://github.com/dirkbeck/DM_space_model/tree/main/circuit_trajectories |
| Analysis of the correlation between cortical neurons, FSIs, sSPNs, and mSPNs during decision-making | This paper | https://github.com/dirkbeck/DM_space_ |

| | | |
|---|---|---|
| | | model/tree/main/neuron_pair_analysis |
| Analysis of the functional connectivity of sSPNs and mSPNs during decision-making | This paper | https://github.com/dirkbeck/DM_space_model/tree/main/Cross%20Correlation%20Pattern%20Counts |
| Software and algorithms | | |
| MATLAB R2021a | Mathworks | https://www.mathworks.com/products/matlab.html |
